# Synthetic Control Enables Reliable Cluster Validation and Marker Discovery in Omics Data: From Single-Cell and Spatial to Population-Scale

**DOI:** 10.1101/2023.07.21.550107

**Authors:** Dongyuan Song, Siqi Chen, Christy Lee, Changhu Wang, Pan Liu, Yiwen Yang, Kexin Li, Yihui Cen, Guanao Yan, Qingyang Wang, Xinzhou Ge, Weijian Wang, Tanye Wen, Daniel Shams, Kris Sankaran, Nikos Konstantinides, Wei Li, Jingyi Jessica Li

**Affiliations:** Department of Genetics and Genome Sciences, University of Connecticut Health Center, Farmington, CT, USA 06030; Interdepartmental Program of Bioinformatics, University of California, Los Angeles, CA, USA 90095-7246; Department of Statistics and Data Science, University of California, Los Angeles, CA, USA 90095-1554; Department of Statistics, Oregon State University, Corvallis, OR, USA 97331-4606; Department of Computational Mathematics, Science and Engineering, Michigan State University, East Lansing, MI, USA 48824; Department of Computational Medicine, University of California, Los Angeles, CA, USA 90095-1766; Department of Biomedical Engineering, University of California-Irvine, Irvine, CA, USA 92697; Department of Statistics, University of Wisconsin-Madison, WI, USA 53703; Division of Computational Biomedicine, Department of Biological Chemistry, University of California, Irvine, Irvine, CA, USA 92697; Université Paris Cité, CNRS, Institut Jacques Monod, F-75013 Paris, France; Biostatistics Program, Public Health Sciences Division, Fred Hutchinson Cancer Center, Seattle, WA, USA 98109; Department of Biostatistics, University of Washington, Seattle, WA, USA 98195-1617

**Keywords:** Clustering-induced bias, Post-clustering inference, Synthetic null controls, Marker discovery, False discovery rate control, Single-cell RNA-seq, Spatial transcriptomics, Bulk transcriptomics, Multi-omics, Microbiome

## Abstract

Post-clustering differential analysis is widely used across omics, including single-cell and spatial transcriptomics, multi-omics, population-scale bulk transcriptomics, and microbiome data. After a clustering algorithm finds clusters based on omics features as putative cell types, spatial domains, or subpopulations, statistical tests are typically applied to the same data to identify differential features as potential markers. Because the features that drive clustering are inherently more likely to appear differential, this clustering-induced bias can yield false-positive markers and mislead the interpretation of clusters as meaningful biological entities, especially when clusters are spurious, resulting in ambiguously defined cell types, spatial domains, or subpopulations. This dual challenge—determining whether clusters themselves are reliable and, if so, identifying their true markers—requires a unified statistical solution. To address this challenge, we propose ClusterDE, a statistical method designed to identify post-clustering differential features (with differentially expressed (DE) genes as a primary example) as reliable markers of cell types, spatial domains, or subpopulations while controlling the false discovery rate (FDR), regardless of clustering quality. The core of ClusterDE involves generating synthetic null data as an *in silico* negative control representing a single homogeneous group (such as a cell type, spatial domain, or population), allowing for the detection and removal of spurious markers caused by clustering-induced bias. In single-cell and spatial transcriptomics analyses, ClusterDE controls the FDR, prioritizes canonical cell-type and spatial-domain markers among the top discoveries, and deprioritizes housekeeping genes, and it can refine underlying cell-type hierarchies by merging spurious or over-clustered groups. ClusterDE also mitigates spurious bifurcating trajectories in single-cell analyses that arise from over-clustering and retrospectively evaluates contested cell-type claims in high-profile studies, including a retracted human fetal cerebellum atlas and a challenged COVID-19 developing-neutrophil trajectory. At atlas scale, applying ClusterDE to the Allen Human Middle Temporal Gyrus taxonomy merges fine-grained clusters that are not statistically supported by single-cell transcriptomics, while preserving transcriptomically distinct and spatially supported cell types. Moreover, built-in synthetic null quality checks allow users to experiment with multiple state-of-the-art simulators and retain those that pass the diagnostics for their specific datasets. ClusterDE is compatible with widely used analysis pipelines such as Seurat and Scanpy and supports flexible, data-specific synthetic null generation, enhancing post-clustering inference across diverse omics modalities, with demonstrated efficacy in population-scale bulk transcriptomics and microbiome analyses.

## 1 Introduction

Single-cell RNA-seq (scRNA-seq) and spatially resolved transcriptomics (SRT) technologies have revolutionized transcriptomic studies by providing unprecedented snapshots of gene expression in individual cells and within their spatial contexts. A major task in scRNA-seq and SRT data analysis is to annotate cell types and spatial domains using marker genes. For example, in scRNA-seq data analysis, a standard annotation procedure includes two steps: (1) partitioning cells into clusters, and (2) finding differentially expressed (DE) genes between cell clusters as marker genes for annotating clusters as potential cell types [1, 2, 3, 4], as in state-of-the-art analysis pipelines such as the R package Seurat [5] and the Python module Scanpy [6]. However, it has been realized that this post-clustering differential expression (DE) procedure is conceptually flawed. For instance, Seurat contains the warning message that “*P* values should be interpreted cautiously, as the genes used for clustering are the same genes tested for differential expression.” This issue is commonly referred to as “double dipping,” meaning that the same gene expression data are used twice: first to find cell clusters and then to identify DE genes. The double dipping issue leads to an inflated false discovery rate (FDR) when identifying post-clustering DE genes as putative cell-type marker genes. Particularly, the FDR inflation is more pronounced when the cell clusters are more ambiguous. This raises two closely related but distinct questions: whether the clusters themselves are reliable, and, if so, which genes reliably distinguish them—a dual problem that motivates the method we introduce below. To explain this double-dipping issue in scRNA-seq cell-type annotation, we discuss two extreme scenarios. In one scenario, if two cell types are considered “distinct” (i.e., cell-type marker genes exhibit bimodal expression distributions, with the high-expression mode corresponding to the cell type in which they are highly expressed) and the inferred clusters perfectly match the cell types, then double-dipping has no impact on post-clustering DE analysis. In this case, DE analysis can successfully identify cell-type marker genes that are highly expressed in a cluster corresponding to a distinct cell type (Fig. 1a, top). In the opposite scenario, when a clustering algorithm over-clusters a single cell type into two clusters, post-clustering DE analysis will identify genes highly expressed in each cluster due to double-dipping: the clustering is based on gene expression data, and variations in the expression of certain genes drive the clustering, resulting in these genes being highly expressed in one of the clusters (Fig. 1a, bottom). However, since only one true cell type exists, these post-clustering DE genes cannot be used to justify the two clusters as distinct cell types and should not be regarded as cell-type marker genes.

**Fig. 1:**
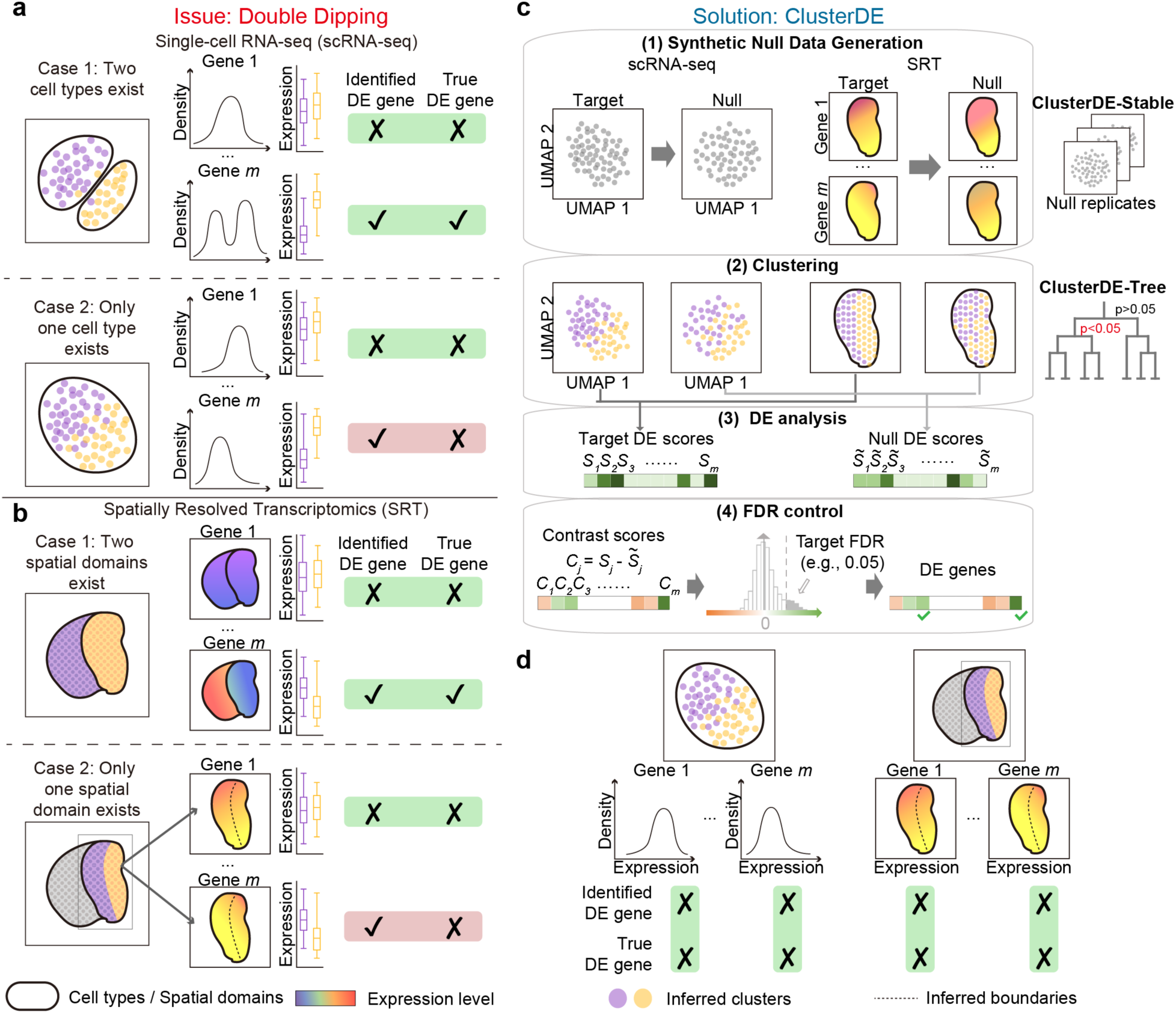
ClusterDE mitigates double dipping in post-clustering DE analysis for identifying cell-type and spatial-domain marker genes. **a-b**, An illustration of the double-dipping issue in (**a**) scRNA-seq and (**b**) SRT post-clustering DE analysis. In case 1, where two cell types or spatial domains exist and are accurately identified through clustering, double dipping has minimal effect, and the identified post-clustering DE genes align well with the true cell-type or spatial-domain marker genes (top row in panels **a** and **b**). However, in case 2, where a single cell type or spatial domain is incorrectly split into two clusters, double dipping results in post-clustering DE genes that are false-positive cell-type or spatial-domain marker genes (bottom row in panels **a** and **b**; e.g., gene *m* is identified as a false-positive marker gene). **c**, An overview of ClusterDE, consisting of four steps. Step (1): Given the target real data (“Target”), ClusterDE generates synthetic null data (“Null”) to represent the null hypothesis of a single “hypothetical” cell type (scRNA-seq) or spatial domain (SRT). Steps (2)–(3): A clustering algorithm is used to divide cells or spatial spots into two clusters, followed by a DE test to compare each gene between the clusters. This process is applied to both the target data and the synthetic null data, producing two DE scores per gene—one from each dataset. Step (4): ClusterDE calculates a contrast score for each gene by subtracting the synthetic null DE score from the target DE score. It then applies the FDR-control method Clipper to determine a contrast score cutoff based on all genes’ contrast scores and a target FDR (default: 0.05). DE genes are identified as those with contrast scores exceeding this cutoff, as indicated by the vertical dashed line in the contrast score distribution across genes. In addition to its core algorithm, ClusterDE includes two extensions: ClusterDE-Stable, which improves reproducibility by aggregating results across multiple synthetic null replicates, and ClusterDE-Tree, which supports hierarchical and multi-cluster marker identification and provides cluster-level split tests within a cluster tree. **d**, ClusterDE mitigates the false discoveries caused by double dipping in case 2, shown in **a** and **b** (e.g., gene *m* is no longer identified as a false-positive marker gene).

Some biologists might argue that even if the two clusters are “indistinct” (i.e., all genes exhibit unimodal expression distributions across all cells), they could still reflect meaningful biological variation and thus represent potential cell sub-types. While this argument may hold in certain scenarios, we argue that if the two clusters are indistinct, it is impossible to determine whether they should be classified as two distinct cell types or merely arbitrary divisions along a continuum of cell state changes, without external knowledge. A more critical issue is that if clusters are not required to be distinct, cells could be continuously divided into finer and finer clusters, which would make it unreasonable to consider any such clusters as potential cell types. This would also hinder transferring cell-type labels across datasets, a key barrier to translating single-cell findings into clinical practice [7]. Given this issue and the need for a clear definition of cell types in cell atlases, as reflected by the ongoing debate about what constitutes a cell type [8, 9], we focus on the practice of using post-clustering DE analysis to annotate distinct cell clusters as potential cell types. Notably, several prominent cell-type claims in high-profile studies have later been challenged as arising from precisely this kind of ambiguous, indistinct clustering (see “Results”), underscoring the practical stakes of this problem. Methods aimed at addressing the double-dipping issue in scRNA-seq post-clustering DE analysis include the Truncated Normal (TN) test [10] and the Countsplit method [11]. The TN test method relies on splitting cells and consists of five steps: (1) dividing the cells into two sets—training and test cells; (2) applying a clustering algorithm to the training cells to identify two clusters; (3) training a support vector machine (SVM) classifier on the training cells to predict cluster labels based on gene expression; (4) using the trained classifier to predict the cluster labels of the test cells; and (5) identifying DE genes between the two clusters in the test cells using the TN test. Since the TN test method requires gene expression levels to follow normal distributions, it applies the log-transformation to the scRNA-seq count data before the five steps. Instead of splitting cells, the Countsplit method splits the scRNA-seq cell-by-gene count matrix into training and test matrices of the same dimensions (cells and genes) using a procedure called data thinning [12], which splits each count into two counts, whose sum equals the original count, through binomial sampling. Countsplit identifies cell clusters by applying a clustering algorithm to the training matrix. Since the test and training matrices contain the same cells, the cell cluster assignments can be directly applied to the test matrix. Countsplit then identifies DE genes between the clusters based on the test matrix, allowing the use of any suitable statistical test for count data, beyond the TN test. Although both methods claimed to provide well-calibrated *P* values (i.e., uniformly distributed between 0 and 1 under the null hypotheses), one per gene, our findings indicate that their *P* values are anti-conservative (i.e., enriched near 0) in the presence of gene-gene correlations (see “Results”). The reason behind this issue is that the validity check of *P* values in the TN test and Countsplit papers relied on simulation studies that implicitly assumed genes to be independent [11], an assumption that, however, does not hold in real scRNA-seq data. As a result of their anti-conservative *P* values, both methods report numerous false discoveries on real single-cell RNA-seq data from a single cell line (see “Results”).

Several cluster-free DE tests attempt to circumvent the double-dipping issue by bypassing the cell clustering step [13, 14, 15, 16, 17, 18]. However, it is important to note that these cluster-free methods are not designed to identify potential cell types. Consequently, the DE genes identified by these methods cannot be interpreted as marker genes for specific cell types, unlike the DE genes identified post-clustering. In other words, cluster-free DE genes and post-clustering DE genes serve different purposes and are not conceptually comparable. Another class of methods aims to assess the quality of clustering results, such as evaluating the “purity” of a cluster or determining if two clusters should be merged [19, 20, 21, 22, 23, 24, 25, 26]. However, these methods lack a direct statistical test for identifying DE genes and rely on setting a threshold for clustering quality—an arbitrary decision to mitigate concerns about double dipping. Importantly, we find that even when cell types are truly distinct, imperfect clustering (e.g., mixing a fraction of cells) can still bias post-clustering DE, elevating highly variable genes that drive clustering but are not markers of any cell type (e.g., housekeeping genes; see “Results”). In this study, we focus on identifying reliable marker genes for cell clusters during the cell-type annotation process, as well as on determining whether the clusters themselves warrant retention or merging. Hence, we do not consider cluster-free DE tests and methods that only assess clustering quality as competing alternatives in our investigation.

The post-clustering DE bias issue is not unique to scRNA-seq data; it is also relevant for more recent SRT data, which captures tissue context information of cells or cell neighborhoods, depending on the spatial resolution of the SRT technology used [27]. A common analysis task is spatial domain detection, which involves clustering proximal spatial spots (representing cells or cell neighborhoods) based on their gene expression profiles, with the goal of identifying spatial domains that correspond to functionally distinct tissue structures [28]. Although not widely discussed, the double-dipping issue can lead to unreliable marker genes for “indistinct” spatial domains, where no genes show sharp expression changes at the identified domain boundaries [29]. However, we argue that reliable spatial domains should be “distinct” from nearby domains, meaning that the domain’s marker genes should exhibit sharp expression changes at the boundaries. Otherwise, similar to the challenge of defining cell types in scRNA-seq data, indistinct spatial domains cannot have their labels reliably transferred across datasets. Despite this challenge, no methods have been developed to circumvent the double-dipping issue in spatial domain annotation. Motivated by the need for annotating distinct spatial domains, we define spatial domain marker genes as those exhibiting discontinuity or sharp changes in expression at domain boundaries (Fig. 1b, top). In contrast, genes that are not defined as spatial domain markers may still show spatial variation, but the variation is smooth (Fig. 1b, bottom). Similar to scRNA-seq data analysis, our goal is to enable post-clustering DE analysis to identify reliable marker genes for annotating distinct spatial domains. Beyond scRNA-seq and SRT, clustering followed by differential analysis is common across omics. For example, in population-scale bulk transcriptomics [30, 31, 32], clustering individuals by transcriptome profiles and then performing DE between clusters can spuriously “validate” over-clustered groups as distinct subpopulations. In single-cell multi-omics, clustering may be performed using RNA-seq while differential accessibility (DA) is assessed on ATAC-seq [33, 34, 35]. Although such cross-modality analyses avoid literal double dipping, they remain prone to inflated false positives if clustering does not reflect distinct cell types or subtypes. Notably, existing methods such as the TN test and Countsplit are inapplicable here because they require clustering and DE to use the same features within a single modality.

Here we introduce ClusterDE, a synthetic-control framework designed to serve two related goals: (i) determining whether putative clusters are statistically supported or should be merged, and (ii) identifying reliable marker genes for distinct cell types, spatial domains, or subpopulations while avoiding the inflated FDR caused by clustering-induced bias. ClusterDE effectively controls the FDR when identifying marker genes, even in cases where clusters are spurious. The ultimate goal is to use the identified marker genes to reliably annotate distinct cell types, spatial domains, or subpopulations. It is important to note that ClusterDE is not intended to replace existing pipelines that perform clustering followed by DE (e.g., Seurat or Scanpy for scRNA-seq); rather, it serves as an add-on that yields more reliable discoveries. A practical advantage of ClusterDE is that it makes an abstract statistical null hypothesis concrete by generating synthetic null data (i.e., *in silico* negative controls). This allows users to assess whether the synthetic null data reflects the negative-control scenario they envision and, if needed, to select among multiple state-of-the-art simulators and verify, via built-in quality checks, that the chosen simulator generates reliable synthetic null data. ClusterDE demonstrates strong performance across omics. In scRNA-seq, ClusterDE was benchmarked against Seurat (which includes double dipping), the TN test, and Countsplit, and was the only method to effectively control the FDR and eliminate false discoveries in five cell-line datasets. It deprioritizes unlikely marker genes (e.g., housekeeping genes) and prioritizes canonical markers in peripheral blood mononuclear cell (PBMC) datasets, and it identifies spurious cell clusters and improves cell-type separation in an adult *Drosophila* neuronal visual system dataset. Moreover, ClusterDE identifies cell-type markers that respect an underlying cell-type hierarchy and mitigates spurious bifurcating trajectories arising from over-clustering. At atlas scale, applying ClusterDE-Tree to the Allen Human Middle Temporal Gyrus taxonomy refines over-resolved cell-type clusters while preserving transcriptomically distinct and spatially supported cell types, as validated against an independent SRT dataset. ClusterDE also retrospectively flags cell-type claims from high-profile studies that were later disputed by independent reanalyses, including a retracted human fetal cerebellum atlas and a challenged COVID-19 developing-neutrophil trajectory. In SRT, ClusterDE outperforms a double-dipping workflow (BayesSpace for clustering + Seurat for DE analysis), eliminating false-positive DE genes between spurious spatial clusters in human brain and human pancreatic cancer tissue and yielding domain markers that better align with annotated regions. Beyond scRNA-seq and SRT, ClusterDE generalizes to single-cell multi-omics (RNA/ATAC), population-scale bulk transcriptomics (microarray and RNA-seq), and microbiome analyses: it removes spurious discoveries when clusters are not distinct and yields more specific, interpretable markers when they are, thanks to flexible, data-specific synthetic null generation.

Taken together, these findings highlight that ClusterDE addresses a general form of post-clustering inference bias that extends beyond literal “double dipping.” Even when clustering and differential testing rely on different features or modalities, inaccurate clustering or over-clustering can still induce spurious differential signals. By constructing data-specific synthetic negative controls, with built-in diagnostics to ensure their adequacy, ClusterDE distinguishes true biological differences from clustering-induced artifacts across diverse omics settings, positioning it as a broadly applicable framework for reliable post-clustering cluster and marker inference.

## 2 Results

### 2.1 ClusterDE: a synthetic control and contrastive approach for reliable cluster and marker discovery

ClusterDE introduces a novel synthetic control and contrastive approach for identifying reliable differential features that are robust to clustering-induced bias, a general problem for which double dipping is a special case. It applies to diverse omics modalities where clustering is followed by differential analysis, including scRNA-seq, SRT, single-cell multi-omics, bulk transcriptomics, and microbiome data. The approach centers on establishing an *in silico* negative-control dataset (referred to as the “synthetic null data”) that is analyzed in parallel with the real dataset (referred to as the “target data”) through the same computational pipeline, which includes clustering followed by differential analysis. The contrastive approach then identifies reliable cluster markers by comparing differential analysis results from the target data to those from the synthetic null data. The same synthetic null data idea also provides a complementary cluster-level test: by generating multiple synthetic null replicates and comparing a cluster-separation statistic computed on the target data to its empirical distribution under the null, ClusterDE determines whether two clusters are statistically distinct or should be merged, addressing the dual goals of validating clusters and identifying their markers (see “Supplementary Methods” and later subsections for details).

For illustration, we describe ClusterDE in the contexts of scRNA-seq and SRT, where the target data consists of cells or spatial spots divided into two potentially ambiguous clusters, which are then analyzed by DE analysis to identify cell-type or spatial-domain marker genes. To generate the synthetic null data, ClusterDE provides two null models: an scRNA-seq null model, which assumes that all cells belong to a homogeneous cell type, with gene expression exhibiting unimodal distributions across the cells; and an SRT null model, which assumes that all spatial spots belong to a homogeneous domain, with gene expression varying smoothly within the domain. Under these null models, no cell-type or spatial-domain marker genes are expected to be detected. To identify potential cell-type or spatial-domain marker genes, ClusterDE involves four main steps (Fig. 1c).

Step 1 of ClusterDE is synthetic null data generation, where a statistical simulator is used to generate synthetic null data representing a hypothetical homogeneous cell type or spatial domain. This step is simulator-agnostic by design: ClusterDE’s synthetic control serves as a calibration device for inference, and any simulator capable of generating a homogeneous, single-population dataset that preserves the target data’s key statistics—gene means, variances, and, importantly, gene-gene rank correlations—while maintaining the same number of cells or spatial spots and the same set of genes, can in principle be used. To ensure this holds regardless of which simulator a user selects, ClusterDE performs a set of built-in quality checks on the synthetic null data to verify that key statistical properties of the target data are preserved and that no residual cluster structure remains (see “Results” and “Supplementary Methods”). For scRNA-seq and SRT data, ClusterDE uses the statistical simulator scDesign3 [36] by default, based on a systematic evaluation of several representative alternatives (see “Results”) showing that scDesign3 most reliably satisfies these requirements for these two modalities (Figs. S2 for scRNA-seq; Figs. S3 and S4 for SRT). This component is pluggable, and for other modalities—such as single-cell multi-omics (RNA/ATAC), bulk transcriptomics, and microbiome data—ClusterDE instead uses modality-specific null generators better suited to their data characteristics (see “Supplementary Methods” for details). Users may also replace scDesign3 with an alternative null generator for scRNA-seq or SRT data, for instance to accelerate null generation. Steps 2 and 3 of ClusterDE comprise a user-defined pipeline for clustering and subsequent DE analysis (e.g., Seurat in R or Scanpy in Python for scRNA-seq). ClusterDE is modular: users may choose any clustering algorithm and gene-level DE test appropriate for their data type and analysis goals, provided the DE test returns a per-gene score (e.g., a test statistic or a monotone transform of a *P* value). The pipeline is applied in parallel to the target and synthetic null data. When used with Seurat, ClusterDE naturally supports Seurat’s built-in DE tests (demonstrated in the next subsection). These two steps yield a “target DE score” and a “null DE score” for each gene.

Step 4 of ClusterDE implements a contrastive strategy: for each gene, ClusterDE computes a “contrast score” by subtracting its null DE score from its target DE score, and a gene is identified as a reliable cell-type or spatial-domain marker only if its contrast score is significantly greater than zero. True non-marker genes are expected to have contrast scores that are symmetric about zero or, more generally, at most left-skewed. ClusterDE uses the FDR control method Clipper [37] to determine a contrast score cutoff based on a target FDR (e.g., 0.05). Genes with contrast scores at or above the cutoff are identified as DE genes. For a formal proof that ClusterDE controls the FDR with a large number of genes, see “Theoretical justification of ClusterDE” in the Supplementary Material.

Through these four steps, ClusterDE effectively eliminates false-positive marker genes caused by clustering-induced bias, particularly when the target data consists of a single cell type or spatial domain (Fig. 1d). In addition to this core framework, referred to as ClusterDE-Core, ClusterDE includes two extensions: ClusterDE-Stable, which enhances reproducibility by aggregating results across multiple synthetic null replicates, and ClusterDE-Tree, which enables hierarchical and multi-cluster marker identification, together with principled merging of statistically unsupported splits, particularly useful for cell-type annotation in scRNA-seq data (Fig. 1c). Both extensions are described and evaluated in later subsections.

ClusterDE is designed to integrate seamlessly with existing analysis workflows while providing tools that promote rigor and specificity in post-clustering marker identification. To improve scalability, ClusterDE-Fast accelerates synthetic null generation, enabling efficient use of ClusterDE-Stable even on large-scale datasets. For clustering-based trajectory inference, our trajectory extension adapts ClusterDE to bifurcating lineage structures to correct over-clustering-induced bias in lineage marker identification. These components together make ClusterDE a practical, robust, and interpretable tool for post-clustering differential analysis across diverse omics modalities, spanning single-cell, spatial, and population-scale data.

### 2.2 Simulations verify ClusterDE’s reliable FDR control and statistical power in scRNA-seq data

We conducted extensive simulation studies on scRNA-seq data to validate ClusterDE as a post-clustering DE method with reliable FDR control under double dipping. (Results on simulated SRT data are presented in Section “ClusterDE reveals reliable spatial-domain marker genes.”) Unless otherwise noted, ClusterDE below refers to ClusterDE-Core, which uses a single synthetic null replicate; the stability-enhanced extension ClusterDE-Stable is introduced and evaluated later in this subsection. We also compared ClusterDE with Seurat, the most widely used analysis pipeline that involves double dipping, and two methods designed to address double dipping: the TN test [10] and Countsplit [11]. For all four methods, we used the default Louvain clustering algorithm in Seurat, as it is commonly employed in scRNA-seq data analyses. In the DE analysis step for ClusterDE, Seurat, and Countsplit, we evaluated five DE tests available in Seurat: the Wilcoxon rank-sum test (Wilcoxon; the default in Seurat), the two-sample *t*-test (t-test), the negative binomial generalized linear model (NB-GLM), logistic regression (LR), and the likelihood-ratio test (bimod). The TN test was an exception, as it requires its own DE test. Since Seurat, Countsplit, and the TN test all produce a *P* value for each gene, we applied the Benjamini-Hochberg (BH) procedure to determine a *P* value cutoff for a given target FDR (e.g., 0.05). Genes with *P* values at or below the cutoff were identified as DE genes.

In the first simulation setting, we simulated target data from a single cell type, mimicking näıve cytotoxic T cells in a PBMC scRNA-seq dataset [38] (Fig. 2a, top left; see “Simulation designs” in the Supplementary Material), representing the most severe double-dipping scenario. Since the data consist of only a single cell type, any identified DE genes are false discoveries. At a target FDR of 0.05, Seurat, Countsplit, and the TN test all failed to control the actual FDR below 0.05 (Fig. 2b). Seurat, which employs a double-dipping approach, performed the worst, with all five DE tests yielding actual FDRs of 1, indicating that Seurat consistently identified false DE genes across 200 simulation runs. Although Countsplit and the TN test were designed to mitigate double dipping, their actual FDRs still far exceeded 0.05 due to anti-conservative *P* values in the presence of gene-gene rank correlations (Fig. S5, middle to right), despite their validity in settings where genes were simulated as independent, as shown in their original studies [11]. In contrast, ClusterDE successfully controlled the FDR below 0.05 for three of the five DE tests: Wilcoxon, t-test, and LR (Fig. 2b). This is supported by contrast scores of genes under the null, which satisfy the required weak symmetry about zero (Fig. S5, left; Figs. S6–S7; see “Theoretical justification of ClusterDE” in the Supplementary Material). While ClusterDE did not control the FDR below 0.05 for the NB-GLM and bimod tests, likely due to violations of their parametric modeling assumptions, the FDR inflation for these tests was substantially less severe than that observed with Countsplit. Specifically, ClusterDE’s actual FDRs were 0.28 and 0.16 for NB-GLM and bimod, respectively, compared to Countsplit’s actual FDRs of 0.68 and 1 for the same tests (Fig. 2b).

**Fig. 2:**
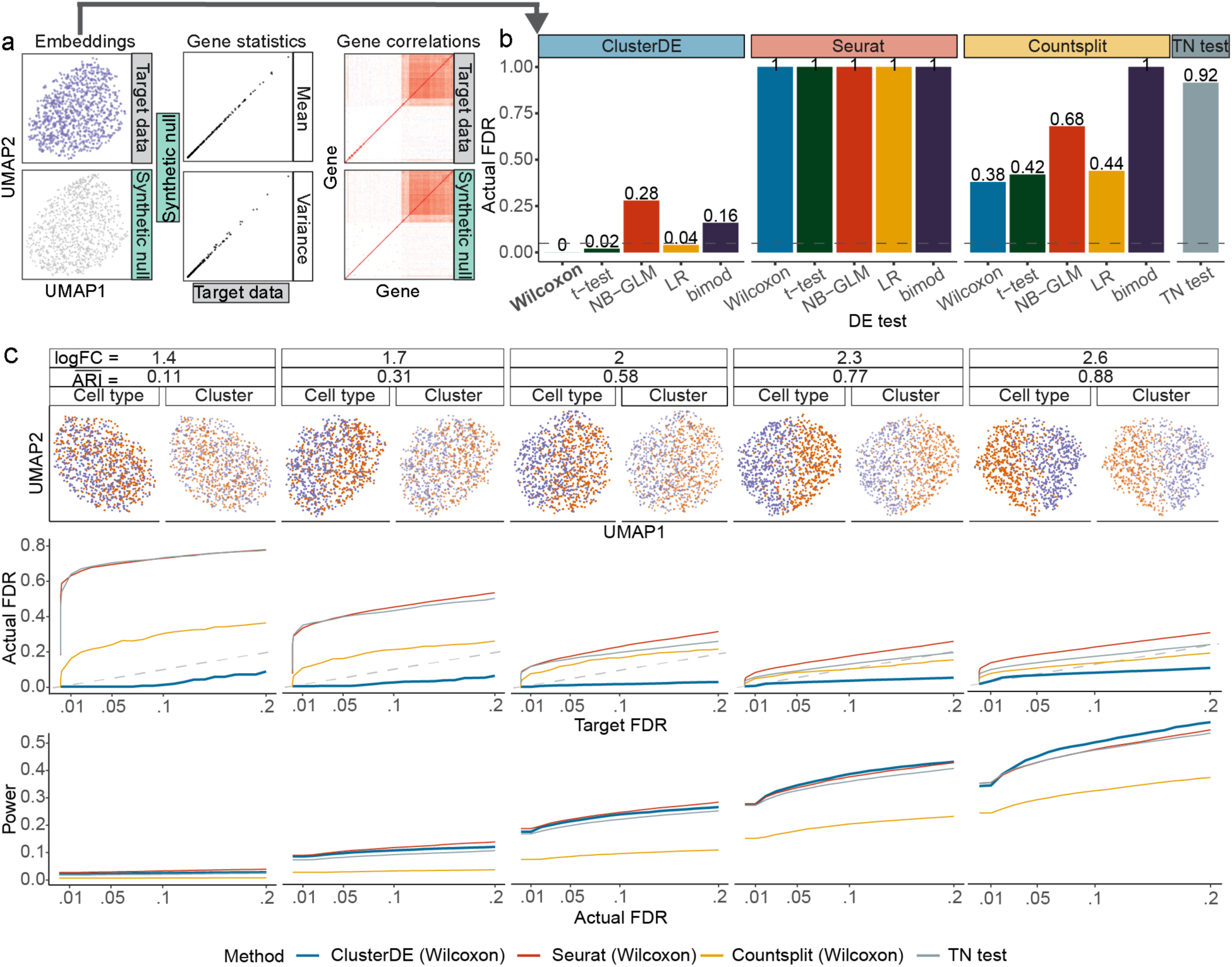
ClusterDE achieves reliable FDR control and good statistical power in simulation studies. **a**, When the target data contain cells from a single type (simulation; see Supplementary Methods “Simulation setting with one cell type” in the Supplementary Material), the synthetic null data generated by ClusterDE resemble the target data well in terms of UMAP cell embeddings (left), per-gene expression mean and variance statistics (middle), and gene–gene rank correlations (right). **b**, On the target data in **a**, ClusterDE (with five DE tests) outperforms three other methods—including Seurat (which involves double dipping), Countsplit (which aims to address double dipping and works with any DE test), and TN test (which aims to address double dipping and has its own DE test)—in FDR control. The horizontal dashed line indicates the target FDR of 0.05. The five DE tests are the Wilcoxon rank-sum test (Wilcoxon), t-test, negative binomial generalized linear model (NB-GLM), logistic regression model predicting cluster membership with likelihood-ratio test (LR), and likelihood-ratio test for single-cell gene expression (bimod). **c**, The FDRs and power of ClusterDE and the three other methods under various severity levels of double dipping. The log fold change (logFC) summarizes the average gene expression difference between two cell types in simulation (see Supplementary Methods “Simulation setting with two cell types and 200 true DE genes” in the Supplementary Material). Corresponding to a small logFC, a small adjusted Rand index (ARI) represents a poor agreement between cell clusters and cell types, representing a more severe double-dipping issue. Across various severity levels of double dipping, ClusterDE controls the FDRs under the target FDR thresholds (diagonal dashed line) and achieves comparable or higher power compared to the three other methods at the same actual FDRs.

In the second simulation setting, we generated datasets with varying degrees of double-dipping, again mimicking näıve cytotoxic T cells from the same PBMC scRNA-seq dataset [38] (Fig. 2c, top; see “Simulation designs” in the Supplementary Material). Each simulated dataset consisted of two cell types with equal numbers of cells and 200 pre-specified true DE genes with varying expression level differences between the cell types, summarized as the log fold change (logFC), where larger logFC values indicate greater distinctions between cell types. After applying Seurat’s default Louvain clustering algorithm to each dataset to identify two clusters, we assessed the agreement between the identified clusters and the true cell types using the adjusted Rand index (ARI), with lower ARI values indicating more inaccurate clustering—that is, a greater mismatch between clusters and cell types (i.e., more severe clustering-induced bias), as visualized by UMAP (Fig. 2c, top row). For DE analysis, we used Wilcoxon, the default DE test in Seurat, as it provided the best FDR control for both ClusterDE and Countsplit, while the TN test employed its own DE test. Results show that ClusterDE consistently controlled the FDR across a range of target thresholds under varying degrees of clustering-induced bias, with larger bias corresponding to lower ARI. In contrast, Seurat, Countsplit, and the TN test failed to control the FDR under the target thresholds and exhibited greater FDR inflation as the bias increased (Fig. 2c, middle row). Notably, ClusterDE achieved comparable or superior statistical power relative to Seurat, Countsplit, and the TN test at their actual FDR levels (Fig. 2c, bottom). These findings held across four other DE tests (t-test, NB-GLM, LR, and bimod) when used with ClusterDE, Seurat, and Countsplit (Fig. S8). To further evaluate performance in realistic scenarios with unbalanced cell type sizes, we simulated datasets with cell type size ratios of 1:4 and 1:9, where ClusterDE still outperformed the other three methods in FDR control across target thresholds and varying degrees of clustering-induced bias. Specifically, ClusterDE consistently demonstrated robust FDR control and comparable or superior statistical power to the other methods when using Wilcoxon as the DE test (Figs. S8–S10).

Technically, ClusterDE and the knockoff methods share the concept of controlling the FDR by generating *in silico* negative control data [39], but their applications differ significantly. The knockoff methods are designed to identify important features in a high-dimensional predictive model under a supervised-learning setting, in contrast to the unsupervised-learning setting addressed by ClusterDE. Broadly, the knockoff methods generate features uncorrelated with the outcome variable given the other features while preserving feature-feature correlations. To assess their applicability to our unsupervised learning setting, we applied the default model-X knockoff method [40] to the target data from the aforementioned simulation, treating genes as features and the cell cluster label as the outcome variable. While the method controlled the FDR, it consistently yielded zero statistical power, making it impractical for DE gene identification. Furthermore, we compared three alternative strategies for synthetic null data generation in Step 1 of ClusterDE, whose default null generator for scRNA-seq is scDesign3 [36]: the model-X knockoff method, a permutation-based approach (where genes are independently permuted across all cells to create a homogeneous cell population), and a variant of ClusterDE using scDesign3 without copula modeling (assuming gene independence). Fig. S11 shows that ClusterDE’s synthetic null data effectively preserved per-gene mean and variance statistics, as well as gene-gene rank correlations. In contrast, the model-X knockoffs method failed to maintain gene mean and variance statistics, while the permutation-based approach and the ClusterDE variant (labeled as “scDesign3 Ind”), as expected, destroyed gene-gene correlations. As a result, only ClusterDE’s synthetic null data mimics the target data while creating a single hypothetical cell type (Fig. S11). Overall, results on simulated datasets demonstrate that ClusterDE achieved the most robust FDR control and the best statistical power among the four strategies for synthetic null generation (Fig. S12).

Building on these comparisons of alternative null-generation strategies, we further evaluated two state-of-the-art simulators, SPARSim and ZINB-WaVE, for generating synthetic null data. To provide practical guidance to users, we implemented a built-in quality-check procedure for synthetic null data covering eleven checks: per-gene mean and variance, gene–gene correlation, normalized expression distributions (via quantile-quantile plots), principal-component variance profiles, cell-embedding similarity (mean local inverse Simpson’s index, mLISI), multivariate expression similarity (Friedman–Rafsky test), clustering structure (silhouette score), and DE results on real data from a single cell line or from two cell types (Fig. S13a–i). The null data generated by ClusterDE and ClusterDE-Fast passed all eleven checks. In contrast, the null data from SPARSim and ZINB-WaVE did not accurately capture gene–gene correlations (Fig. S13c), showed discrepancies in normalized expression distributions and principal-component variance profiles (Fig. S13d– e), did not preserve the global cell topology of the target data (Fig. S13f), produced false discoveries in the cell line data (Fig. S13h), and identified many housekeeping genes as DE genes in the two-cell-subtype data (Fig. S13i).

We evaluated the robustness of ClusterDE to randomness in synthetic null data generation and clustering initialization. Between two independently generated synthetic null datasets, the DE genes identified by ClusterDE were largely overlapping, indicating robustness to randomness in null data generation (Fig. S14). To further enhance stability, we developed ClusterDE-Stable, a stability-enhanced extension of ClusterDE. The procedure generates multiple synthetic null replicates, applies ClusterDE-Core to each, and aggregates the resulting DE calls into gene-level selection frequencies, reporting frequently selected genes as stable DE genes. ClusterDE-Stable exhibited high robustness to randomness in null generation and clustering initialization, as gene selection frequencies obtained from two independent implementations on the same data were highly consistent. Empirically, the reproducibility of stable DE gene detection increased with the number of synthetic null replicates, with good stability achieved using 4—10 replicates (Table S1). We next assessed the robustness of ClusterDE-Stable to inaccuracies in null parameter estimation by perturbing the null model parameters. Specifically, we systematically altered key parameters of the null model (gene means, dispersions, and gene-gene correlations), either individually or jointly, across varying perturbation magnitudes and proportions of perturbed genes. Under these perturbations, the gene-level selection frequencies remained strongly concordant with those obtained under the unperturbed null model (Fig. S15), demonstrating that ClusterDE-Stable is not sensitive to moderate deviations in the estimated null model.

### 2.3 ClusterDE refines cell clusters and enhances marker specificity in scRNA-seq data

We applied ClusterDE to multiple scRNA-seq datasets to examine how it improves the reliability of cell clusters as potential cell types and the biological specificity of cell-type marker identification. These analyses reveal that ClusterDE (i) avoids false-positive markers arising from over-clustering of a homogeneous cell line, (ii) prioritizes canonical markers over housekeeping genes when distinguishing closely related subtypes, and (iii) identifies spurious over-clustered cell-type claims and supports biologically coherent merging and separation of clusters. While ClusterDE-Stable is also applicable, in all analyses below, we focus on ClusterDE-Core, referred to as ClusterDE, to evaluate the performance of the core procedure.

In the first application, we aimed to demonstrate that ClusterDE avoids false-positive cell-type marker discoveries arising from over-clustering when all cells originate from a homogeneous cell type. To this end, we analyzed five pure cell-line datasets (Jurkat, HEK293T, HCC827, H2228, and A549) that are widely used as gold standards of a “single cell type” in benchmarking studies [41, 42] (Fig. 3a, left). While biological heterogeneity within a cell line may include variations in cell cycle, differentiation, or subclonal populations, a cell line is generally not considered to comprise distinct cell subtypes that form separable clusters [43, 44]. Accordingly, any post-clustering DE genes identified within a cell line should not be interpreted as between-cell-type DE genes. These datasets therefore served as negative controls to assess the extent of FDR inflation in existing post-clustering DE workflows and to evaluate whether ClusterDE can eliminate such inflation. As a sanity check, we verified that the synthetic null data generated by ClusterDE closely resembled the target data (Fig. 3a, right; Fig. S16), as confirmed by multiple distributional similarity assessments, including UMAP embeddings, Friedman–Rafsky tests, quantile–quantile plots for top DE genes, preservation of gene–gene rank correlations, and agreement in principal component variance explained. Applying ClusterDE, Seurat, Countsplit, and the TN test to the five datasets, we found that all methods except ClusterDE identified thousands of DE genes, often exceeding 50% of all genes, indicating severe FDR inflation at the target threshold of 5%. In contrast, ClusterDE identified zero DE genes in 22 out of 25 combinations of the five datasets and five DE tests (Wilcoxon, t-test, NB-GLM, LR, and bimod). Notably, ClusterDE consistently identified zero DE genes in all five datasets when using Wilcoxon, which we therefore designate as the default DE test for ClusterDE. To understand the source of these false-positive DE genes, we examined gene-level dropout rates. False-positive DE genes were strongly enriched for low dropout proportions (Fig. S17a), and the same pattern held when restricting to housekeeping genes (Fig. S17b). This is expected: genes with higher expression and variability are more likely to influence clustering and thus be falsely declared as DE, and such genes naturally exhibit fewer zeros. Importantly, removing sparse genes or imputing zeros does not mitigate the FDR inflation problem we observed. In fact, applying an imputation method, scImpute [45], increased the number of false-positive DE genes from 1,023 to 5,150 in the H2228 dataset (Fig. S17c), demonstrating that imputation can exacerbate, rather than resolve, clustering-induced false discoveries.

**Fig. 3:**
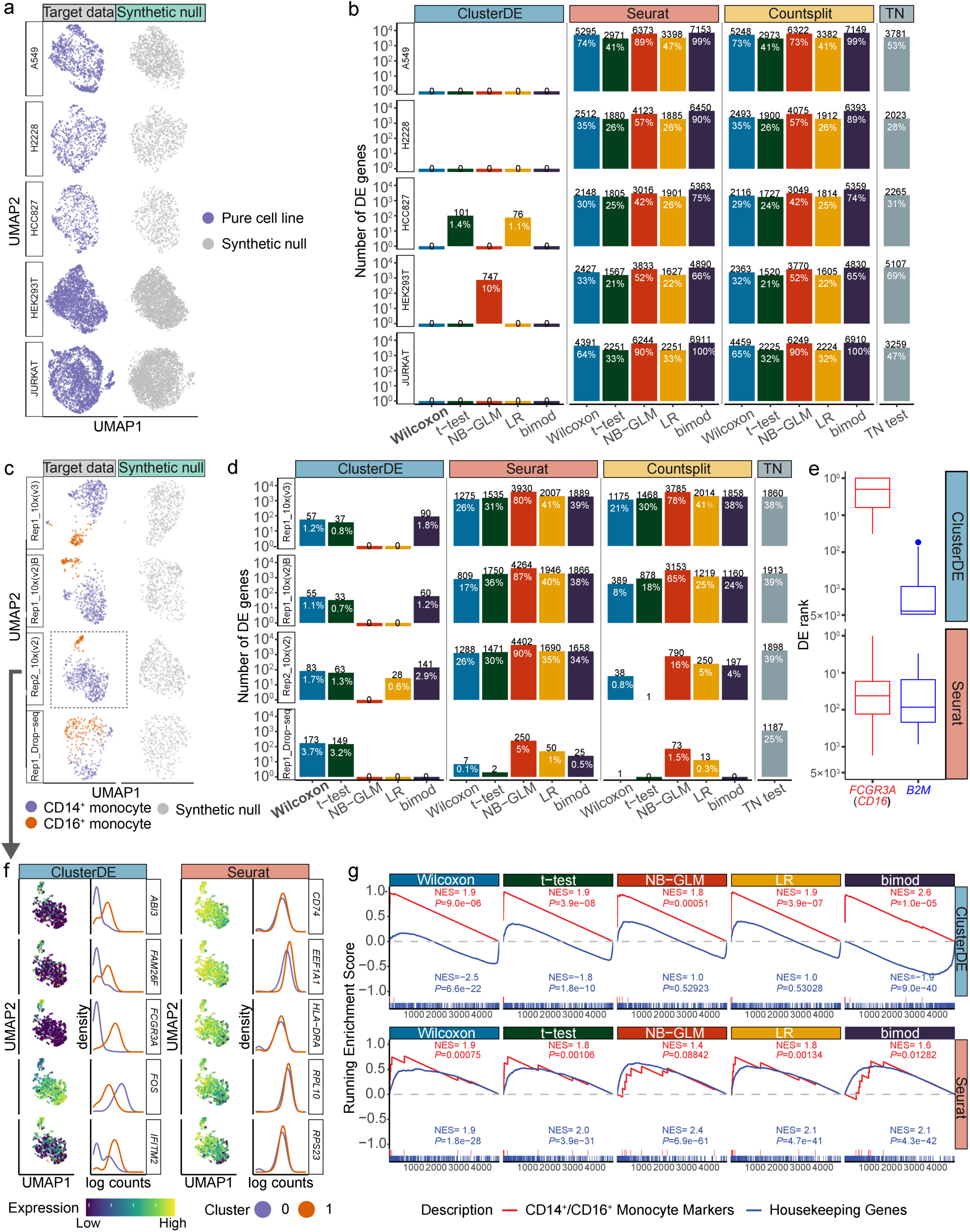
ClusterDE achieves reliable FDR control and good statistical power in identifying cell-type marker genes from real scRNA-seq data. **a**, UMAP visualizations of target data (left) and synthetic null data (right) of five cell lines. **b**, Numbers of DE genes (at the target FDR of 0.05) identified by ClusterDE and the three other methods (Seurat, Countsplit, and TN test). While the three other methods find thousands of DE genes—false positives when interpreted as cell-type marker genes—within a single cell line, ClusterDE makes no false discoveries when used with most DE tests. The numbers in black and white indicate the number of DE genes and the proportions of DE genes among all genes, respectively. The five DE tests are the Wilcoxon rank-sum test (Wilcoxon), t-test, negative binomial generalized linear model (NB-GLM), logistic regression model predicting cluster membership with likelihood-ratio test (LR), and likelihood-ratio test for single-cell gene expression (bimod). **c**, UMAP visualizations of target data (left) and synthetic null data (right) for four datasets containing two monocyte subtypes: CD14^+^ monocytes and CD16^+^ monocytes. The synthetic null data capture the global topology of cells in the target data while filling the gap between the two cell subtypes. The dashed box labels the dataset used in panels **f** and **g**. **d**, ClusterDE identifies DE genes between the two cell subtypes. The numbers in black and white indicate the number of DE genes and the proportions of DE genes among all genes, respectively. **e**, The ranks of two exemplary genes (a monocyte subtype marker *FCGR3A* in red and a well-known housekeeping gene *B2M* in blue) in the DE gene lists of ClusterDE and Seurat across the five DE tests and the four datasets in **c**. In each boxplot representing the distribution of 20 ranks, the center horizontal line represents the median, and the box limits represent the upper and lower quartiles. **f**, The top DE genes identified by ClusterDE exhibit distinct expression patterns in the two cell clusters identified by Seurat clustering, a phenomenon not observed for the top DE genes identified by Seurat. For ClusterDE and Seurat, the top DE genes are defined as the most common DE genes found by the five DE tests in **d** among top 50 genes. The UMAP plots show each top DE gene’s normalized expression levels in the dataset “Rep2 10x(V2)” (marked by the dashed box in **c**; see “Dimensionality reduction and visualization” in the Supplementary Material). The density plots depict each top DE gene’s normalized expression distributions in the two cell clusters. **g**, Gene set enrichment analysis (GSEA) of the ranked DE gene lists identified by ClusterDE and Seurat with five DE tests from the dataset Rep2 10x(V2). The red lines represent the enrichment of the CD14^+^*/*CD16^+^ monocyte marker gene set, and the blue lines represent the enrichment of the housekeeping gene set. The normalized enrichment score (NES) reflects the direction and magnitude of enrichment, and the *P* value indicates the significance of enrichment.

In the second application, we analyzed eight PBMC datasets of CD14^+^/CD16^+^ monocytes [46] to evaluate ClusterDE’s ability to detect known or potential marker genes for the two monocyte subtypes. The datasets were generated from two technical replicates using four UMI-based scRNA-seq protocols (10X Genomics Versions 2 and 3, Drop-seq, and inDrop). After applying the default Seurat clustering to identify two clusters in each dataset, we found that four datasets exhibited relatively accurate clustering results (ARI *>* 0.5; Fig. 3c, left; Fig. S18, top), while the other four showed poor agreement between clusters and the two monocyte subtypes (ARI *<* 0.2; Fig. S18, bottom). Consistent with our motivation, we expected that a reliable post-clustering DE approach should detect meaningful subtype markers only when clustering quality is sufficiently high, because low-quality clusters do not accurately reflect the true distinction between the two cell subtypes.

On the four datasets with high-quality clusters (Fig. S18, top), which align well with the two monocyte subtypes, we expected post-clustering DE genes to reveal cell-subtype markers. As a sanity check, we verified that ClusterDE’s synthetic null data resembled the target data but filled the “gap” between CD14^+^ and CD16^+^ monocytes, representing a single “hypothetical” cell type in each dataset (Fig. 3c, right). When applied to these four datasets, ClusterDE with Wilcoxon identified between 55 and 173 DE genes (corresponding to 1–4% of all genes) at the target FDR of 5%. In contrast, Seurat and Countsplit identified between 1 and 1,288 DE genes (0–26% of all genes), while the TN test consistently identified at least 1,187 DE genes (25% of all genes) (Fig. 3d). Given that CD14^+^ and CD16^+^ monocytes represent related subtypes with modest transcriptional differences, identifying thousands of putative subtype markers is unlikely to be biologically plausible, whereas the number of DE genes reported by ClusterDE is more consistent with biological expectations. On the four datasets with low-quality clusters, Seurat, Countsplit, and the TN test often identified hundreds to thousands of DE genes (Fig. S19), whereas ClusterDE did not identify any DE genes in most cases (Fig. S19), aligning with the expectation that poor clustering should not be interpreted as subtype separation.

Examining the post-clustering DE genes identified by ClusterDE and Seurat on the four high-quality datasets, we found that ClusterDE more effectively separated known subtype marker genes from housekeeping genes. For each dataset, both methods performed DE analysis on the same two clusters obtained from Seurat’s default workflow, with ClusterDE serving as an add-on to correct clustering-induced bias. A clear example is *FCGR3A* (*CD16*), a canonical marker for distinguishing CD14^+^ from CD16^+^ monocytes, and *B2M*, a widely used housekeeping gene [47]. Across the combinations of four datasets and five DE tests, ClusterDE consistently ranked *FCGR3A* among its top DE genes (typically between ranks 1 and 10) and placed *B2M* near the bottom of its DE lists (generally below rank 1,000) (Fig. 3e, top), whereas Seurat ranked both genes similarly (between ranks 10 and 100), obscuring the distinction between a subtype marker and a housekeeping gene (Fig. 3e, bottom). Using the dataset “Rep2 10x(V2)” as an example, we further compared the five most frequently selected DE genes (among the top 50 genes from each of the five DE tests) for ClusterDE and Seurat (Fig. 3f). The genes identified by ClusterDE showed clearer expression differences between the two clusters than those identified by Seurat, indicating that ClusterDE more reliably prioritizes bona fide subtype markers.

To further quantify these differences, we performed gene set enrichment analysis (GSEA) on two gene sets, known CD14^+^/CD16^+^ monocyte markers and housekeeping genes, within the post-clustering DE gene lists from ClusterDE and Seurat. GSEA showed strong enrichment of monocyte markers among ClusterDE’s top-ranked DE genes, clearly separated from housekeeping genes (Fig. 3g, top). Seurat’s top-ranked genes, in contrast, showed weaker marker enrichment and a similar pattern for housekeeping genes, blurring the distinction between the two gene sets (Fig. 3g, bottom); the other three datasets showed the same pattern (Fig. S20). Since researchers commonly focus on only the top *k* DE genes (e.g., *k* = 100), we also counted monocyte markers and housekeeping genes among the top *k* = 1 to 100 genes identified by each method across the five DE tests and four datasets. ClusterDE consistently included more monocyte markers and fewer housekeeping genes than Seurat in this range (Fig. S21). Minus-average (MA) plots [48] help explain this pattern (Fig. S22): four housekeeping genes (*ACTB*, *ACTG1*, *B2M*, *GAPDH*, in blue) had both large target and null DE scores, yielding near-zero contrast scores that excluded them from ClusterDE’s top genes, whereas Seurat ranked them highly based on target DE scores alone. Conversely, four monocyte markers (*CD14*, *FCGR3A*, *MS4A7*, *LYZ*, in red) had large target scores but small null scores, letting ClusterDE correctly identify them as top DE genes.

In the third application, we used a scRNA-seq atlas of the adult *Drosophila* neuronal visual system [49] to illustrate how ClusterDE can guide the merging of spurious clusters caused by over-clustering. The original study clustered the data with the Louvain algorithm at resolution 10, yielding 208 clusters, and used a random forest-based method in Seurat to flag neuron clusters that should be merged. We applied ClusterDE to three such pairs of neuron clusters (Fig. 4a). For all three pairs, ClusterDE identified zero DE genes, supporting the original merging decisions, whereas Seurat (with Wilcoxon as the DE test) reported hundreds of DE genes at the target FDR of 5% (Fig. 4b). Gating plots, commonly used to assess cluster separation based on marker expression, also indicated that each pair should be merged (Fig. 4c). We further examined three genes—*alrm*, *nrv2*, and *Gs2* —that Seurat called DE in at least two of the three pairs. These genes are glial markers [50, 51], unlikely to be expressed in neurons and more plausibly attributable to ambient glial RNA contamination. Together, these results show that ClusterDE can accurately flag over-clustering and avoid false-positive marker calls between spurious clusters.

**Fig. 4:**
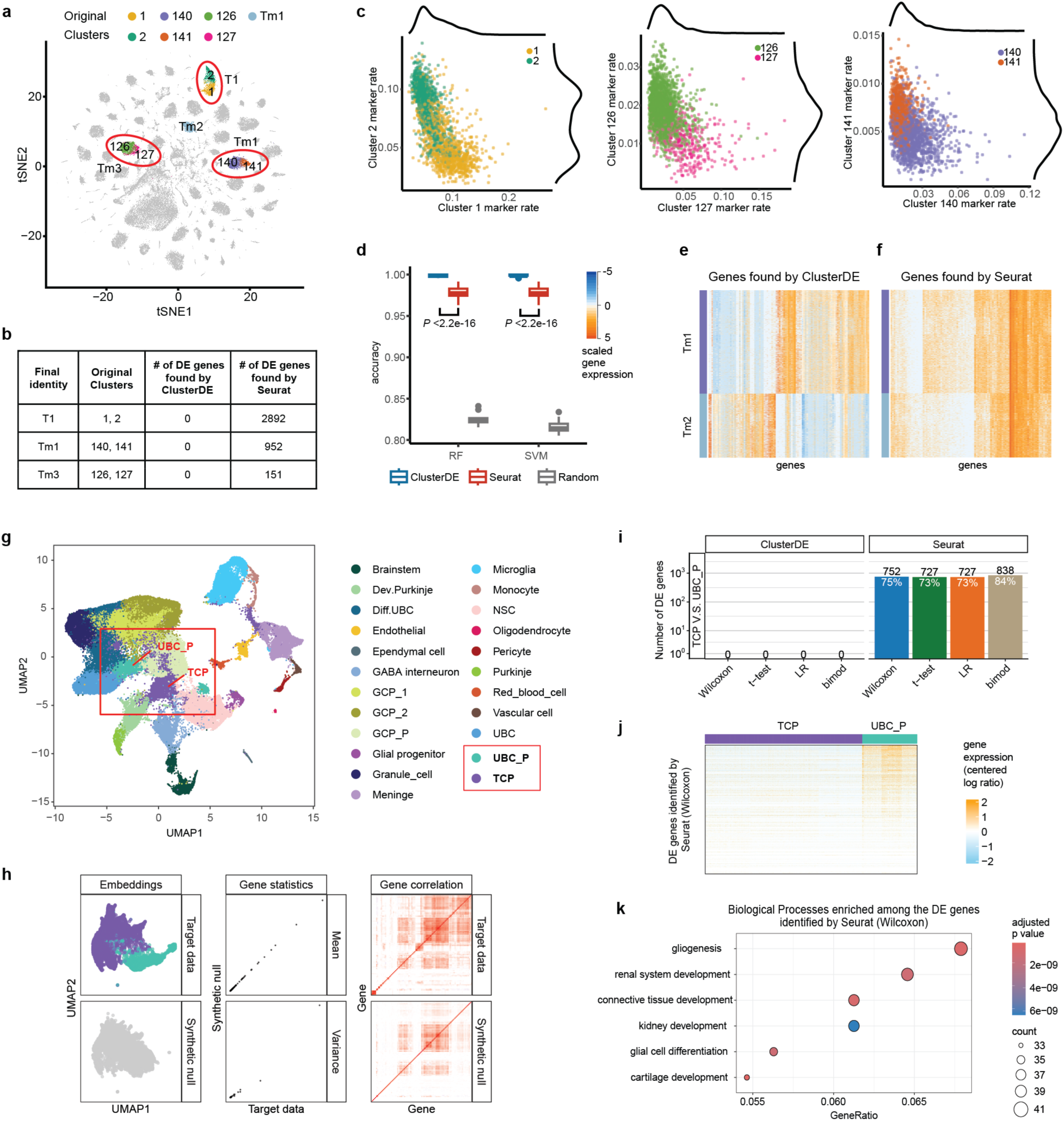
ClusterDE flags over-clustering and refines cell-type marker genes in an adult *Drosophila* visual system scRNA-seq atlas, and invalidates a spurious progenitor cell type in human fetal cerebellar scRNA-seq data. **a–c**, Over-resolved clusters in the adult *Drosophila* visual system atlas. **a**, t-SNE visualization of the data with highlighted cell types: T1 (Clusters 1 and 2), Tm3 (Clusters 126 and 127), Tm1 (Clusters 140 and 141), and Tm2. **b**, Numbers of DE genes (at the target FDR of 0.05) identified by ClusterDE and Seurat (with Wilcoxon as the DE test) for three pairs of neuron clusters flagged for merging. **c**, Gating plots comparing Clusters 1 vs. 2 (left), Clusters 126 vs. 127 (middle), and Clusters 140 vs. 141 (right), showing that each pair is poorly separated and should be merged. **d–f**, Tm1 vs. Tm2 marker gene reliability in the same dataset. **d**, Boxplot showing accuracy achieved by random forest (RF) and support vector machine (SVM) models using the 105 DE genes found by ClusterDE, the top 105 genes (ordered by increasing *P* value) found by Seurat only, and 105 random genes. A two-sided t-test comparing the accuracy obtained using the DE genes from ClusterDE vs. the DE genes from Seurat only shows a significant difference (*P* value *<* 2.2 × 10^-16^*,n* = 100 samples each) for both RF and SVM models. **e**, Heatmap of the 105 DE genes found by ClusterDE. **f**, Heatmap of the top 105 DE genes found by Seurat only. In both **e** and **f**, genes are ordered by hierarchical clustering as implemented by the R function `heatmap.2`. **g–k**, Contested TCP vs. UBC P progenitor claims in human fetal cerebellar scRNA-seq data. **g**, UMAP visualization of the human fetal cerebellar scRNA-seq data with cell type annotations provided by [53]. The purportedly distinct TCP and unipolar brush cell progenitor (UBC P) populations are highlighted. **h**, For the human fetal cerebellar data subset containing the annotated TCPs and UBC Ps, synthetic null data generated by ClusterDE closely resemble the target data in per-gene mean and variance (middle) and gene–gene rank correlations (right) while filling the gap between the two cell populations (left). **i**, Number and proportion of DE genes (at target FDR 0.05) between the annotated TCPs and UBC Ps identified by ClusterDE and Seurat (with DE tests Wilcoxon, t-test, LR, and bimod). Numbers on bars indicate the DE gene count (black) and proportion among all genes (white). **j**, Heatmap displaying the expression profiles of the DE genes identified by Seurat (with Wilcoxon as the DE test) between the annotated TCPs and UBC Ps. **k**, Gene Ontology (GO) enrichment analysis of the DE genes identified by Seurat. The top enriched biological processes (e.g., renal system development, cartilage development) are biologically implausible for a neuronal progenitor transition, confirming that the results from Seurat are driven by artifacts and that ClusterDE correctly suppresses these false positives.

We then analyzed the Tm1 and Tm2 cell types in the same dataset to assess the reliability of cell-type marker genes identified by ClusterDE and Seurat. ClusterDE identified 105 DE genes, whereas Seurat identified 768, including the 105 ClusterDE genes plus an additional 663 genes. To evaluate how well these gene sets distinguish Tm1 from Tm2, we trained binary classifiers (random forest and support vector machine) using three sets of features: the 105 DE genes from ClusterDE, the top 105 DE genes uniquely identified by Seurat (ranked by *P* value), and a random set of 105 genes as a baseline. Tested on held-out cells (20% of the dataset), classifiers based on ClusterDE genes achieved significantly higher accuracy than those based on Seurat’s unique genes (*P* value *<* 2.2 × 10^-16^, two-sided t-test, *n* = 100 train–test splits), with a larger improvement over the random baseline as well. Gene expression heatmaps showed that ClusterDE genes exhibited clearer expression differences between Tm1 and Tm2 than genes identified only by Seurat. GSEA further revealed that ClusterDE genes were enriched in visual system–related pathways, including G protein-coupled receptor (GPCR) activity and eye development, consistent with known roles of Tm1 and Tm2. The enrichment of “positive regulation of response to stimulus” in Tm2 also matches the known delayed response of Tm1 relative to Tm2 [52]. In contrast, DE genes unique to Seurat were enriched in more generic pathways (Fig. S23). These findings indicate that ClusterDE yields more biologically coherent and reliable cell-type marker genes than Seurat.

### 2.4 ClusterDE retrospectively evaluates contested cell-type claims in high-profile datasets

Beyond controlled benchmarks, we asked whether ClusterDE could have flagged cell-type claims that were later disputed in the literature. We examined two such cases from high-profile publications, one ultimately retracted and one challenged by an independent reanalysis. Preprocessing and ClusterDE analysis workflows for these two case studies are detailed in the Supplementary Methods (Sections S1.6.6 and S1.6.7).

In the first case, we considered a single-cell transcriptomic study of the human fetal cerebellum, originally published in *Nature*, in which Luo et al. [53] reported a novel, intermediate cell type termed the “transitional cerebellar progenitor” (TCP). The authors proposed a discrete developmental trajectory starting from neural stem cells (NSCs), passing through TCPs, and then branching into granule cell progenitors (GCPs), unipolar brush cells (UBCs), and Purkinje cells (Fig. 4g). Motivated by this putatively distinct progenitor state, they carried out extensive downstream analyses, including multi-omics profiling, inference of TCP regulatory networks, and predictions that TCPs might serve as the cellular origin for aggressive subgroup 3 medulloblastoma. Subsequently, Smith et al. [54] argued that the TCP population was a technical artifact. Re-evaluating the original dataset, they showed that TCP cells arose almost entirely from a single biological sample (PCW12) and exhibited profiles consistent with low-quality droplets contaminated by ambient RNA. When the PCW12 sample was removed, the distinct TCP cluster disappeared, and the remaining TCP cells merged into UBC progenitor (UBC P) and GCP populations. As a result, the core TCP-related claims in Luo et al. [53] were deemed unsupported, leading to the retraction of the study.

To formally test whether TCPs represent a distinct cell type, we applied ClusterDE to the subset of cells annotated as TCP and UBC P. ClusterDE first generated synthetic null data that preserved key characteristics of the target data while filling the gap between the two cell populations (Fig. 4h). When we performed DE analysis on this subset, ClusterDE identified exactly zero DE genes between TCP and UBC P, indicating that the two annotated populations are transcriptionally indistinguishable and that TCPs do not constitute a distinct cell type. In contrast, Seurat reported 727–838 significant DE genes across four DE tests (Fig. 4i). Inspection of the 752 DE genes identified by Seurat with Wilcoxon as the DE test revealed only weak separation between TCP and UBC P in these genes’ expression (Fig. 4j). Gene Ontology (GO) analysis of these genes further highlighted their implausibility: top biological process terms included renal, kidney, and cartilage development—organ systems unrelated to cerebellar neurons—as well as gliogenesis and glial cell differentiation, implying a divergent glial lineage that contradicts the established neuronal fate of both TCPs and UBC Ps (Fig. 4k). Together, these results show that ClusterDE would have flagged the TCP cluster as spurious at the time of analysis and could have prevented the substantial downstream effort based on this false-positive cell type.

In the second case, we examined a single-cell RNA-seq study of peripheral immune responses to COVID-19, in which Wilk et al. [55], originally published in *Nature Medicine*, profiled PBMCs from patients with severe disease. The authors annotated a purportedly novel “developing neutrophil” population enriched in patients with acute respiratory distress syndrome (ARDS). Based on their dimensionality reduction results, this population appeared to form a continuous bridge with plasmablasts (Fig. S24a), and they proposed a transdifferentiation trajectory connecting these two cell types, which belong to distant lymphoid and myeloid lineages. The existence of developing neutrophils and their continuous trajectory from plasmablasts was presented as a distinctive pathophysiological feature of ARDS in severe COVID-19. This transdifferentiation claim was later challenged by Alquicira-Herńandez et al. [56], who showed that the reported continuum was not reproducible in other patients with mild or severe COVID-19 and could be a computational artifact. In particular, when cell-cycle signals were properly regressed out during normalization, the apparent bridge between plasmablasts and developing neutrophils vanished.

To rigorously assess this trajectory, we applied ClusterDE to the annotated IgM plasmablasts (IgM PBs) and developing neutrophils. After we validated the synthetic null data (Fig. S24b), ClusterDE identified 163 DE genes between the two populations (Fig. S24c), indicating that IgM PBs and developing neutrophils are discrete, transcriptomically distinct cell types rather than adjacent cells along a continuous transdifferentiation path. Seurat also reported many DE genes, but the top DE genes identified by ClusterDE exhibited much sharper expression contrasts between the two cell populations than those identified only by Seurat with Wilcoxon as the DE test (Fig. S24d). These results show that ClusterDE both confirms the discrete nature of these immune cell types and provides high-contrast transcriptomic signatures that clearly separate them.

### 2.5 ClusterDE generalizes to hierarchical cell types, atlas-scale taxonomies, and cell trajectories in scRNA-seq data

In many biological systems, cell types have a hierarchical relationship, with broad cell types comprising finer subtypes through branching. Over-clustering can affect this hierarchy construction at any branch, producing spurious leaf clusters that should be merged into their more distinct parent clusters. To address this issue by refining clusters and identifying marker genes for the reliable clusters that remain, we developed ClusterDE-Tree, a hierarchical extension of ClusterDE that operates directly on the cluster tree produced by a standard clustering pipeline (e.g., Seurat).

ClusterDE-Tree evaluates each sibling split in the cluster tree, starting from the leaves, using a Ward linkage statistic calibrated against an empirical null built from multiple synthetic null datasets [23]; branches with statistically unsupported splits are merged, while supported splits are retained. Marker genes for the resulting refined clusters are then identified using ClusterDE-Stable (see “ClusterDE-Tree details” in the Supplementary Methods). We applied ClusterDE-Tree to the PBMC scRNA-seq dataset [38] clustered at two Louvain resolutions: a lower resolution (0.2) yielding fewer, broader clusters, and a higher resolution (1.0) yielding many finer clusters, some of which are over-clusters (Fig. 5a–b). ClusterDE-Tree correctly flagged the over-clustered branches at the higher resolution as statistically unsupported and merged them, bringing the two cluster trees into close agreement after refinement, while recovering canonical PBMC cell-type marker genes at the branches that remained (Fig. 5a–b). These results show that ClusterDE-Tree yields cell-type annotations that are robust to the choice of clustering resolution and respect the underlying cell-type hierarchy, without requiring users to arbitrarily select which cluster pairs to compare for merging decisions. Moreover, ClusterDE-Tree provides a systematic procedure for defining cell-type marker genes within a cell-type hierarchy.

**Fig. 5:**
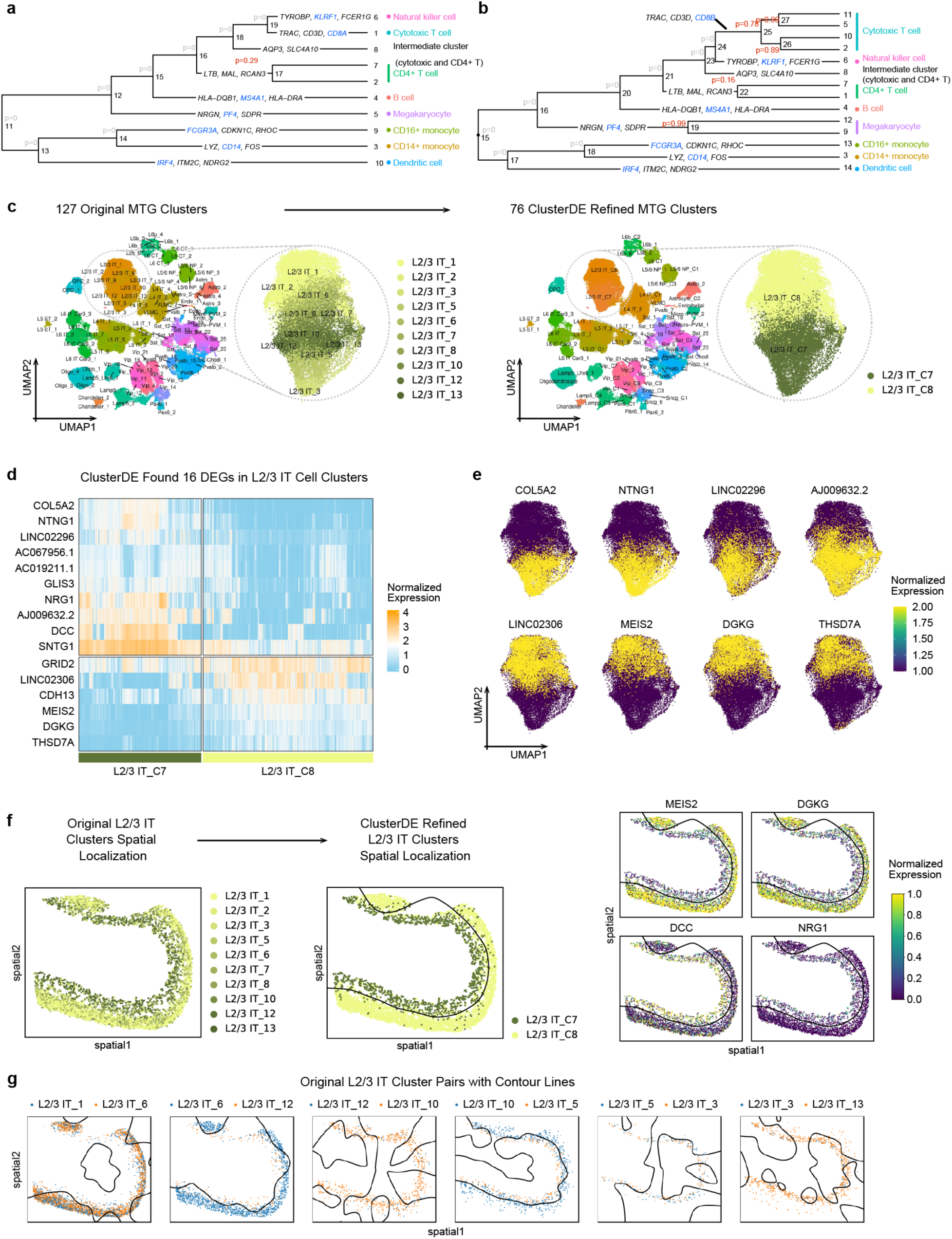
ClusterDE-Tree refines PBMC cluster hierarchies and the Allen Human MTG cell-type taxonomy and validates refined MTG clusters with SRT. **a–b**, ClusterDE-Tree provides statistical significance for branching events and marker gene identification in PBMC scRNA-seq cluster hierarchies. **a**, Cluster hierarchy constructed from a PBMC scRNA-seq dataset using a lower Louvain resolution (0.2), yielding fewer, broader clusters. **b**, Cluster hierarchy of the same PBMC scRNA-seq dataset constructed using a higher Louvain resolution (1.0), producing a larger number of finer clusters, some of which are over-clusters. ClusterDE *P* -values indicate whether each branching event represents a statistically significant split of a cluster into two, with insignificant *P* -values that drive cluster merging highlighted in red. ClusterDE-Tree effectively merges spurious clusters, bringing the two cluster hierarchies into agreement after refinement and recovering canonical PBMC cell-type marker genes (highlighted in blue). **c–g**, ClusterDE-Tree refines the Allen Human MTG cell-type taxonomy and supports refined clusters using an external MERFISH dataset. **c**, UMAP visualization of the Allen Human MTG snRNA-seq atlas colored by the original cell cluster annotations (left) and the ClusterDE-refined annotations (right). The circled region and corresponding zoomed-in view highlight L2/3 intratelencephalic (L2/3 IT) neurons, which exhibited the largest number of cluster merges during ClusterDE-Tree refinement. **d**, Expression heatmap of the 16 differentially expressed genes identified by ClusterDE between the two refined L2/3 IT cell clusters. Cells are grouped according to the refined annotations. **e**, UMAP projection of representative ClusterDE marker genes identified between the two refined L2/3 IT cell clusters. **f**, Spatial validation in the Allen Human MTG MERFISH dataset. The left panel shows spatial mapping using the original cluster annotations. The middle panel shows spatial mapping using the ClusterDE-refined annotations, with solid contour lines generated from kernel density estimation (KDE) using zero-density level sets. The right panel shows spatial expression of representative ClusterDE marker genes identified from the snRNA-seq reference atlas and projected onto the MERFISH dataset. **g**, Spatial mapping of six representative pairs of L2/3 IT clusters based on the original annotations (the left panel in **f**), with solid contour lines generated from KDE using zero-density level sets, illustrating that many of the original L2/3 IT clusters are not spatially separable.

Beyond this proof-of-concept PBMC dataset, we evaluated whether ClusterDE-Tree could refine a cell-type hierarchy in a real atlas-scale setting. We applied ClusterDE-Tree to single-nucleus RNA-seq (snRNA-seq) data from five donors in the Allen Human Middle Temporal Gyrus (MTG) taxonomy. Starting from 127 transcriptomic “supertypes” [57], ClusterDE-Tree merged statistically unsupported splits to yield 76 refined clusters. The largest refinement occurred among layer 2/3 intratelencephalic (L2/3 IT) neurons, an excitatory cortical projection-cell class, where ten original clusters were reduced to two refined clusters (Fig. 5c). ClusterDE identified 16 DE genes between the resulting L2/3 IT C7 and L2/3 IT C8 clusters with clear expression contrasts (Fig. 5d); eight representative genes showed expression patterns aligned with their separation in the snRNA-seq UMAP (Fig. 5e).

We then evaluated this transcriptome-based refinement in an independent Allen Human MTG MERFISH dataset [58]. Although no spatial information was used to construct the snRNA-seq hierarchy, spots assigned to L2/3 IT C7 and L2/3 IT C8 formed spatially segregated regions consistent with the spatial expression of their snRNA-seq marker genes (Fig. 5f). In contrast, spots assigned to the original ten L2/3 IT supertypes had substantially overlapping spatial distributions (Fig. 5g). Thus, ClusterDE-Tree retained two clusters supported both by transcriptomic marker contrasts and by spatial coherence in an independent dataset.

ClusterDE-Tree is not intended to replace biologically informed fine-grained atlas annotations, which may incorporate evidence from additional modalities, developmental context, morphology, connectivity, or prior knowledge. Rather, it asks a narrower question: at what resolution are subdivisions statistically supported as distinct populations by the transcriptomic data used to define them? Without such a criterion, clustering can subdivide cells indefinitely, and conventional post-clustering differential analysis can yield apparent markers for increasingly fine partitions that may not be robust or transferable. ClusterDE-Tree supplies a data-driven stopping rule by retaining a split only when it is distinguishable from a homogeneous synthetic null population and merging unsupported splits. We therefore do not claim that the original supertypes lack biological relevance; rather, the Allen analysis shows that, at this resolution, their finer subdivisions are not supported as distinct transcriptomic populations by snRNA-seq alone.

We next extended the same synthetic null calibration principle to a related problem: clustering-induced bias propagating into downstream trajectory inference. Slingshot [59], a popular trajectory-inference algorithm, relies on pre-inferred clusters; over-clustering can therefore create artificial bifurcations and spurious DE signals that incorrectly “validate” such bifurcations. To illustrate this issue, we compared a standard clustering→trajectory→DE workflow with ClusterDE on two datasets known not to contain true bifurcating trajectories (i.e., negative controls). The details of applying ClusterDE to trajectory analysis are described in the “ClusterDE trajectory-based DE” subsection (Section S1.4.5) and in the “Trajectory analysis details” subsection (Section S1.6.9) of the Supplementary Methods. The first is a homogeneous lung adenocarcinoma cell line (H2228) [42] (Fig. S25a–c), which lacks any developmental trajectory. The second is a human embryonic glutamatergic neurogenesis dataset [60] (Fig. S25d–f), where cells progress along a single continuous lineage from radial glia to neuroblasts. Using Slingshot followed by Wilcoxon rank-sum testing between inferred terminal clusters, the standard pipeline identified 5,560 DE genes (77% of all genes) in H2228 and 1,216 DE genes in the neurogenesis dataset. In contrast, ClusterDE identified zero DE genes in both cases (Fig. S25b,e), correctly indicating that the inferred bifurcations were artifacts of over-clustering. These negative-control analyses illustrate how clustering-induced bias can mislead bifurcation inference and between-trajectory DE analysis, and they highlight ClusterDE’s ability to suppress such false-positive trajectory markers.

Having established that ClusterDE suppresses false bifurcations in negative controls, we next asked whether it retains sensitivity to true lineage structure, evaluating it on a well-characterized mouse bone marrow dataset [61], in which common myeloid progenitors (CMPs) bifurcate into granulocyte–monocyte progenitors (GMPs) and erythroid precursors. ClusterDE generates synthetic null data representing a hypothetical single-lineage scenario (Fig. S26a, middle and bottom) to assess whether observed lineage differences are statistically supported. Compared with the standard pipeline, ClusterDE more accurately identified genes with clear lineage-specific expression patterns. The top DE genes from ClusterDE showed clear separation between the two lineages (Fig. S26b), whereas genes uniquely reported by the standard pipeline exhibited ambiguous differences and are likely false positives (Fig. S26c). Known lineage markers—including *Klf1*, *Elane*, *Gata1*, *Fcgr3*, *Cebpa*, and *Cd34* —received consistently higher rankings under ClusterDE (Fig. S26d), and heatmaps along pseudotime further demonstrated that genes identified by ClusterDE display clear lineage-specific expression, whereas those unique to the standard pipeline do not (Fig. S26e).

Together, these results show that ClusterDE generalizes beyond pairwise cluster comparisons to hierarchical cell-type annotation, crucial for cell atlas construction, and to trajectory inference, both settings where over-clustering artifacts must be suppressed to avoid spurious biological conclusions. Practical guidelines for applying ClusterDE-Tree to cluster hierarchies, including atlas-scale taxonomies, and for applying ClusterDE to trajectory-based analyses are provided in the Supplementary Methods (Sections S1.2.3 and S1.2.4).

### 2.6 ClusterDE reveals reliable spatial-domain marker genes in SRT data

Having shown that ClusterDE resolves double-dipping and over-clustering in scRNA-seq cluster hierarchies and trajectories, we next asked whether the same framework could address analogous issues in SRT. In spatial domain detection and annotation, spatial clustering (e.g., using BayesSpace [62]) is first applied to SRT data to infer putative spatial domains (referred to as “spatial clusters”). DE analysis is then performed to identify potential domain marker genes, creating a double-dipping issue analogous to that in scRNA-seq analysis. We define “spatial-domain marker genes” as genes exhibiting discontinuous expression changes between adjacent domains while showing elevated expression in one domain relative to others (Fig. S27). In contrast, some genes exhibit spatial variation but vary smoothly across domain boundaries and therefore should not be considered spatial-domain markers (Fig. S27). While ClusterDE-Stable is also applicable, in all analyses below we focus on ClusterDE-Core, referred to simply as ClusterDE, to evaluate the performance of the core procedure.

Although both scRNA-seq and spatial clustering rely on expression profiles, spatial clustering algorithms additionally incorporate spatial coordinates, inducing spatial dependence among neighboring spots. ClusterDE accounts for this dependence when generating synthetic null SRT data (see “Supplementary Methods” in the Supplementary Material). Using both simulated and real SRT data, we demonstrated that ClusterDE effectively resolves double dipping in post-spatial clustering DE analysis and facilitates the identification of reliable and interpretable spatial-domain marker genes.

We first simulated an SRT dataset containing a single spatial domain (see “Supplementary Methods” in the Supplementary Material) and applied ClusterDE as an add-on to a commonly used pipeline consisting of BayesSpace for clustering and Seurat’s Wilcoxon or t-test for DE analysis (referred to as BayesSpace+Seurat). Because the dataset contains only one domain, no spatial-domain marker genes should be detected. However, only ClusterDE met this expectation, whereas BayesSpace+Seurat identified hundreds of false-positive spatial-domain marker genes, failing to control the target FDR of 0.05 (Fig. S28a). Examination of the *P* values from BayesSpace+Seurat revealed an anti-conservative bias, whereas ClusterDE’s contrast scores were symmetric around zero (Fig. S28b), fulfilling the symmetry condition required for valid FDR control. Thus, in the single-domain setting, ClusterDE avoided spurious discoveries while BayesSpace+Seurat did not.

We next simulated two datasets containing more than one domain: (1) a two-domain dataset with 200 pre-specified true spatial-domain marker genes and (2) a three-domain dataset with 200 and 168 true marker genes for each pair of adjacent domains, respectively. These marker genes defined clear spatial boundaries that were approximately captured by BayesSpace clustering (Fig. S28c for two domains; Fig. 6a for three domains). Because ClusterDE compares two spatial clusters at a time, for the three-domain dataset we analyzed Layer 5 vs. Layer 6 and Layer 6 vs. WM separately. In both the two- and three-domain settings, BayesSpace+Seurat again failed to control the FDR at 0.05 (Fig. S28d; Fig. S29). In contrast, ClusterDE maintained valid FDR control while achieving high power to detect most true spatial-domain marker genes. Visualization of top-ranked genes revealed that those identified by ClusterDE displayed clear, discrete expression boundaries, whereas those from BayesSpace+Seurat lacked such specificity (Fig. 6b). These simulation results confirm that ClusterDE reliably identifies spatial-domain marker genes with enhanced specificity.

**Fig. 6:**
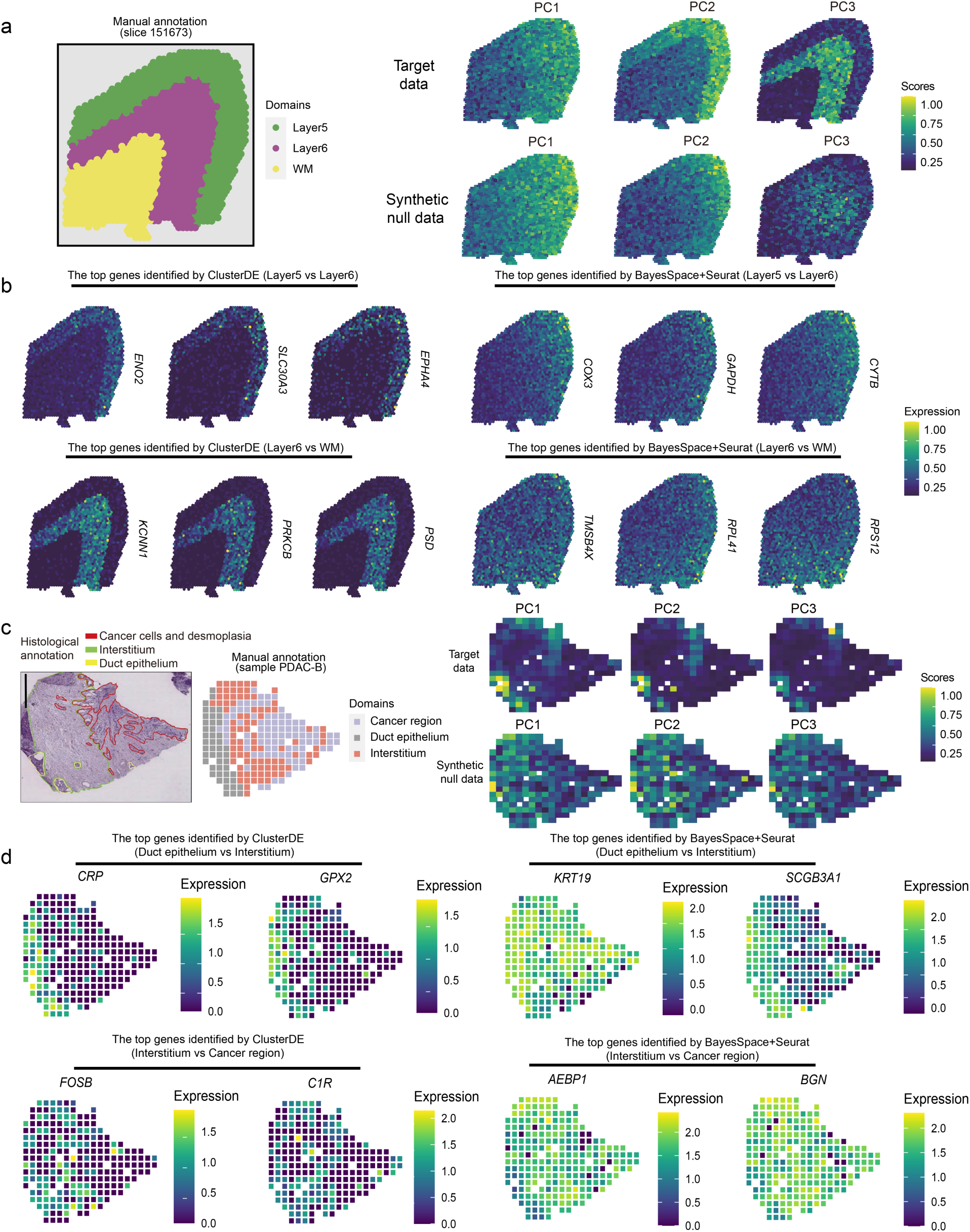
ClusterDE mitigates double dipping and identifies reliable spatial-domain marker genes in SRT post-clustering DE analysis. ClusterDE demonstrates robust statistical power in identifying spatial-domain marker genes from both a simulated SRT dataset containing three domains and the PDAC-B cancer slice from the human primary pancreatic cancer (PDAC) SRT dataset. **a**, Visualization of annotated spatial domains alongside the spatial expression patterns of three representative principal components (PCs) among the top five PCs in the target data (simulated) and synthetic null data. **b**, Spatial expression patterns of top domain marker genes. The first and second rows display the expression of top domain marker genes of Layer 5 and Layer 6, respectively, detected by ClusterDE and BayesSpace+Seurat (BayesSpace for clustering and Seurat for DE analysis). **c**, Visualization of annotated domains alongside the spatial expression patterns of three representative PCs among the top five PCs in the target data (PDAC-B) and synthetic null data. **d**, Spatial expression patterns of top domain marker genes. The first and second rows display the expression of top domain marker genes of the Duct Epithelium and Interstitium, respectively, detected by ClusterDE and BayesSpace+Seurat.

We next applied ClusterDE to real SRT datasets, including slices from the human dorsolateral prefrontal cortex (DLPFC) [63] and the human primary pancreatic cancer (PDAC) dataset [64]. For the DLPFC dataset, we first examined Layers 1, 4, and 5, each of which corresponds to a homogeneous spatial domain based on manual annotation. Within each layer, BayesSpace identified two clusters, and we compared these clusters using both ClusterDE and BayesSpace+Seurat. ClusterDE identified nearly zero DE genes, whereas BayesSpace+Seurat identified hundreds to thousands (Fig. S30), indicating severe double-dipping effects and demonstrating ClusterDE’s robustness.

We then applied both methods to the full DLPFC slice using BayesSpace without constraining the number of clusters. Several annotated layers were subdivided into smaller clusters. Focusing on Layers 3 and WM, we compared pairs of clusters within each annotated domain. ClusterDE again identified no DE genes, supporting the merging of these clusters and correcting over-clustering (Fig. S31). In contrast, BayesSpace+Seurat frequently reported DE genes between clusters that should not be distinct domains, illustrating its susceptibility to double-dipping. These analyses demonstrate that ClusterDE prevents the erroneous creation of spurious spatial domains and avoids false marker identification.

Finally, in the PDAC dataset (Fig. 6c), we evaluated two pairs of adjacent spatial domains: duct epithelium vs. interstitium and interstitium vs. cancer. ClusterDE’s top-ranked marker genes showed clear domain specificity, whereas those identified by BayesSpace+Seurat showed smooth gradients inconsistent with domain boundaries (Fig. 6d). For example, *CRP*, identified by ClusterDE, was expressed exclusively in duct epithelium, while top-ranked genes from BayesSpace+Seurat lacked domain specificity. Examination of known marker and housekeeping genes further supported these findings: ClusterDE ranked marker genes (e.g., *VWA1*, *POSTN*, *SFRP2*, *F3*) substantially higher, and ranked housekeeping genes (e.g., *TMSB10*, *S100A6*) substantially lower than BayesSpace+Seurat (Fig. S32).

In summary, these simulation and real-data results show that ClusterDE provides reliable post-clustering DE analysis in SRT data, consistently identifying spatial-domain marker genes with clear, biologically meaningful boundaries while suppressing false positives from double dipping and over-clustering in normal brain and pancreatic cancer. Practical guidelines for applying ClusterDE to SRT data are provided in Section S1.2.5 of the Supplementary Methods.

### 2.7 ClusterDE corrects clustering-induced bias across diverse omics modalities

Clustering followed by differential analysis is widely used beyond scRNA-seq and SRT. In many omics workflows, including single-cell multi-omics, population-scale bulk transcriptomics, and microbiome data, the same data modality (or correlated modalities) is used both to define clusters and to identify cluster-specific features. This practice can introduce clustering-induced bias, leading to inflated false-positive discoveries and overconfident biological interpretations. To assess the generality of ClusterDE as a calibration framework for post-clustering inference, we evaluated it across three additional omics settings.

### Single-cell multi-omics

In single-cell multiome assays (joint profiling of RNA-seq and ATAC-seq from the same cells), clustering is typically performed using the RNA modality, whereas differential accessibility analysis is conducted on the ATAC modality [33, 34, 35]. Although this cross-modality workflow avoids literal double dipping, it remains vulnerable to clustering-induced spurious differential accessibility signals when RNA-based clusters do not reflect distinct subpopulations. Because RNA and ATAC signals are correlated, over-clustering the RNA modality can propagate artifacts into the ATAC modality. Methods such as Countsplit [11], which rely on splitting a single count matrix, are not applicable here and cannot resolve cross-modality artifacts.

To illustrate this issue, we analyzed a single-cell multiome dataset of the GM12878 human lymphoblastoid cell line, a widely used homogeneous control with matched RNA-seq and ATAC-seq profiles for 11,437 cells [65]. When cells were over-clustered into two groups based on RNA expression using Seurat’s default Louvain clustering, the artificial split propagated to the ATAC-seq modality: differential accessibility testing identified 1,335 significant regions, including a peak near *HIST1H1C* (Fig. S33a). Gene Ontology analysis revealed enrichment for muscle-related pathways (Fig. S33b), contradicting the known lymphoblastoid identity of GM12878. These false discoveries arise from over-clustering combined with cross-modality correlation, rather than double dipping.

ClusterDE evaluated whether the RNA-defined clusters represented genuine subpopulations. The synthetic null data closely resembled the RNA-seq modality (Fig. S33c), and ClusterDE identified zero DE genes, correctly indicating that the two RNA clusters were not biologically meaningful. In contrast, standard post-clustering DE analysis on the RNA modality reported thousands of DE genes (Fig. S33d), which would falsely validate the spurious clusters and propagate artifacts into the ATAC modality. These results demonstrate that ClusterDE suppresses over-clustering artifacts at their source, preventing the propagation of false discoveries in downstream multi-omics analyses.

Having shown that ClusterDE correctly avoids false-positive marker identification in a negative control case (a cell line) where the clusters are known to be spurious, we next asked whether it retains power in identifying differentially accessible (DA) regions (called peaks) when true cell subpopulations exist. We analyzed a single-cell multiome dataset of human PBMCs, applying ClusterDE to naive CD4 T cells (CD4 Naive, 1,419 cells) and CD4 central memory T cells (CD4 TCM, 1,149 cells). The synthetic null data generated by ClusterDE closely resembled both the RNA and ATAC modalities (Fig. 7a, c). ClusterDE identified a smaller set of DE genes and DA peaks than the standard post-clustering Wilcoxon rank-sum test, consistent with the moderate transcriptomic and epigenomic differences expected between these related T-cell subtypes (Fig. 7b, d). The DA peaks identified by ClusterDE also displayed stronger subtype-specific contrasts than those from the standard post-clustering Wilcoxon rank-sum test alone (Fig. 7e). Fold-enrichment analysis of the ClusterDE-selected DA peaks showed overall stronger enrichment of known T-cell transcription factor binding motifs (Fig. 7f), further demonstrating that ClusterDE improves DA peak marker specificity when true cell subtypes are present.

**Fig. 7:**
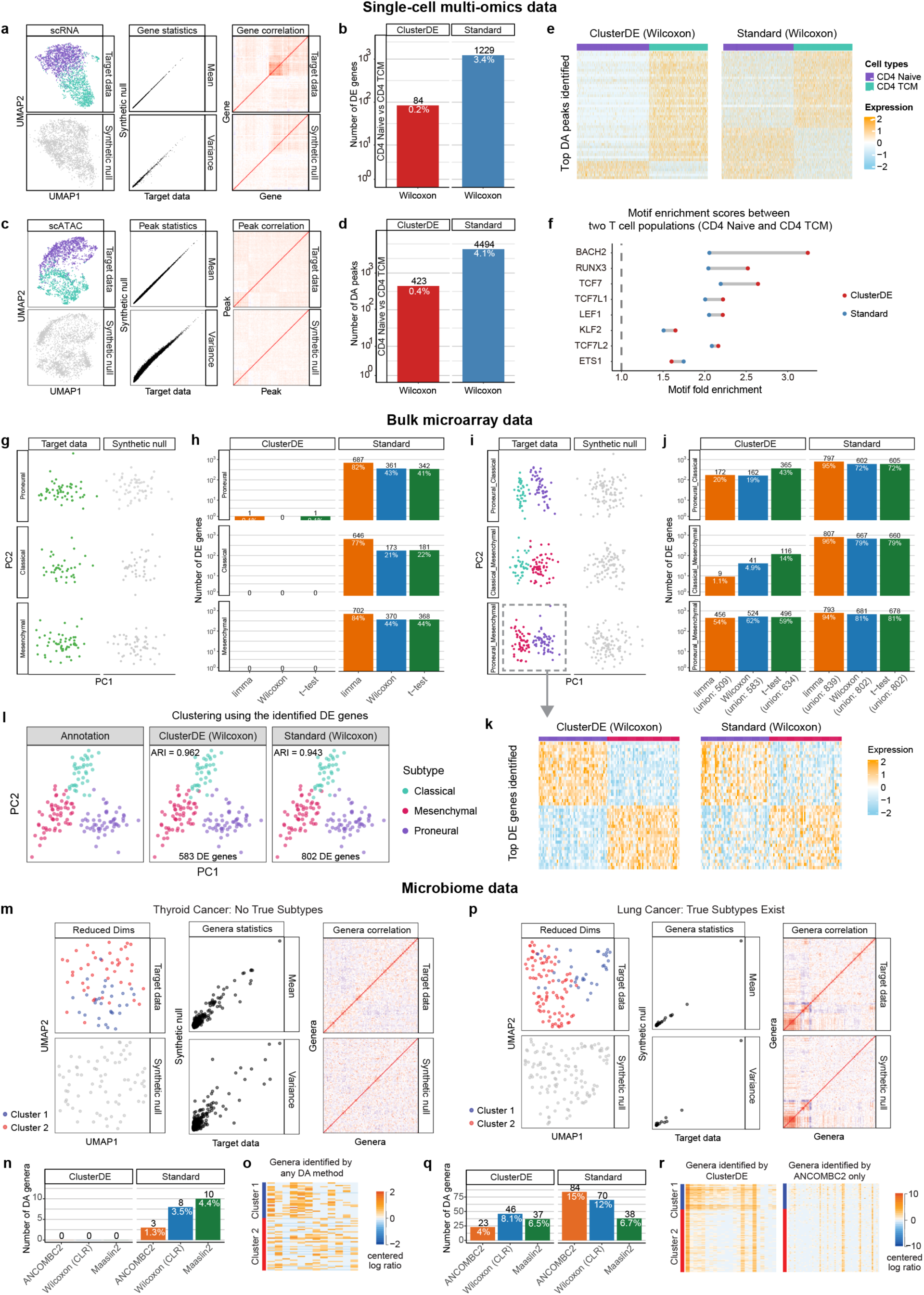
ClusterDE corrects clustering-induced bias and enhances post-clustering inference across diverse omics modalities. **a-f**, ClusterDE identifies biologically meaningful differentially accessible (DA) peaks and DE genes in single-cell multiome data when true cell subtypes are present. Analysis was performed on a single-cell multiome dataset of human PBMCs, applying ClusterDE to naive CD4 T cells (CD4 Naive, 1,419 cells) and CD4 central memory T cells (CD4 TCM, 1,149 cells). **a**, ClusterDE generates a synthetic null dataset that closely resembles the RNA modality in UMAP (left), per-gene mean and variance (middle), and gene–gene correlation structure (right). **b**, Bar plot comparing the number and percentage (black and white text, respectively) of DE genes identified by ClusterDE versus the standard post-clustering Wilcoxon rank-sum test, consistent with the moderate transcriptomic differences expected between these related T-cell subtypes. **c**, ClusterDE generates a synthetic null dataset that closely resembles the ATAC modality in UMAP (left), per-peak mean and variance (middle), and peak–peak correlation structure (right). **d**, Bar plot comparing the number and percentage (black and white text, respectively) of DA peaks identified by ClusterDE versus the standard post-clustering Wilcoxon rank-sum test, consistent with the moderate epigenomic differences expected between these related T-cell subtypes. **e**, Heatmaps of the top DA peaks between CD4 Naive and CD4 TCM cell types identified by ClusterDE with the Wilcoxon test (left), which constitute a subset of those identified by the standard Wilcoxon rank-sum test, and of the top DA peaks identified only by the standard Wilcoxon rank-sum test (right), showing that the ClusterDE-selected DA peaks display stronger subtype-specific contrasts. **f**, Fold-enrichment comparison of known T-cell transcription factor binding motifs for eight transcription factors associated with CD4 T-cell differentiation, comparing DA peak sets identified by ClusterDE (red) and the standard Wilcoxon rank-sum test (blue), showing overall stronger motif enrichment for the ClusterDE-selected DA peaks. **g-l**, ClusterDE controls false positives and retains power in bulk microarray analyses of glioblastoma (GBM) subtypes. **g**, PCA visualizations of the target data (left) and synthetic null data (right) for three GBM subtypes. **h**, Number and proportion of DE genes (at target FDR 0.05) identified by ClusterDE and standard post-clustering DE tests (limma, Wilcoxon, and t-test) within each GBM subtype analyzed separately. Numbers on bars indicate the DE gene count (black) and proportion among all genes (white). **i**, PCA visualizations of the target and synthetic null data for three GBM subtype pairs. **j**, Number and proportion of DE genes (at target FDR 0.05) identified by ClusterDE and standard post-clustering DE tests for each subtype pair. Numbers on bars indicate the DE gene count (black) and proportion among all genes (white). Totals below the x-axis summarize the union of DE genes across all three pairwise comparisons. **k**, Heatmaps of the top DE genes between Classical and Mesenchymal subtypes identified by ClusterDE with the Wilcoxon test (left), which constitute a subset of those identified by the standard Wilcoxon test, and of the top DE genes identified only by the standard Wilcoxon test (right). **l**, PCA visualization of all GBM samples colored by known subtype labels (left), with clustering based on DE genes selected by ClusterDE with Wilcoxon test (middle) and by the standard Wilcoxon test (right). **m-r**, ClusterDE controls false positives and improves specificity in post-clustering differential abundance (DA) analysis of microbiome data. **m**, For the thyroid cancer dataset without subgroups, synthetic null data generated by ClusterDE closely resembled the target data in UMAP structure (left), per-genus mean and variance (middle), and genus–genus rank correlations (right). **n**, Bar plot comparing the number and percentage (black and white text, respectively) of DA genera identified by standard post-clustering DA methods versus ClusterDE, which did not identify any DA genera. **o**, Heatmap showing centered log-ratio (CLR) normalized data for the DA genera identified by standard methods. **p**, For the lung cancer dataset with multiple subgroups, synthetic null data resembled the target data in per-genus mean and variance and overall rank correlations, but formed a homogeneous group—eliminating subgroup structure. **q**, Bar plot comparing the number and percentage of DA genera found by standard methods versus ClusterDE. **r**, Heatmap showing CLR-normalized data for DA genera identified by ClusterDE (left), a subset of those found by ANCOMBC2, and DA genera found by ANCOMBC2 only (right).

In summary, these two case studies illustrate a practical two-step strategy for single-cell multi-omics analysis: first use ClusterDE to validate RNA-based clusters, then proceed with downstream DA analysis only for clusters confirmed to be biologically distinct (see “Single-cell multi-omics analysis guidelines” in Section S1.2.6 of the Supplementary Methods).

#### Population-scale bulk transcriptomics

Clustering followed by DE analysis is common in bulk transcriptomic studies, where patient samples are grouped to characterize subtypes and associated marker genes [30, 31, 32]. As in single-cell analyses, this practice can yield clustering-induced false positives. We applied ClusterDE to two population-scale datasets previously analyzed using post-clustering DE.

The first dataset is a glioblastoma (GBM) microarray study of 147 samples comprising three established subtypes: Proneural, Classical, and Mesenchymal [30]. To construct a synthetic null dataset appropriate for bulk microarray data, we modeled the joint gene distribution using a multivariate Gaussian (Section **??**). The synthetic null data closely resembled the target data and filled the gaps between subtypes (Fig. 7g,i; Fig. S34), representing a homogeneous population under the null. That is, the null-generation step of ClusterDE is data type–adaptive.

To assess over-clustering, we first analyzed samples within each subtype, where no further structure is expected. Following the original study’s clustering approach, hierarchical clustering of samples within a subtype produced two clusters. Standard post-clustering DE methods (limma, Wilcoxon, t-test) detected hundreds of DE genes (i.e., false-positive cluster markers), whereas ClusterDE identified at most one gene in any subtype (Fig. 7h), showing that ClusterDE suppresses false discoveries in a homogeneous population.

Having shown that ClusterDE correctly avoids false-positive marker identification within a homogeneous subtype, we next evaluated whether ClusterDE retains statistical power when true subtypes exist, using the three established GBM subtypes. The DE genes identified by ClusterDE displayed stronger subtype-specific contrasts than those from standard post-clustering DE methods alone (Fig. 7j,k). Clustering on these ClusterDE-selected DE genes more accurately recovered the known subtypes (Fig. 7l), confirming that ClusterDE improves marker specificity when true subtypes exist.

The second dataset is from a TCGA ovarian cancer study [31], where expression of interferon-stimulated genes (ISGs) was used to define two subgroups via consensus K-means clustering followed by DE analysis. Because this dataset is RNA-seq based, we generated a synthetic null dataset using gene-wise negative binomial modeling rather than the Gaussian model used for microarrays. The synthetic null resembled the target data (Fig. S35a), demonstrating the flexibility of ClusterDE’s null-generation strategy. Standard post-clustering DE analyses using DESeq2 and edgeR identified 198 and 1,163 DE genes, respectively (Fig. S35b). In contrast, ClusterDE identified zero DE genes for both methods, raising concerns about the validity of the reported subgroups. Supporting evidence includes minimal expression contrast (Fig. S35c), GO terms unrelated to ovarian cancer (Fig. S35d), and poor subgroup separation in a UMAP embedding (Fig. S35a, top left). These results show that ClusterDE prevents false validation of spurious clusters in bulk transcriptomic studies.

Together, these two case studies illustrate a practical strategy for population-scale bulk transcriptomic analysis: select an appropriate null model based on data type (e.g., a multivariate Gaussian model for microarray data and a multivariate negative binomial model via Gaussian copula for RNA-seq), then use ClusterDE to validate candidate clusters before treating them as true subtypes and reporting their marker genes (see “Population-scale bulk transcriptomics analysis guidelines” in Section S1.2.7 of the Supplementary Methods).

#### Microbiome data

Microbiome studies commonly employ clustering followed by differential abundance (DA) analysis to identify microbial signatures associated with host clusters [66, 67]. This workflow is also susceptible to false positives induced by over-clustering and double dipping. We applied ClusterDE to three microbiome datasets generated using whole-genome sequencing (WGS) and 16S rRNA sequencing.

The first dataset comprises WGS microbiome profiles from thyroid cancer samples in TCGA [68], where no cancer subtypes are expected. To adapt ClusterDE, we developed a synthetic null generator for microbiome data by modeling each genus with a negative binomial distribution including log sequencing depth as a covariate. The synthetic null data preserved key distributional features of the target data (Fig. 7m). Between two clusters obtained via K-means clustering, ClusterDE consistently detected zero differentially abundant (DA) genera, whereas standard DA methods (ANCOMBC2, Wilcoxon, MaAsLin2) identified 3–10 DA genera (Fig. 7n), despite a lack of clear abundance differences (Fig. 7o) or cluster separation (Fig. 7m). These results indicate that the two clusters are spurious and that ClusterDE correctly suppresses false-positive DA calls.

Having shown that ClusterDE suppresses false-positive DA calls in a setting where no subtypes are expected, we next asked whether ClusterDE retains statistical power when true subtypes exist. The second dataset consists of lung cancer WGS microbiome profiles previously shown to contain two subtypes [69]. With the same null-generation strategy, the synthetic null data resembled a homogeneous population (Fig. 7p). ClusterDE identified a subset of DA genera reported by ANCOMBC2, and these genera showed clear abundance contrasts between clusters (Fig. 7r, left). In contrast, many genera identified only by ANCOMBC2 showed weak or noisy differences (Fig. 7r, right). Thus, ClusterDE retains biologically meaningful markers while filtering out likely false positives.

The third dataset is a 16S rRNA vaginal microbiome dataset [70]. The vaginal microbiome comprises five community state types, typically distinguished by the abundance of *Lactobacillus*. We focused on two clusters corresponding to community state types I and IV. Synthetic null data preserved key characteristics of the original data while filling gaps between clusters (Fig. S36a). DA analysis using ClusterDE produced a more selective set of ASVs with strong abundance contrasts, forming a subset of those identified by ANCOMBC2 alone (Fig. S36b), demonstrating improved marker specificity even at fine taxonomic granularity.

Together, these three case studies illustrate a practical strategy for microbiome data analysis: construct a synthetic null using a multivariate negative binomial model via Gaussian copula, modeling each taxon (e.g., genus or ASV) with sequencing depth as a covariate, then use ClusterDE to distinguish spurious over-clustering-induced DA taxa from microbial markers of distinct clusters for both WGS and 16S datasets (see “Microbiome analysis guidelines” in Section S1.2.8 of the Supplementary Methods).

In summary, across these diverse data types, ClusterDE consistently removed false discoveries arising from over-clustering and clustering-induced bias, yielding more reliable and specific markers. These results illustrate that ClusterDE operates as a general calibration framework for post-clustering inference, applicable across omics modalities and robust to variations in data structure and measurement technology.

## 3 Discussion

ClusterDE provides a principled solution to a long-standing challenge in post-clustering differential analysis: controlling false discoveries that arise when the same data are used both to form clusters and to test for differential expression or abundance. It also addresses a related but distinct problem in multi-omics settings, where clustering on one modality (e.g., RNA) induces spurious differential signals in a correlated second modality (e.g., ATAC), even without literal double dipping. Across scRNA-seq, SRT, single-cell multi-omics, population-scale bulk transcriptomics, and microbiome profiling, ClusterDE consistently suppresses over-clustering-induced false positives while recovering biologically meaningful markers when genuine cluster structure exists.

A practical consideration is scalability. The computational cost of ClusterDE depends primarily on the generation of synthetic null data. In our experiments, null-data generation completed within practical runtimes (Fig. S37), and because the procedure is fully parallelizable, substantial speedups can be achieved with additional CPUs (Fig. S37). To further improve practicality, we developed a fast approximation algorithm that delivers a fourfold reduction in runtime with minimal performance loss (Fig. S38), thereby making ClusterDE readily applicable to large single-cell, spatial, and population-scale datasets.

Another consideration is statistical power. Because the synthetic null data represents a homogeneous population (e.g., a single cell type, spatial domain, patient population, or microbiome community), ClusterDE intentionally emphasizes specificity (i.e., suppressing false positives) when evaluating whether observed expression or abundance differences reflect genuine biological separation. This conservatism is essential for avoiding over-clustering–induced false discoveries across diverse omics modalities. In settings where two distinct biological populations are present, this emphasis on specificity may reduce sensitivity to weaker markers. However, this is generally less concerning given the high level of noise inherent in omics data: for annotation tasks such as defining cell types, spatial domains, or subpopulations, only a modest number of informative markers are typically required—often on the order of 10–20 strong features [71, 72]. In practice, ClusterDE reliably identifies these high-specificity markers even under conservative null assumptions. Developing strategies that increase sensitivity while preserving strict control of false discoveries represents an important direction for future research.

ClusterDE is highly adaptable. It operates upstream of any differential testing method and downstream of any clustering algorithm, and it supports data-specific null-generation strategies beyond those used for single-cell and spatial transcriptomics. This flexibility enables its application to diverse omics modalities. Moreover, we extended ClusterDE to handle more complex structures, including hierarchical cell types (ClusterDE-Tree) and continuous cell trajectories, allowing the framework to generalize beyond simple pairwise cluster comparisons and to address clustering-induced bias in common downstream single-cell workflows. Applied to an atlas-scale taxonomy of the human brain, ClusterDE-Tree refined over-resolved cell-type annotations while preserving cell types whose distinctions were supported by independent SRT data, demonstrating that this hierarchical extension scales to real-world cell atlas construction. To facilitate practical adoption, we provide concrete, modality-specific guidelines in the Supplementary Methods for applying ClusterDE to single-cell RNA-seq (including cluster hierarchies and trajectories), SRT, single-cell multi-omics, population-scale bulk transcriptomics, and microbiome data.

Finally, the conceptual foundation of ClusterDE—the construction of synthetic null data as an *in silico* negative control—has broad implications beyond the specific applications explored here. Clustering-induced bias, including but not limited to literal double dipping, is pervasive across single-cell, spatial, integrative multi-omic, bulk transcriptomic, and microbiome analyses. It also arises in downstream workflows such as post-pseudotime differential testing [73], integration-based comparisons, and other procedures in which data-driven structure is inferred and then reused for statistical testing. Synthetic null data and contrast-based testing provide a general strategy for calibrating these analyses, yielding more reliable inferences even in complex, high-dimensional settings. We anticipate that this framework will motivate new methodological developments aimed at mitigating prior-step-induced biases and reducing false discoveries across a wide range of high-throughput sequencing applications.

## Code and Data Availability

The R package ClusterDE is available at https://github.com/SONGDONGYUAN1994/ClusterDE. The tutorials of ClusterDE are available at https://songdongyuan1994.github.io/ClusterDE/index.html. The source code and data for reproducing the results are available at: http://doi.org/10.5281/zenodo.8161964 [74].

## Acknowledgements

The authors appreciate the comments and feedback from the Junction of Statistics and Biology members (https://jsb-lab.org). This material is based upon work supported by the National Science Foundation Graduate Research Fellowship Program under Grant No.DGE-2034835. Any opinions, findings, and conclusions or recommendations expressed in this material are those of the author(s) and do not necessarily reflect the views of the National Science Foundation.

