## Supplementary for "Synthetic Control Enables Reliable Cluster Validation and Marker Discovery in Omics Data: From Single-Cell and Spatial to Population-Scale"

#### Table of contents

– S1. Supplementary Methods
- S1.1. Overview of ClusterDE variants
- S1.2. Practical guidelines for using ClusterDE
  - S1.2.1. Usage for cell-cluster marker identification
  - S1.2.2. Usage for cluster merging decision
  - S1.2.3. Practical guidelines for applying ClusterDE-Tree to cluster hierarchies and atlas-scale taxonomies
  - S1.2.4. Practical guidelines for applying ClusterDE to trajectory-based analyses
  - S1.2.5. Practical guidelines for applying ClusterDE to SRT data
  - S1.2.6. Single-cell multi-omics analysis guidelines
  - S1.2.7. Population-scale bulk transcriptomics analysis guidelines
  - S1.2.8. Microbiome analysis guidelines
- S1.3. ClusterDE-Core framework and four-step procedure
  - S1.3.1. Notations for the double-dipping problem in post-clustering DE analysis
  - S1.3.2. ClusterDE Step 1: synthetic null data generation
  - S1.3.3. ClusterDE Step 2: cell clustering
  - S1.3.4. ClusterDE Step 3: DE analysis
  - S1.3.5. ClusterDE Step 4: FDR control
- S1.4. ClusterDE extensions
  - S1.4.1. ClusterDE-Stable details
  - S1.4.2. ClusterDE-Tree details
  - S1.4.3. ClusterDE-Fast details
  - S1.4.4. Synthetic null data quality checks and simulator comparison
  - S1.4.5. ClusterDE trajectory-based DE
- S1.5. scRNA-seq simulation studies and evaluations
  - S1.5.1. scRNA-seq simulation designs
  - S1.5.2. Simulation setting with one cell type
  - S1.5.3. Simulation setting with two cell types and 200 true DE genes
  - S1.5.4. Implementation of the TN test and Countsplit
  - S1.5.5. Alternative strategies for synthetic null data generation
  - S1.5.6. Validity checks of contrast scores and $P$ values
  - S1.5.7. Random-seed stability and robustness to null-parameter perturbation
- S1.6. scRNA-seq real-data analyses
  - S1.6.1. Pure cell-line datasets
  - S1.6.2. $CD14^+$/$CD16^+$ monocyte datasets
  - S1.6.3. Dimensionality reduction and visualization
  - S1.6.4. Gene set enrichment analysis for monocyte markers and housekeeping genes
  - S1.6.5. Adult *Drosophila* neuronal visual atlas
  - S1.6.6. Retracted human fetal cerebellum TCP dataset
  - S1.6.7. COVID-19 PBMC dataset
  - S1.6.8. PBMC $CD4^+$ and cytotoxic T-cell subsets

    * S1.6.9. H2228 cell line and human glutamatergic neurogenesis datasets
    * S1.6.10. Mouse bone marrow dataset
    * S1.6.11. Allen Human MTG atlas analysis
  • S1.7. Spatially resolved transcriptomics (SRT) data analyses
    * S1.7.1. SRT simulation designs
    * S1.7.2. Simulation setting with one spatial domain
    * S1.7.3. Simulation setting with two spatial domains and 200 true spatial-domain marker genes
    * S1.7.4. Simulation setting with three spatial domains and 200 and 168 true spatial-domain marker genes
    * S1.7.5. DLPFC and PDAC SRT datasets
  • S1.8. Analyses across other omics modalities
    * S1.8.1. Single-cell multiome analysis
    * S1.8.2. Population-scale bulk transcriptomics analysis
    * S1.8.3. Microbiome analysis
– S2. Theoretical justification of ClusterDE
  • S2.1. Assumptions for FDR control of ClusterDE
  • S2.2. Theoretical justification for preserving gene–gene correlation in synthetic null data
– S4. Supplementary Tables
– S3. Supplementary Figures
– S5. Supplementary Algorithms
– References

### S1 Supplementary Methods

#### S1.1 Overview of ClusterDE variants

Before presenting detailed usage guidelines, we briefly summarize the four ClusterDE variants introduced throughout this Supplementary Material and their relationships:

- **ClusterDE-Core** is the base procedure. It generates a single synthetic null dataset and performs marker gene detection at a target FDR (Section S1.3).
- **ClusterDE-Fast** is a computationally efficient alternative to ClusterDE-Core. It replaces the synthetic null data generation step with a low-rank (PCA-based) approximation, substantially reducing runtime while preserving the same downstream marker-detection logic (Section S1.4.3).
- **ClusterDE-Stable** extends ClusterDE-Core (or ClusterDE-Fast, for large-scale data) by repeatedly generating $B$ synthetic null datasets and aggregating the results for two purposes: (1) testing whether two clusters are separated with statistical significance, to inform merging decisions, and (2) identifying stable marker genes via stability selection across the $B$ runs (Section S1.4.1).
- **ClusterDE-Tree** recursively applies the ClusterDE-Stable cluster-separation test to an entire cluster hierarchy, moving bottom-up from the leaves to prune statistically unsupported splits, and then applies ClusterDE-Stable's marker-selection procedure to identify stable markers for the retained clusters (Section S1.4.2).

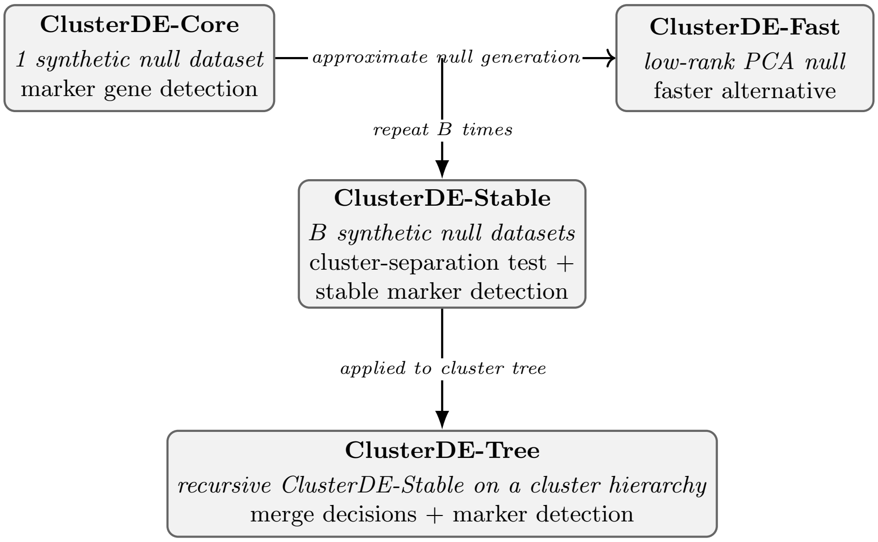


#### S1.2 Practical guidelines for using ClusterDE

**S1.2.1 Usage for cell-cluster marker identification** ClusterDE is designed to identify reliable cell-cluster marker genes through pairwise comparisons of potentially ambiguous clusters obtained from cell clustering. For single-cell RNA-seq (scRNA-seq) data, distinct cell types or subtypes are expected to have marker genes with clearly separated expression patterns, often exhibiting bimodal expression distributions across all cells, indicating the presence of two cell subpopulations.

We recommend the following workflow for identifying cell-cluster marker genes from scRNA-seq data:

1. From cells already clustered, select two clusters of interest to compare (whether their separation is ambiguous or not). For scRNA-seq data analyzed with Seurat, the `BuildClusterTree` function can be used to organize clusters hierarchically and identify sibling clusters for comparison. See Section S1.2.3, "Practical guidelines for applying ClusterDE-Tree to cluster hierarchies and atlas-scale taxonomies."
2. Subset the data to include only the cells belonging to the two selected clusters.
3. Use the subsetted data as the "target data" input for ClusterDE. Specify a target FDR.

4. Perform diagnostics on the quality of synthetic null data generated by ClusterDE. The synthetic null data should contain synthetic cells that preserve key properties of the target data—such as feature means, variances, correlations, and embeddings—while representing a single homogeneous population (i.e., without the separation between the two original clusters, if any).
5. Review the marker genes identified by ClusterDE and assess whether their expression patterns and biological functions support the interpretation of the two clusters as distinct cell types or subtypes.

The above workflow generates one synthetic null dataset and is considered the core ClusterDE procedure, referred to as **ClusterDE-Core** (detailed in Section S1.3 "ClusterDE-Core framework and four-step procedure" and Algorithm S1). It is the default mode of ClusterDE used throughout the main text, unless otherwise noted.

If computational resources allow and users want to find more stable marker genes across multiple synthetic null datasets, users can use **ClusterDE-Stable**, which repeats ClusterDE-Core multiple times, each time using an independently generated synthetic null dataset, and finally aggregates identified genes across runs. ClusterDE-Stable reports a gene as a stable marker when it is selected in at least half of the runs at the target FDR. Details of this procedure are given under "ClusterDE-Stable details" in Section S1.4.1.

We note that ClusterDE can also be used to compare two clusters that are already confidently distinct, rather than only ambiguous ones. In this setting, ClusterDE typically reports fewer marker genes than standard post-clustering DE tests, but these genes tend to be more biologically specific to each cluster. We expect that the higher specificity of ClusterDE's marker genes makes them more useful for annotating clusters based on literature-curated markers.

Users may alternatively allow ClusterDE to re-cluster the target data if they suspect that the original two-cluster partition does not reflect the true separation between two cell subpopulations (rather than suspecting that only one true population exists). In this case, the resulting two clusters from re-clustering may differ from the original input clusters. This option is therefore appropriate for evaluating the newly formed clusters but is not recommended when the goal is to assess whether the two originally given clusters represent distinct biological populations.

ClusterDE focuses on identifying reliable post-clustering marker genes, but if users are interested in knowing whether two clusters should be merged first, they should refer to Section S1.2.2, "Usage for cluster merging decision."

**S1.2.2 Usage for cluster merging decision** If users are interested in deciding whether two clusters should be merged using a statistical test, we recommend the following workflow:

1. Consider two clusters obtained from cell clustering.
2. Calculate their observed normalized Ward linkage statistic, which measures the separation between the two clusters in the selected low-dimensional representation. The formal definition of the normalized Ward linkage statistic is given in the "Stable cluster-separation assessment" paragraph of "ClusterDE-Stable details" in Section S1.4.1.
3. Generate multiple homogeneous synthetic null datasets and calculate the same statistic for each null dataset.
4. Perform diagnostics on the quality of the synthetic null datasets, following the same checks recommended in Section S1.2.1, "Usage for cell-cluster marker identification."
5. Compute the "split $P$-value" as the proportion of synthetic null datasets whose normalized Ward linkage statistic is at least as large as the observed statistic, following the same calculation detailed in Section S1.4.1. A sufficient number of synthetic null datasets should be generated to obtain a stable empirical $P$-value.
6. Retain the two clusters when the split $P$-value is below 0.05. Otherwise, consider the split to be not statistically significant, and merge the clusters.

Marker-gene results from Section S1.2.1 provide complementary biological evidence (for example, for two ambiguous clusters, we expect to find zero or few marker genes), but are not the formal cluster-level decision criterion. In particular, the absence of ClusterDE marker genes may support merging but should not replace the split-significance test.

This split-significance procedure applies to a single pair of clusters and is formally defined as part of **ClusterDE-Stable** (Section S1.4.1). When the goal is to build or refine an entire cluster hierarchy—for example, a cell atlas organized in a hierarchy—the same test is applied recursively from the leaf clusters of the hierarchy, and the procedure is referred to as **ClusterDE-Tree**; see Section S1.2.3, "Practical guidelines for applying ClusterDE-Tree to cluster hierarchies and atlas-scale taxonomies." Details of the ClusterDE-Tree recursive procedure are given under "ClusterDE-Tree details" in Section S1.4.2.

**S1.2.3 Practical guidelines for applying ClusterDE-Tree to cluster hierarchies and atlas-scale taxonomies** ClusterDE-Tree extends the pairwise merging-decision procedure in Section S1.2.2 to an entire cluster hierarchy, such as an atlas-scale taxonomy, by applying the ClusterDE-Stable cluster-separation test (Section S1.4.1) recursively to internal nodes, moving from the leaves upward and stopping along each branch once a non-merging decision is reached. We recommend the following workflow:

1. Construct a binary cluster tree using a standard clustering pipeline (e.g., Seurat's `BuildClusterTree` for scRNA-seq atlases), following the tree-building method described in "ClusterDE-Tree details" in Section S1.4.2.
2. Starting from the leaves and moving upward toward the root, evaluate each internal node (i.e., each sibling split) using the same workflow described in Section S1.2.2, "Usage for cluster merging decision"—including diagnosing synthetic null quality before interpreting the split $P$-value. Prune splits whose $P$-values are at or above the chosen threshold (typically 0.05) by merging the corresponding child nodes, and retain splits whose $P$-values fall below the threshold. Continue this bottom-up procedure along each branch until a split is retained; testing does not continue further up that branch once its children are not merged.
3. After pruning, treat each remaining leaf as a refined cluster. For each refined cluster, identify its markers by comparing it against its sister clade (the other subtree connected to the same parent node) using ClusterDE-Stable, as detailed in Section S1.4.1. Report genes selected in at least half of the synthetic null datasets as stable markers, at the target FDR.
4. **(Optional but desirable if possible)** For atlas-scale taxonomies (e.g., the Allen MTG snRNA-seq atlas), interpret refined clusters and their markers by mapping them to external datasets (such as SRT) when available, as illustrated in the main text. This provides biological validation of refined cluster boundaries and markers.

For atlas-scale taxonomies with a large number of internal nodes and cells, generating synthetic null data in Steps 2 and 3 above becomes computationally demanding using the ClusterDE-Core null generation. We recommend substituting the synthetic null data generation step with **ClusterDE-Fast**, which approximates the synthetic null distribution via a low-rank (PCA-based) representation of the data, as detailed in "ClusterDE-Fast details" in Section S1.4.3. ClusterDE-Fast substantially reduces the computational cost of the recursive tree-pruning and marker-identification steps while maintaining stable marker genes detected by ClusterDE-Stable compared with using ClusterDE-Core to generate synthetic null data.

These practical guidelines implement the ClusterDE-Tree extension detailed in Section S1.4.2, which recursively applies the ClusterDE-Stable cluster-separation test and ClusterDE-Stable-based marker identification (Section S1.4.1) across a cell-cluster hierarchy. For large-scale taxonomies, ClusterDE-Fast (Section S1.4.3) can be substituted for the synthetic null data generation step to improve scalability.

**S1.2.4 Practical guidelines for applying ClusterDE to trajectory-based analyses** ClusterDE can be adapted to trajectory-based analyses by treating trajectory-defined lineages as groups that may or may not represent true bifurcations. We recommend the following workflow, consistent with the "ClusterDE trajectory-based DE" subsection in Section S1.4.5:

1. Infer trajectories using a standard trajectory-inference algorithm (e.g., Slingshot), following the original study's preprocessing and dimensionality reduction steps. Designate a root cluster and terminal clusters, and identify candidate lineage pairs (e.g., two branches emerging from a bifurcation) where DE analysis is of interest.

2. For each lineage pair, restrict attention to non-root cells confidently assigned to exactly one lineage (e.g., cells with lineage weight 1.0 for one lineage and 0 for all others). Perform DE testing between the two lineages using a suitable test (e.g., Wilcoxon rank-sum on log-normalized expression) to obtain target $P$-values and corresponding target DE scores, as described in "ClusterDE trajectory-based DE" (Section S1.4.5).
3. Generate synthetic null data representing a homogeneous single-lineage scenario by applying the ClusterDE null-generation procedure (ClusterDE-Core or ClusterDE-Fast, as described in "ClusterDE Step 1: synthetic null data generation," Section S1.3.2) to the non-root cells, and combining synthetic counts with the original root-cell counts. Project synthetic null cells into the same low-dimensional space as the target data, apply the same clustering procedure, and re-run trajectory inference (e.g., Slingshot) to obtain null lineage assignments and null DE $P$-values for the same lineage pair.
4. Construct contrast scores by transforming both target and null $P$-values to DE scores (e.g., via $-\log_{10}$) and subtracting null scores from target scores, as in "ClusterDE trajectory-based DE" and "ClusterDE Step 4: FDR control" (Section S1.3.5). Apply the Clipper-based FDR control procedure to these contrast scores to identify lineage-specific DE genes at the desired FDR level.
5. Interpret results as follows: in datasets where no true bifurcation is expected (negative controls), if ClusterDE reports zero DE genes between trajectory-defined groups, treat the inferred split as an artifact of over-clustering and do not interpret the trajectory markers as biologically meaningful. In datasets with known bifurcations (e.g., common myeloid progenitors splitting into granulocyte–monocyte progenitors and erythroid precursors), prioritize genes with large positive contrast scores as lineage-specific markers, and use them to validate the inferred trajectory structure.

**S1.2.5 Practical guidelines for applying ClusterDE to SRT data** For SRT data, ClusterDE is used to distinguish genuine spatial domains from spurious clusters and to identify spatial-domain marker genes with discontinuous expression boundaries. We recommend the following workflow:

1. Perform spatial clustering (e.g., using BayesSpace) to obtain spatial clusters that represent putative domains, following the clustering procedure described in "ClusterDE Step 2: cell clustering" for SRT data (Section S1.3.3). These clusters serve as candidate spatial domains or subdomains for subsequent DE analysis.
2. Select a pair of adjacent domains to identify domain markers. Subset the data to the spots in the two clusters being compared, and treat this subset as the target data input to ClusterDE.
3. Generate synthetic null SRT data representing a spatially homogeneous scenario by applying the spatial NB–Gaussian-copula null model described in "ClusterDE Step 1: synthetic null data generation" (Section S1.3.2) to the subset of spots. In this null model, each gene's mean expression is modeled as a smooth surface over spatial locations, so that sharp domain boundaries are absent by construction. Use synthetic null data diagnostics (e.g., per-gene means/variances, gene–gene correlations, embedding-based mixing metrics) to confirm smooth spatial variation and the absence of discontinuous domain boundaries.
4. Apply the same spatial clustering and DE workflow (e.g., BayesSpace plus Wilcoxon or t-test on log-normalized expression) to both target and synthetic null data, and then construct contrast scores by transforming target and null $P$-values to DE scores (e.g., via $-\log_{10}$) and subtracting null scores from target scores, as in "ClusterDE Step 4: FDR control" (Section S1.3.5). Use ClusterDE's Clipper-based contrast-score thresholding to identify spatial-domain markers at the desired FDR level.
5. Interpret results as follows: within a homogeneous annotated domain, ClusterDE should report nearly zero spatial-domain markers between subclusters, supporting the merging of over-clustered regions and indicating that boundaries are artifacts of over-clustering rather than true domain differences. Between adjacent domains, bona fide ClusterDE markers should exhibit discontinuous spatial boundaries and strongly domain-specific expression patterns, whereas genes with smooth spatial gradients should not be identified as spatial-domain markers.

The ClusterDE-Core workflow for SRT, including the spatial NB–Gaussian-copula null model, BayesSpace-based clustering, and contrast-score FDR control, is summarized in Section S1.3 and Algorithm S2. Extensions such as ClusterDE-Stable and synthetic null quality checks are described in Section S1.4.

**S1.2.6 Single-cell multi-omics analysis guidelines** For single-cell multiome assays (joint RNA-seq and ATAC-seq), clustering is typically performed in the RNA modality and differential accessibility analysis is conducted in the ATAC modality. We recommend the following workflow, as described in the main text:

1. Apply ClusterDE to RNA-defined clusters to determine whether the split between clusters is statistically supported and biologically meaningful, using the split-significance workflow described in Section S1.2.2. If the conclusion is that the two RNA clusters are ambiguous, treat the split as spurious and merge the clusters before any ATAC analysis.
2. For RNA-defined splits that are statistically supported (split $P$-value below 0.05), use the marker-selection workflow described in Section S1.2.1 with ClusterDE-Stable (Section S1.4.1) to identify stable RNA markers, reporting genes selected in at least half of synthetic null datasets at the target FDR.
3. Retain the same cell clusters, and apply ClusterDE-Stable to the ATAC modality, using the null ATAC model described in “ClusterDE Step 1: synthetic null data generation” and the clustering/DE tests described in “ClusterDE Step 2” and “ClusterDE Step 3” (Section S1.3). Report peaks selected in at least half of synthetic null datasets as stable differentially accessible markers.
4. When RNA-defined cell clusters are not statistically supported, do not interpret ATAC differences as evidence of distinct cell subpopulations. When both RNA and ATAC markers show clear between-cluster contrasts, treat these genes and peaks as cluster-specific multi-omic markers.

The ClusterDE workflow for single-cell multiome data (paired RNA and ATAC modalities) is described in Algorithm S3.

**S1.2.7 Population-scale bulk transcriptomics analysis guidelines** For bulk microarray and bulk RNA-seq studies, clustering followed by DE analysis is a common strategy for defining patient subtypes and associated markers, but is susceptible to clustering-induced false positives. We recommend the following workflow:

1. Choose a clustering method (e.g., hierarchical clustering or K-means) consistent with the original study, and construct clusters corresponding to putative subtypes, using the clustering procedure described in “ClusterDE Step 2: cell clustering” (Section S1.3.3).
2. For microarray data, fit the multivariate Gaussian null model described in “ClusterDE Step 1: synthetic null data generation”; for RNA-seq data, fit the multivariate negative binomial model via Gaussian copula described in the same subsection (Section S1.3.2). Generate multiple synthetic null datasets representing a homogeneous population with similar global expression structure.
3. Apply the split-significance procedure from Section S1.2.2 within established subtypes (where no further structure is expected) and across putative subtypes. For clusters within a homogeneous subtype, ClusterDE should suggest a cluster merge and identify at most a negligible number of DE genes, signaling spurious within-subtype splits. Between two established subtypes, ClusterDE should suggest the split to retain.
4. For each retained split, apply ClusterDE (or ClusterDE-Stable) with appropriate bulk DE tests (limma, t-test, DESeq2, edgeR) as described in “ClusterDE Step 3: DE analysis” (Section S1.3.4) to identify subtype markers. The identified markers are expected to show clear subtype-specific expression contrasts.

The bulk transcriptomic version of ClusterDE-Core, covering both microarray and RNA-seq null models and clustering strategies, is presented in Section S1.3 and Algorithm S4. ClusterDE-Stable, which applies stability selection on top of this core procedure, is described in Section S1.4.

**S1.2.8 Microbiome analysis guidelines** For microbiome data (whole-genome sequencing or 16S), clustering followed by differential abundance (DA) analysis is widely used to identify microbial signatures associated with host clusters, but can be affected by over-clustering and confounded by sequencing depth. We recommend the following workflow:

1. Construct host-sample clusters using the same distance or ordination strategy as in the original study (e.g., K-means on compositional principal components or hierarchical clustering on Bray–Curtis distances), following the clustering procedures summarized in “ClusterDE Step 2: cell clustering” (Section S1.3.3).

2. Fit a microbiome-specific null model using a multivariate negative binomial distribution via Gaussian copula, with genus- or ASV-level marginals including log sequencing depth as a covariate, as described in "ClusterDE Step 1: synthetic null data generation" (Section S1.3.2). Generate multiple synthetic null datasets representing homogeneous microbial samples.
3. Apply the split-significance procedure from Section S1.2.2 to candidate cluster splits. When ClusterDE suggests cluster merging, treat the inferred clusters as spurious and avoid interpreting DA results as biological differences.
4. For statistically supported splits, use ClusterDE (or ClusterDE-Stable) with established DA methods (ANCOMBC2, Wilcoxon rank-sum test, MaAsLin2) as described in "ClusterDE Step 3: DE analysis" (Section S1.3.4) to identify stable differentially abundant taxa. The identified genera or ASVs are expected to show clear between-cluster abundance contrasts.

Algorithm S5 provides the ClusterDE-Core workflow for microbiome data, based on depth-adjusted NB marginals and a Gaussian copula for synthetic null data generation, as a specification of the ClusterDE-Core framework in Section S1.3. ClusterDE-Stable, which applies stability selection on top of this core procedure, is described in Section S1.4.

#### S1.3 ClusterDE-Core framework and four-step procedure

**S1.3.1 Notations for the double-dipping problem in post-clustering DE analysis.** The target data is represented as $\mathbf{Y} = [Y_{ij}] \in \mathbb{N}_{\geq 0}^{n \times m}$, where $n$ observations are rows, $m$ genes are columns, and $Y_{ij}$ denotes the UMI count of gene $j = 1, \ldots, m$ in observation $i = 1, \ldots, n$. In scRNA-seq data, each observation corresponds to a cell, while in SRT data, observations can vary in spatial resolution—for instance, each spot may contain a single cell or multiple cells. For simplicity, we may refer to spots as cells. Each observation $i$ is represented as an $m$-dimensional gene expression vector, $\mathbf{Y}_i = (Y_{i1}, \ldots, Y_{im})^{\mathsf{T}}$.

In our formulation of the post-clustering DE problem for scRNA-seq data, the $n$ cells belong to two latent cell types, which may or may not be identical, and are partitioned into two clusters by a clustering algorithm. We represent cell $i$'s latent cell type using $Z_i \in \{0, 1\}$. We define the "ideal DE test" as the test that decides whether a gene has equal mean expression in two cell types. For gene $j$, we assume that its expression levels in cells of type 0, $\{(Y_{ij}|Z_i = 0)\}_{i=1}^n$, share a common mean, denoted as $\mu_{0j} = \mathbb{E}[Y_{ij}|Z_i = 0]$, and its expression levels in cells of type 1, $\{(Y_{ij}|Z_i = 1)\}_{i=1}^n$, share a common mean, denoted as $\mu_{1j} = \mathbb{E}[Y_{ij}|Z_i = 1]$. The ideal DE test is then formulated with the following null hypothesis $H_{0j}$ and alternative hypothesis $H_{1j}$:

$$H_{0j} : \mu_{0j} = \mu_{1j} \quad \text{vs.} \quad H_{1j} : \mu_{0j} \neq \mu_{1j} \, .$$

Hence, gene $j$ is a true DE gene if and only if $H_{0j}$ does not hold. If all $n$ cells belong to a single cell type (i.e., the two latent cell types are identical), then all $m$ null hypotheses, $H_{01}, \ldots, H_{0m}$, hold simultaneously.

Note that the null hypothesis depends on the DE test used. In most cases, the DE test evaluates the equality of means. However, the default DE test in Seurat, the Wilcoxon rank-sum test, assesses whether a gene's expression distributions are identical between the two latent cell types. Nonetheless, the above 'equal mean' formulation can be easily generalized.

Since $Z_i$'s are unobserved, standard scRNA-seq data analysis partitions cells into two clusters using a clustering algorithm $g$ (e.g., the Louvain algorithm in Seurat) applied to $\mathbf{Y}$. We denote cell $i$'s cluster membership as $\widehat{Z}_i = g_{\mathbf{Y}}(\mathbf{Y}_i) \in \{0, 1\}$, where $g_{\mathbf{Y}} : \{\mathbf{Y}_1, \ldots, \mathbf{Y}_n\} \to \{0, 1\}$ is the clustering function constructed from the clustering algorithm $g$ and the data $\mathbf{Y}$, mapping a cell's gene expression vector to a cluster membership.

For spot $i$ in SRT data, in addition to the gene expression vector $\mathbf{Y}_i$, we also observe the two-dimensional spatial location $X_i = (x_{i1}, x_{i2})^{\mathsf{T}}$. Typically, spatial clustering algorithms incorporate both gene expression data $\mathbf{Y}$ and spatial locations $X = [X_1, \cdots, X_n]^{\mathsf{T}}$ when clustering spots. Consequently, the cluster membership for spot $i$ is given by $\widehat{Z}_i = g_{\mathbf{Y},X}(\mathbf{Y}_i, x_{i1}, x_{i2}) \in \{0, 1\}$, where $g_{\mathbf{Y},X}$ represents the clustering function utilizing both data types.

After clustering, standard scRNA-seq and SRT analyses conduct a DE test for each gene based on $\mathbf{Y}_1, \ldots, \mathbf{Y}_n$ and $\widehat{Z}_1, \ldots, \widehat{Z}_n$. In this process, the data $\mathbf{Y}$ is used twice—once in clustering and again in DE analysis—a phenomenon referred to as the "double-dipping (DD) issue." The standard post-clustering DE analysis in Seurat has this DD issue and employs a statistical test that differs from the ideal DE test.

Specifically, for gene $j$, we define $\mu_{0j}^{\mathrm{DD}} = \mathbb{E}[Y_{ij}|\widehat{Z}_i = 0]$ and $\mu_{1j}^{\mathrm{DD}} = \mathbb{E}[Y_{ij}|\widehat{Z}_i = 1]$. The post-clustering DE analysis in Seurat corresponds to the following null hypothesis $H_{0j}^{\mathrm{DD}}$ and alternative hypothesis $H_{1j}^{\mathrm{DD}}$:

$$H_{0j}^{\mathrm{DD}} : \mu_{0j}^{\mathrm{DD}} = \mu_{1j}^{\mathrm{DD}} \quad \text{vs.} \quad H_{1j}^{\mathrm{DD}} : \mu_{0j}^{\mathrm{DD}} \neq \mu_{1j}^{\mathrm{DD}} .$$

Hence, gene $j$ would be falsely identified as a cell-type marker gene if $H_{0j}^{\mathrm{DD}}$ is rejected but $H_{0j}$ holds, leading to an inflated FDR in identifying cell-type marker genes. Fig. S39 provides a toy example illustrating this issue; the same problem also arises in post-clustering DE analysis for SRT data.

**S1.3.2 ClusterDE Step 1: synthetic null data generation** Previous studies on scRNA-seq data have shown that, within a single cell type, each gene's UMI counts can be well-modeled by a negative binomial (NB) distribution [1, 2, 3]. Furthermore, the joint UMI counts for all genes can be effectively approximated by a multivariate negative binomial (MVNB) distribution defined using the Gaussian copula [4]. Based on these findings, ClusterDE employs an MVNB distribution specified by the Gaussian copula as the scRNA-seq null model, representing a single "hypothetical" cell type. In step 1 of ClusterDE, the null model is fitted to the target data $\mathbf{Y}$, and synthetic null data are subsequently sampled from the fitted null model. The assumption behind this null model is that a single cell type is a homogeneous population where the marginal count distribution for each gene follows an NB distribution, and the gene dependence structure is specified by the Gaussian copula. Moreover, if users have prior knowledge of a more appropriate marginal distribution, ClusterDE can generate synthetic null data from other models, including multivariate Gaussian, multivariate Poisson, multivariate zero-inflated Poisson, and multivariate zero-inflated NB distributions.

For scATAC-seq data, our null model assumes a homogeneous population within each cell type. For each peak, the marginal count distribution is modeled by the NB distribution, as our practice indicates that a zero-inflated NB model does not provide a better fit for the ATAC modality in the tested multiome data. Users may select the marginal distribution according to the characteristics of their own data. The dependence structure among peaks is modeled using a Gaussian copula, analogous to the RNA setting.

For SRT data, we define a null model representing a single "hypothetical" spatial domain. In this model, each gene exhibits smooth spatially variable expression across the spots, without abrupt changes that would suggest the presence of more than one spatial domain. The gene dependence structure is still modeled using the Gaussian copula.

For bulk RNA-sequencing and microbiome data, our null model assumes a homogeneous population where the expression or abundance of each feature (gene or microbial taxon) follows a NB marginal distribution, and the dependence structure across features is characterized using a Gaussian copula. After estimating the marginal parameters and copula correlation matrix from the data, synthetic null datasets are generated by sampling from the resulting multivariate negative binomial distribution. For bulk microarray data, where normalized expression measurements are continuous and approximately Gaussian, the null model is specified as a multivariate normal distribution with mean vector and covariance matrix estimated from the normalized data. Synthetic null data are then drawn from this fitted Gaussian distribution.

The concept of fitting a null model to the target data, regardless of whether the target data aligns with the null model, underpins the widely used likelihood-ratio test in statistics [5]. In this test, the maximum likelihood under the null hypothesis is calculated from the target data and compared with the maximum likelihood under a more flexible alternative hypothesis, also calculated from the target data. The null hypothesis is rejected only if the maximum likelihood under the null is significantly smaller than that under the alternative. ClusterDE extends this principle by sampling synthetic null data from the null model fitted to the target data using maximum likelihood estimation. This approach allows any post-clustering DE pipeline, regardless of complexity, to be applied simultaneously to both the synthetic null data and the target data. A contrastive strategy is then used to identify trustworthy DE genes—those with DE scores significantly higher in the target data than in the synthetic null data.

1. **Specifying a null model**
   Under the null model for scRNA-seq and scATAC-seq data, we assume that $Y_{ij}$, gene $j$'s UMI count in cell $i$, independently (across cells, not genes) follows the $\mathrm{NB}(\mu_j, \sigma_j)$ distribution with the probability

mass function:

$$\mathbb{P}(Y_{ij} = y; \mu_j, \sigma_j) = \frac{\Gamma\left(y + \frac{1}{\sigma_j}\right)}{\Gamma\left(\frac{1}{\sigma_j}\right)\Gamma(y+1)} \left(\frac{1}{1+\sigma_j\mu_j}\right)^{\frac{1}{\sigma_j}} \left(\frac{\sigma_j\mu_j}{1+\sigma_j\mu_j}\right)^{y};$$

$$y \in \{0, 1, 2, \cdots\},$$

where $\mu_j$ and $\sigma_j$ are the mean and dispersion parameters of the NB distribution for gene $j$. That is,

$$Y_{1j}, \cdots, Y_{nj} \overset{\text{i.i.d.}}{\sim} \text{NB}(\mu_j, \sigma_j),$$

with "i.i.d." short for "independent and identically distributed," meaning that the $n$ cells' counts for gene $j$ represent a random sample from $\text{NB}(\mu_j, \sigma_j)$.

For SRT data, we have additional information: the two-dimensional spatial location of each cell/spot, $X_i = (x_{i1}, x_{i2})^{\mathsf{T}}$. In the null model for SRT data, each gene's mean expression is modeled as a smooth surface over spatial locations. Specifically, we assume that gene $j$'s mean expression at spot $i$ is given by $\log(\mu_{ij}) = \alpha_j + f_j^{GP}(x_{i1}, x_{i2}, K)$, where $\alpha_j$ denotes gene $j$'s specific intercept, and the $f_j^{GP}(\cdot)$ represents a penalized Gaussian process regressor with $K$ bases specific to gene $j$. The parameter $K$ defines the upper bound of the "wiggliness" of the smooth surface. In our default ClusterDE setting, we use $K = 4$, the minimum possible value for $K$, ensuring the smoothest spatial pattern that a two-dimensional Gaussian process regressor can represent. This choice reflects the smoothest spatial variation that aligns with the null hypothesis, avoiding overfitting to gene expression measurement noise. That is,

$$Y_{ij} \overset{\text{ind.}}{\sim} \text{NB}(\mu_{ij}, \sigma_j),$$

where $\mu_{ij} = \alpha_j + f_j^{GP}(x_{i1}, x_{i2}, 4)$. In other words, we assume that under the null model, each gene within a single domain exhibits a smooth spatial expression surface.

For bulk microarray data, the raw probe intensities are continuous, right-skewed, and not well captured by a standard parametric distribution. Therefore, we work with preprocessed expression data (e.g., after background correction, normalization, and log transformation), which are commonly approximated as Gaussian [6]. Specifically, we assume that $Y_{ij}$, the normalized expression level of gene $j$ in sample $i$, independently follows a univariate normal distribution $N(\mu_j, \sigma_j^2)$ across samples. That is,

$$Y_{1j}, \cdots, Y_{nj} \overset{\text{i.i.d.}}{\sim} N(\mu_j, \sigma_j^2).$$

For bulk RNA sequencing (RNA-seq) data, we assume that $Y_{ij}$, the read count of gene $j$ in sample $i$, follows a negative binomial distribution with gene-specific mean $\mu_j$ and dispersion $\sigma_j$, independently across samples (but not across genes). That is

$$Y_{1j}, \cdots, Y_{nj} \overset{\text{i.i.d.}}{\sim} \text{NB}(\mu_j, \sigma_j).$$

For microbiome data, we model each genus's abundance with a negative binomial distribution including log sequencing depth as a covariate. Specifically, we assume $Y_{ij}$, the abundance of genus $j$ in sample $i$, independently follows

$$Y_{ij} \overset{\text{ind.}}{\sim} \text{NB}(\mu_{ij}, \sigma_j)$$

with $\log(\mu_{ij}) = \alpha_j + \beta_j \log(d_i)$, where $d_i$ denotes the sequencing depth of sample $i$.

Let $F_j$ denote the cumulative distribution function (CDF) of $\text{NB}(\mu_j, \sigma_j)$ for scRNA-seq data, scATAC-seq data, and bulk RNA-sequencing data, $\text{NB}(\mu_{ij}, \sigma_j)$ for SRT data and microbiome data, or $N(\mu_j, \sigma_j)$ for bulk microarray data, where $j = 1, \ldots, m$ and $i = 1, \ldots, n$, the MVNB distribution specified by the

Gaussian copula is

$$\left.\begin{matrix}\left(\Phi^{-1}(F_1(Y_{11})),\cdots,\Phi^{-1}(F_m(Y_{1m}))\right)^\top,\\ \vdots \\ \left(\Phi^{-1}(F_1(Y_{n1})),\cdots,\Phi^{-1}(F_m(Y_{nm}))\right)^\top\end{matrix}\right. \overset{\text{i.i.d.}}{\sim} N_m\left(\mathbf{0},\mathbf{R}\right),$$

where $\Phi$ is the CDF of the standard Gaussian distribution $N(0,1)$, and $N_m\left(\mathbf{0},\mathbf{R}\right)$ is an $m$-dimensional Gaussian distribution with an $m$-dimensional $\mathbf{0}$ mean vector and an $m$-by-$m$ correlation matrix $\mathbf{R}$ (which is also the covariance matrix because all $m$ Gaussian variables have unit variances). This null model assumes that, after each gene is transformed to a standard Gaussian random variable, the $n$ cells represent a random sample from an $m$-dimensional Gaussian distribution with zero means, unit variances, and a correlation matrix $\mathbf{R}$. Technically, to address the issue that the Gaussian copula is unidentifiable for discrete $F_j$'s, each $F_j$ is converted into a continuous CDF using a random interpolation procedure called the "distribution transform" (detailed below) [7]. Note that, for bulk microarray data, fitting the gene-wise Gaussian marginals and then estimating gene-gene correlation using a Gaussian copula is equivalent to directly modeling the joint distribution of all genes as a multivariate Gaussian distribution $N_m(\boldsymbol{\mu},\boldsymbol{\Sigma})$, where $\boldsymbol{\mu}=(\mu_1,\ldots,\mu_m)^\top$ and $\boldsymbol{\Sigma}=\text{diag}(\sigma_1,\ldots,\sigma_m)\,\mathbf{R}\,\text{diag}(\sigma_1,\ldots,\sigma_m)$.

In summary, the null models for scRNA-seq, scATAC-seq, bulk RNA-seq, and bulk microarray data are parameterized by $\{\mu_j,\sigma_j\}_{j=1}^m$ and $\mathbf{R}$, the null model for microbiome data is specified by $\{\alpha_j,\beta_j,\sigma_j\}_{j=1}^m$ and $\mathbf{R}$, and the SRT null model is specified by $\{\alpha_j,f_j^{GP},\sigma_j\}_{j=1}^m$ and $\mathbf{R}$.

2. **Fitting the null model to target data (parameter estimation)**
First, $F_j$ is fitted for each gene $j=1,\ldots,m$. For the null model of scRNA-seq, scATAC-seq, bulk microarray, bulk RNA-seq, or microbiome data, the parameters $\{\mu_j,\sigma_j\}_{j=1}^m$ are estimated using maximum likelihood estimation for the $m$ marginal distributions. Specifically, $Y_{1j},\ldots,Y_{nj}$ are used to estimate $\mu_j$ and $\sigma_j$ as $\widehat{\mu}_j$ and $\widehat{\sigma}_j$, respectively, for each $j=1,\ldots,m$. For the SRT null model, $Y_{1j},\ldots,Y_{nj}$ and spatial locations $X_1,\ldots,X_n$ are used to estimate $\mu_{ij}$ as $\widehat{\mu}_{ij}$ and $\sigma_j$ as $\widehat{\sigma}_j$ through a penalized Gaussian process. The resulting fitted CDFs are denoted as $\widehat{F}_1,\ldots,\widehat{F}_m$.
Second, to estimate the correlation matrix $\mathbf{R}$, each $Y_{ij}$ is transformed into $U_{ij}=V_{ij}\cdot\widehat{F}_j(Y_{ij})+(1-V_{ij})\cdot\widehat{F}_j(Y_{ij}+1)$, where $V_{ij}\overset{\text{i.i.d.}}{\sim}\text{Uniform}[0,1]$, so that $U_{ij}\sim\text{Uniform}[0,1]$. This procedure is referred to as the "distribution transform" to convert the discrete random variable $Y_{ij}$ into $U_{ij}$, a continuous Uniform$[0,1]$ random variable [7]. Then, $\mathbf{R}$ is estimated as the sample correlation matrix of the transformed data

$$\left(\Phi^{-1}(U_{11}),\cdots,\Phi^{-1}(U_{1m})\right)^\top, \\ \cdots, \\ \left(\Phi^{-1}(U_{n1}),\cdots,\Phi^{-1}(U_{nm})\right)^\top$$

and denoted as $\widehat{\mathbf{R}}$.
In summary, the fitted null model is specified by $\widehat{F}_1,\ldots,\widehat{F}_m$ and $\widehat{\mathbf{R}}$.

3. **Sampling from the fitted null model (synthetic null data generation)**
First, $n$ Gaussian vectors of $m$ dimensions are independently sampled from $N_m(\mathbf{0},\widehat{\mathbf{R}})$ as

$$(\widetilde{Z}_{11},\cdots,\widetilde{Z}_{1m})^\top,\ \cdots,\ (\widetilde{Z}_{n1},\cdots,\widetilde{Z}_{nm})^\top.$$

Second, the $n$ Gaussian vectors are converted to NB count vectors as

$$\widetilde{\mathbf{Y}}_1 := \left(\widehat{F}_1^{-1}(\Phi(\widetilde{Z}_{11})),\cdots,\widehat{F}_m^{-1}(\Phi(\widetilde{Z}_{1m}))\right)^\top, \\ \cdots,$$

$$\widetilde{\mathbf{Y}}_n := \left(\widehat{F}_1^{-1}(\Phi(\widetilde{Z}_{n1})), \cdots, \widehat{F}_m^{-1}(\Phi(\widetilde{Z}_{nm}))\right)^\mathsf{T},$$

which represent the $n$ synthetic null cells, each of which contains $m$ genes' synthetic null counts sampled from the null model.

In summary, the target data is an $n$-by-$m$ count matrix $\mathbf{Y}$, with the $n$ real cells $\mathbf{Y}_1, \ldots, \mathbf{Y}_n$ as the rows. Similarly, the synthetic null data is an $n$-by-$m$ count matrix $\widetilde{\mathbf{Y}}$, with the $n$ synthetic null cells $\widetilde{\mathbf{Y}}_1, \ldots, \widetilde{\mathbf{Y}}_n$ as the rows. Note that there is no one-to-one correspondence between the real cells and the synthetic null cells, as the synthetic null cells are independently sampled from the null model.

**S1.3.3 ClusterDE Step 2: cell clustering** While ClusterDE supports any clustering algorithm, we followed common practice by using the R package Seurat (version 4.2.0) for scRNA-seq data, Signac (version 1.13.0) for scATAC-seq data, and the R package BayesSpace (version 1.14.0) for SRT data. Specifically, we applied the same clustering algorithm to both the target data and the synthetic null data in parallel, resulting in two clusters for each dataset.

For scRNA-seq data, the default Seurat clustering procedure includes the following steps, applied separately to both the target data and the synthetic null data, each of which is stored as a Seurat object, denoted as `Seurat.obj`.

1. Normalize each cell to have a total count of 10,000; then perform log(normalized count + 1) transformation.
   `NormalizeData(Seurat.obj, normalization.method = "LogNormalize", scale.factor = 10000)`
2. Select 2,000 highly variable genes.
   `FindVariableFeatures(Seurat.obj, selection.method = "vst", nfeatures = 2000)`
3. Scale the data.
   `ScaleData(Seurat.obj)`
4. Run PCA on the data.
   `RunPCA(Seurat.obj, features = VariableFeatures())`
5. Compute cells' $k$-nearest neighbors.
   `FindNeighbors(Seurat.obj, dims = 1:30, nn.method = "rann", k.param = 20)`
6. Perform Louvain clustering on the cells.
   `FindClusters(Seurat.obj, resolution)`
   Since the Louvain clustering cannot pre-specify the cluster number, we tried resolutions starting from the default resolution of 0.5 and adjusted the resolution until two clusters were found.

For scATAC-seq data, the default Signac clustering procedure includes the following steps, applied separately to both the target data and the synthetic null data, each of which is also stored as a Signac object, denoted as `Signac.obj`.

1. Normalize each cell with term frequency-inverse document frequency (TF-IDF) normalization, which is a two-step normalization procedure that both normalizes across cells to correct for differences in cellular sequencing depth, and across peaks to give higher values to more rare peaks.
   `RunTFIDF(Signac.obj)`
2. Select the top highly variable genes.
   `FindTopFeatures(Signac.obj, min.cutoff = 'q0')`
3. Run singular value decomposition (SVD) on the TF-IDF matrix, using the peaks selected.
   `RunSVD(Signac.obj)`
4. Compute cells' $k$-nearest neighbors.
   `FindNeighbors(Signac.obj, dims = 1:30, nn.method = "rann", k.param = 20)`
5. Perform Louvain clustering on the cells.
   `FindClusters(Signac.obj, resolution)`
   Since the Louvain clustering cannot pre-specify the cluster number, we tried resolutions starting from the default resolution of 0.5 and adjusted the resolution until two clusters were found.

For SRT data, we employed the spatial clustering algorithm, BayesSpace, for spatial domain detection. Both the target and synthetic null datasets were stored as SingleCellExperiment objects, each denoted as `sce.obj`.

1. Perform log-normalization and then apply PCA.
```
spatialPreprocess(sce.obj, platform="ST", n.PCs=15, log.normalize=TRUE)
```
2. Cluster the spots and add the predicted domain labels to the `sce.obj`.
```
spatialCluster(sce.obj, q=2, platform="ST", d=7, init.method="mclust", model="t", gamma=2,
nrep=1000, burn.in=100, save.chain=TRUE)
```

For bulk microarray, bulk RNA-seq, and microbiome datasets, we applied either K-means or hierarchical clustering to identify subpopulations among samples, following the clustering strategies recommended by the original studies or dataset authors.

After applying the above clustering procedure, we obtained the cluster labels $\widehat{Z}_1, \ldots, \widehat{Z}_n$ from the target data $\mathbf{Y}$, and $\widetilde{Z}_1, \ldots, \widetilde{Z}_n$ from the synthetic null data $\widetilde{\mathbf{Y}}$, respectively, where $\widehat{Z}_i, \widetilde{Z}_i \in \{0, 1\}$, $i = 1, \ldots, n$. Again, there exists no one-to-one correspondence between $\widehat{Z}_1, \ldots, \widehat{Z}_n$ and $\widetilde{Z}_1, \ldots, \widetilde{Z}_n$.

**S1.3.4 ClusterDE Step 3: DE analysis** ClusterDE supports any DE test in principle. For scRNA-seq data, we employ five DE tests included in the Seurat `FindMarkers` function: the Wilcoxon rank-sum test (Wilcoxon, the default), t-test, negative binomial generalized linear model (NB-GLM), logistic regression model predicting cluster membership with a likelihood-ratio test (LR), and likelihood-ratio test for single-cell gene expression (bimod; see [8]). For scATAC-seq data, we use the Wilcoxon rank-sum test. For SRT data, we use the Wilcoxon rank-sum test and t-test, which are commonly applied to identify spatial-domain marker genes. For bulk microarray data, we use limma [9], Wilcoxon rank-sum test, and t-test. For bulk RNA-seq data, we only use DESeq2 [10] and edgeR [11] tests. For microbiome data, we use ANCOMBC2 [12], the Wilcoxon rank-sum test, and MaAsLin2 [13], which are standard methods for identifying differentially abundant genera.

For a given DE test applied to the target dataset, ClusterDE computes a $P$ value $P_j$ for each gene $j$ to test the double-dipping null hypothesis $H_{0j}^{\mathrm{DD}} : \mu_{0j}^{\mathrm{DD}} = \mu_{1j}^{\mathrm{DD}}$, where $\mu_{0j}^{\mathrm{DD}} = \mathbb{E}[Y_{ij} | \widehat{Z}_i = 0]$ and $\mu_{1j}^{\mathrm{DD}} = \mathbb{E}[Y_{ij} | \widehat{Z}_i = 1]$. The target DE score for gene $j$ is then defined as $S_j := -\log_{10} P_j$.

In parallel, on the synthetic null data, ClusterDE calculates a $P$ value $\widetilde{P}_j$ for each gene $j$ to test the null hypothesis $\widetilde{H}_{0j}^{\mathrm{DD}} : \widetilde{\mu}_{0j}^{\mathrm{DD}} = \widetilde{\mu}_{1j}^{\mathrm{DD}}$, where $\widetilde{\mu}_{0j}^{\mathrm{DD}} = \mathbb{E}[\widetilde{Y}_{ij} | \widetilde{Z}_i = 0]$ and $\widetilde{\mu}_{1j}^{\mathrm{DD}} = \mathbb{E}[\widetilde{Y}_{ij} | \widetilde{Z}_i = 1]$. The null DE score of gene $j$ is defined as $\widetilde{S}_j := -\log_{10} \widetilde{P}_j$.

In summary, ClusterDE generates target DE scores $S_1, \ldots, S_m$ and null DE scores $\widetilde{S}_1, \ldots, \widetilde{S}_m$ for $m$ genes.

**S1.3.5 ClusterDE Step 4: FDR control** Given the target DE scores $S_1, \ldots, S_m$ and the null DE scores $\widetilde{S}_1, \ldots, \widetilde{S}_m$, we use the FDR-control method Clipper [14] to identify DE genes at a target FDR threshold $q \in (0, 1)$. Given a set of identified DE genes (i.e., discoveries), the FDR is defined as

$$\mathrm{FDR} := \mathbb{E}\left[\frac{\#\text{ false discoveries}}{\#\text{ discoveries } \vee 1}\right],$$

where $a \vee b$ denotes the maximum of two numbers $a$ and $b$.

To ensure valid FDR control, Clipper requires each gene to have a contrast score such that the true non-DE genes (i.e., non-marker genes defined based on the ideal DE test) have contrast scores symmetric about zero. In ClusterDE, the contrast score for gene $j$ is defined as

$$C_j := S_j - \widetilde{S}_j .$$

Then ClusterDE uses Clipper to find a contrast score cutoff $T$ within $\mathcal{C}$ (the set of non-zero contrast score values) given the target FDR threshold $q$:

$$T := \min\left\{t \in \mathcal{C} : \frac{\mathrm{card}\left(\{j : C_j \le -t\}\right) + 1}{\mathrm{card}\left(\{j : C_j \ge t\}\right) \vee 1} \le q\right\}$$

and outputs $\{j \in \{1, \cdots, m\} : C_j \ge T\}$ as discoveries. Here, $\mathrm{card}(A)$ defines the cardinality (i.e., size) of a set $A$. This contrast score thresholding procedure for FDR control is from the knockoffs method [15].

Under the assumption that the majority of genes are true non-DE genes, the distribution of all genes' contrast scores is expected to have a mode at zero, thereby satisfying Clipper's symmetry requirement for contrast scores under the null hypothesis. Ideally, slightly less than 50% of all genes' contrast scores would be negative. However, in practice, this symmetry requirement may not hold in real datasets. For example:

1. The contrast score distribution might have a positive mode, resulting in too few negative contrast scores and inflated false discoveries.
2. Conversely, the distribution could have a negative mode, leading to too many negative contrast scores and reduced statistical power.

To address this, ClusterDE checks whether contrast scores are symmetric around zero using Yuen's trimmed-mean test (via the function `yuen.t.test()` from the R package PairedData (version 1.1.1)). This test examines the symmetry of the contrast score distribution after excluding the smallest 10% and largest 10% of the contrast scores.

If symmetry is rejected by Yuen's trimmed mean test, ClusterDE adjusts the contrast score distribution to better satisfy the symmetry requirement. In detail, ClusterDE employs "robust fitting of linear models" using the function `rlm()` from the R package MASS (version 7.3-60). A linear model is fitted between the target DE scores (response variable $y$) and the null DE scores (explanatory variable $x$), and the predicted values ($\hat{y}$) are used as the adjusted null DE scores. Then the adjusted contrast scores, defined as the differences between the target DE scores and the adjusted null DE scores, are expected to meet the symmetry requirement more closely.

To maintain conservatism in the adjustment process, ClusterDE applies a one-sided ("greater than") Yuen's trimmed mean test at a significance level of 0.001. Hence, adjustment is performed only when there are too few negative contrast scores, a scenario that would likely inflate false discoveries by Clipper.

#### S1.4 ClusterDE extensions

**S1.4.1 ClusterDE-Stable details** ClusterDE-Stable extends ClusterDE-Core by repeatedly generating synthetic null datasets and aggregating the results across these datasets, either combining a separation statistic into a $p$-value or combining gene-selection events into selection frequencies. In both cases, synthetic null datasets are generated following "ClusterDE Step 1: synthetic null data generation."

*Stable cluster-separation assessment.* ClusterDE-Stable can assess whether two groups of cells are sufficiently separated to be treated as distinct clusters. Following Grabski et al. [16], we quantify their separation using a normalized Ward linkage statistic.

Let $C_1$ and $C_2$ denote the two groups of cells, and let $\mathbf{x}_i$ denote the coordinates of cell $i$ in a low-dimensional representation, such as PC or UMAP space. For a set of cells $C$, define its within-cluster error sum of squares as

$$\mathrm{ESS}(C) = \sum_{i \in C} \left\| \mathbf{x}_i - \frac{1}{|C|} \sum_{k \in C} \mathbf{x}_k \right\|^2 .$$

The Ward linkage between $C_1$ and $C_2$ is

$$W(C_1, C_2) = \mathrm{ESS}(C_1 \cup C_2) - \{\mathrm{ESS}(C_1) + \mathrm{ESS}(C_2)\},$$

which measures the increase in within-cluster variation resulting from merging the two groups. We normalize this quantity by the total number of cells in the two groups:

$$T(C_1, C_2) = \frac{W(C_1, C_2)}{|C_1| + |C_2|}.$$

A larger value of $T(C_1, C_2)$ indicates stronger separation between the two groups. To assess whether the observed separation exceeds what would be expected under the synthetic null distribution, ClusterDE-Stable generates $B$ synthetic null datasets and computes the corresponding normalized Ward linkage statistics $T_1, \ldots, T_B$. Let $T_{\mathrm{obs}}$ denote the statistic computed from the observed data. The empirical $p$-value is

$$p = \frac{1}{B} \sum_{b=1}^{B} \mathbf{1}\{T_b \geq T_{\mathrm{obs}}\}.$$

If $p$ is below a prespecified significance level, such as 0.05, the two groups are considered to be significantly separated; otherwise, they are regarded as insufficiently separated and are candidates for merging.

*Stable DE gene identification.* ClusterDE-Stable adopts the stability selection framework of Meinshausen et al. [17]. It generates $B$ synthetic null datasets, runs ClusterDE-Core on each, and aggregates the results into gene-level selection frequencies.

Specifically, for each synthetic null dataset $b = 1, \ldots, B$, ClusterDE-Core outputs a set of reported DE genes, denoted by

$$\mathcal{G}_b \subseteq \{1, \ldots, m\},$$

at a target FDR level. For gene $j$, its selection frequency is defined as

$$s_j = \frac{1}{B} \sum_{b=1}^{B} \mathbf{1}\{j \in \mathcal{G}_b\},$$

where $s_j \in [0, 1]$ represents the fraction of runs in which gene $j$ is selected. Given a threshold $t_S \in [0, 1]$, typically set to 0.5, ClusterDE-Stable reports

$$\{j \in \{1, \ldots, m\} : s_j \geq t_S\}$$

as the set of stable DE genes.

**S1.4.2 ClusterDE-Tree details** ClusterDE-Tree is a hierarchical extension of ClusterDE-Stable for identifying reliable cell clusters and their stable markers from a potentially over-clustered hierarchy. Each leaf node corresponds to an initial cell cluster, and each internal node represents a binary split into two child subtrees. By default, the cell-cluster hierarchy is constructed using Seurat's `BuildClusterTree` function.

The ClusterDE-Tree workflow consists of two phases:

1. **Tree pruning.** Starting from the leaf nodes and proceeding upward toward the root, ClusterDE-Tree evaluates each internal split using the stable cluster-separation assessment described in ClusterDE-Stable (Section S1.4.1). For an internal node, its two child subtrees define two groups of cells. ClusterDE-Stable tests whether these groups exhibit statistically significant separation based on the normalized Ward linkage statistic calibrated using synthetic null datasets. If the separation is not significant, the two child subtrees are merged. If the separation is significant, the split is retained, and no further merging is performed above that split.
2. **Stable marker identification.** After tree pruning, each remaining leaf node represents a retained cell cluster. For each retained leaf, ClusterDE-Tree compares the cluster with its sister clade, defined as the other subtree descending from the same parent node. Stable DE markers are then identified using the stable DE gene identification procedure in ClusterDE-Stable (Section S1.4.1).

Thus, ClusterDE-Tree uses ClusterDE-Stable at both stages: first to determine which hierarchical splits represent statistically supported cluster distinctions, and then to identify stable DE markers for the retained clusters.

**S1.4.3 ClusterDE-Fast details** We implemented a fast approximation for ClusterDE, named ClusterDE-Fast, to reduce the computational cost of generating synthetic null data. Specifically, instead of jointly modeling all genes in the target data $\mathbf{Y}$ as in ClusterDE Step 1, ClusterDE-Fast models a low-dimensional representation of $\mathbf{Y}$. Because the number of latent dimensions is much smaller than the number of genes, this modeling step is substantially faster. The detailed procedure is as follows:

1. Normalize each cell to have a total count of 10,000; then apply log(normalized count + 1) transformation to obtain the normalized count matrix $\mathbf{E}$, where $E_{ij}$ is gene $j$'s normalized count in cell $i$.
2. Run PCA on $\mathbf{E}$ to obtain the cell-by-PC score matrix $\mathbf{H}$ and the PC-by-gene loading matrix $\mathbf{W}$. We retain the top 200 PCs by default in our implementation.

3. Compute the PCA reconstruction error matrix $\mathbf{R} = \mathbf{E} - \mathbf{HW}$, and then perform gene-wise bootstrap (independent bootstrap of each gene's values) on $\mathbf{R}$ to obtain $\mathbf{R}^{\text{boot}}$.
4. Independently model each PC's scores. Specifically, we assume that $H_{ij}$, the score of PC $j$ in cell $i$, independently follows a univariate normal distribution $N(\mu_j, \sigma_j^2)$ across cells. That is,

$$H_{1j}, \cdots, H_{nj} \overset{\text{i.i.d.}}{\sim} N(\mu_j, \sigma_j^2).$$

5. Obtain the estimated parameters $\{\hat{\mu}_j, \hat{\sigma}_j\}$ for each PC's marginal distribution using maximum likelihood estimation.
6. Independently sample from the fitted marginal distribution for each PC to obtain the synthetic score matrix $\widetilde{\mathbf{H}}$, project $\widetilde{\mathbf{H}}$ back to the gene space using the weight matrix $\mathbf{W}$, and construct the synthetic null data as $\widetilde{\mathbf{E}} = \widetilde{\mathbf{H}}\mathbf{W} + \mathbf{R}^{\text{boot}}$. Adding the bootstrapped reconstruction error helps preserve the noise structure of the target data in the synthetic null data.

Note that ClusterDE-Fast generates synthetic null data in the scale of normalized UMI counts rather than raw UMI counts.

To demonstrate the efficiency and effectiveness of ClusterDE-Fast, we analyzed the PBMC dataset focusing on CD4$^+$ and Cytotoxic T cell types. For ClusterDE-Stable, we generated $B$ synthetic null datasets following the procedure described in Step 1 of ClusterDE, in order to obtain gene-level selection frequencies denoted by $\mathbf{S}_{\text{Stable}}$. For ClusterDE-Fast, we similarly generated $B$ synthetic null datasets using the procedure detailed in this section to obtain $\mathbf{S}_{\text{Fast}}$. We considered values of $B$ from 1, 10, 20, . . ., up to 100. For each setting, we measured the total time required to generate synthetic null replicates, perform the DE analysis, and obtain the final DE genes. The runtime in seconds is summarized in Fig. S38, showing that ClusterDE-Fast substantially reduces computation time.

Then, we compared $\mathbf{S}_{\text{Stable}}$ and $\mathbf{S}_{\text{Fast}}$ ($B = 40$) using the Kuncheva index [18], which quantifies the overlap between two sets of selected genes while accounting for the overlap by chance. Genes were ranked by their selection frequencies in both $\mathbf{S}_{\text{Stable}}$ and $\mathbf{S}_{\text{Fast}}$, and the Kuncheva index was computed at varying values of $k$.

Formally, let $A_k$ and $B_k$ denote the sets of the top $k$ selected genes from two ranking lists, and let $m$ be the total number of candidate genes. The Kuncheva index is defined as:

$$\text{Kuncheva index}(A_k, B_k) = \frac{|A_k \cap B_k| - \frac{k^2}{m}}{k - \frac{k^2}{m}}, \quad k = 1, \ldots, m. \tag{S1}$$

The index ranges from $-1$ to 1, where 1 indicates perfect agreement, 0 corresponds to random overlap, and negative values suggest less agreement than expected by chance. In our setting, a high Kuncheva index across different values of $k$ indicates that ClusterDE-Fast yields gene selection results highly consistent with those of ClusterDE-Stable (Fig. S38).

**S1.4.4 Synthetic null data quality checks and simulator comparison** We compared synthetic null data generated by ClusterDE, ClusterDE-Fast, SPARSim, and ZINB-WaVE using eleven metrics in total. Using the A549 cell-line dataset, we evaluated each synthetic dataset using nine complementary diagnostics. Eight assessed agreement with the target data: per-gene means and variances, gene–gene rank correlations, normalized expression distributions (assessed via quantile–quantile plots), principal-component variance profiles, cell-embedding similarity (measured by the mean local inverse Simpson's index, mLISI, in the joint UMAP space), multivariate expression similarity (assessed by the Friedman–Rafsky test), and clustering structure (measured by the silhouette score after applying default Seurat clustering using the first 50 PCs and requesting two clusters). The ninth diagnostic assessed inferential performance based on the number of DE genes identified in the A549 cell line (where no DE genes are expected). Using the Rep1_10x(v3) monocyte dataset (CD14 versus CD16 monocytes), we further evaluated the inferential performance using two diagnostics: the numbers of known monocyte markers and housekeeping genes identified as DE.

**S1.4.5 ClusterDE trajectory-based DE** ClusterDE trajectory-based DE extends the synthetic null calibration principle from cluster-based DE to lineage-based DE along inferred trajectories. It combines

Slingshot-based lineage inference with ClusterDE's synthetic null generation and contrast-score FDR control to suppress spurious bifurcations and trajectory markers induced by over-clustering.

We assume a count matrix $\mathbf{Y} \in \mathbb{N}_{\geq 0}^{n \times m}$ and an initial clustering that defines one root cluster and two or more terminal clusters. The workflow proceeds in three main steps: lineage construction and DE testing on the target data, construction of a synthetic single-lineage null, and contrast-score-based FDR control.

1. **Lineage construction and DE testing on the target data.** Starting from $\mathbf{Y}$, we perform standard library-size normalization and obtain log-normalized expression values. PCA is applied to derive a low-dimensional embedding $\mathbf{X}$ for clustering and trajectory inference. Using the designated root and terminal clusters, we run Slingshot to construct a lineage structure. Slingshot assigns, for each cell $i$, pseudotime values and lineage weights $(w_{i\ell})_\ell$, where $w_{i\ell} \in [0,1]$ and $\sum_\ell w_{i\ell} = 1$. To obtain lineage-specific DE statistics for a bifurcation between two lineages, say lineage 1 and lineage 2, we restrict attention to non-root cells confidently assigned to exactly one lineage (e.g., $w_{i1} = 1$ or $w_{i2} = 1$ and $w_{i\ell} = 0$ for all other $\ell$). For each gene $j$, we apply the Wilcoxon rank-sum test to compare log-normalized expression between lineage-1 cells and lineage-2 cells, and denote the resulting target $P$-value by $P_j$. The target DE score is defined as
$$S_j := -\log_{10} P_j, \quad j = 1, \ldots, m.$$
2. **Synthetic single-lineage null construction.** To calibrate $S_j$ against a scenario with no true bifurcation, we construct a synthetic null dataset representing a single continuous lineage. We apply ClusterDE's null-generation procedure (ClusterDE-Core or ClusterDE-Fast) to the non-root cells, simulating synthetic counts under a unimodal multivariate distribution that corresponds to a homogeneous population. These synthetic counts are combined with the original root-cell counts to form a null dataset $\widetilde{\mathbf{Y}}$ with the same genes and cells as $\mathbf{Y}$. We normalize and log-transform $\widetilde{\mathbf{Y}}$, project the synthetic null cells into the PCA space learned from the target data, and apply clustering (e.g., K-means with the same number of clusters used for $\mathbf{Y}$) to obtain null cluster labels. Clusters from the null and target data are matched based on their centroids in PCA space (for example, using the Hungarian algorithm [19]), and the target root and terminal labels are transferred to the corresponding null clusters. Using these matched labels, we re-run Slingshot on $\widetilde{\mathbf{Y}}$, restrict DE analysis to non-root cells confidently assigned to exactly one lineage, and compute null Wilcoxon $P$-values $\widetilde{P}_j$ for the same lineage comparison. The null DE scores are then defined as
$$\widetilde{S}_j := -\log_{10} \widetilde{P}_j, \quad j = 1, \ldots, m.$$
3. **Contrast-score construction and FDR control.** ClusterDE trajectory-based DE uses the same contrast-score framework as ClusterDE-Core. For each gene $j$, we define the trajectory contrast score
$$C_j := S_j - \widetilde{S}_j, \quad j = 1, \ldots, m.$$
Genes with no genuine lineage-specific expression differences are expected to have contrast scores symmetric around zero, while true trajectory markers exhibit large positive $C_j$. Given the contrast scores $(C_j)_{j=1}^m$, we apply the Clipper procedure to identify DE genes at a target FDR threshold $q \in (0,1)$, using the same contrast-score thresholding scheme as in ClusterDE-Core. The resulting discoveries are interpreted as lineage-specific DE genes whose observed separation between lineages cannot be explained by clustering-induced bifurcations alone.

##### S1.5 scRNA-seq simulation studies and evaluations

**S1.5.1 scRNA-seq simulation designs** To benchmark post-clustering DE methods in terms of FDR and statistical power, we required known DE and non-DE genes. We used scDesign3 (version 0.99.0) [4] to generate realistic scRNA-seq data from parameters estimated from a real reference dataset.

Under each simulation setting, we simulated 200 replicates. Each replicate contained $n = 998$ cells and $m = 9239$ genes, matching the dimensions of the naïve cytotoxic T cells in the Zhengmix4eq dataset [20] after Seurat's default preprocessing step, which removed genes expressed at extremely low levels. In the following, we let $i$ and $j$ denote the indices of cells and genes, respectively, $i = 1, \ldots, n$; $j = 1, \ldots, m$. The simulation involved estimating the following **one-cell-type model parameters** from the naïve cytotoxic T cells using `scDesign3`. For details of the model formulation, refer to the section "ClusterDE Step 1: synthetic null data generation."

– Per-gene NB mean parameter $\mu_j \in \mathbb{R}^+$, $j = 1, \ldots, m$;
– Per-gene NB dispersion parameter $\sigma_j \in \mathbb{R}^+$, $j = 1, \ldots, m$;
– Gene-gene correlation matrix in the Gaussian copula $\mathbf{R} \in [-1, 1]^{m \times m}$.

Using the estimated model parameters $\{\widehat{\mu}_j, \widehat{\sigma}_j\}_{j=1}^m$ and $\widehat{\mathbf{R}}$, we designed two settings: (1) one cell type and (2) two cell types with 200 true DE genes.

**S1.5.2 Simulation setting with one cell type.** All $n = 998$ cells were simulated as belonging to one cell type, modeled by an MVNB distribution specified by the Gaussian copula with correlation matrix $\widehat{\mathbf{R}}$. Gene $j$'s counts followed an NB distribution with mean $\widehat{\mu}_j$ and dispersion $\widehat{\sigma}_j$, $j = 1, \ldots, m$. The R package `scDesign3` was used to simulate a cell-by-gene count matrix $\mathbf{Y} \in \mathbb{N}_{\geq 0}^{n \times m}$, representing the one-cell-type target data (Fig. S2) in simulation studies.

**S1.5.3 Simulation setting with two cell types and 200 true DE genes.** All $n = 998$ cells were assigned to two cell types, each modeled by its own MVNB distribution specified by the Gaussian copula. To simulate each replicate, we randomly selected 200 true DE genes with distinct mean parameters $\mu_j^0$ and $\mu_j^1$ for the two cell types, based on $\widehat{\mu}_j$.

For the two cell types, we simulated three cell-type size ratios, $r \in \{1, 4, 9\}$, such that cells $i = 1, \ldots, \lfloor n/(r+1) \rfloor$ belonged to cell type 0, and cells $i = \lfloor n/(r+1) \rfloor + 1, \ldots, n$ belonged to cell type 1. For each replicate, we specified 200 true DE genes with the index set $J_{\mathrm{DE}} \subset \{1, \cdots, m\}$. For each true DE gene $j \in J_{\mathrm{DE}}$, we set its mean parameter in cell type 0 as the estimate, i.e., $\mu_j^0 = \widehat{\mu}_j$. Then we modified its mean parameter in cell type 1, $\mu_j^1$, using a pre-specified log fold change, logFC, with a 50% probability of up-regulation and a 50% probability of down-regulation:

$$\mu_j^1 = \begin{cases} \widehat{\mu}_j \times 2^{\mathrm{logFC}}, & \text{if } Z_j = 1 \\ \widehat{\mu}_j \times 2^{-\mathrm{logFC}}, & \text{if } Z_j = 0 \end{cases}, \quad \text{with } Z_j \sim \mathrm{Ber}(0.5), \quad j \in J_{\mathrm{DE}}\,.$$

For each true non-DE gene $j$, we set

$$\mu_j^0 = \mu_j^1 = \widehat{\mu}_j, \quad j \in \{1, \cdots, m\} \backslash J_{\mathrm{DE}}\,.$$

The pre-specified logFC determines the differences between the two cell types and is expected to have an inverse relationship with the severity of double dipping (i.e., greater differences between the two cell types lead to better clustering, which in turn reduces the severity of double dipping). Accordingly, we simulated two cell types using a sequence of logFC values

$$\mathrm{logFC} = 1.05, 1.1, \ldots, 1.95, 2, 2.1, \ldots, 2.9, 3.$$

For each logFC value, we simulated cells from cell types 0 and 1, each modeled by an MVNB distribution specified by the Gaussian copula with correlation matrix $\widehat{\mathbf{R}}$. Specifically, gene $j$'s counts in cell types 0 and 1 followed NB distributions with distinct mean parameters $\mu_j^0$ and $\mu_j^1$, respectively, and a shared dispersion parameter $\widehat{\sigma}_j$, $j = 1, \ldots, m$. For detailed simulation procedures, refer to the section "ClusterDE Step 1: synthetic null generation." The R package scDesign3 was used to simulate a cell-by-gene count matrix $\mathbf{Y} \in \mathbb{N}_{\geq 0}^{n \times m}$, which served as the two-cell-type target data (Fig. S2) in simulation studies.

**S1.5.4 Implementation of the TN test and Countsplit** We compared ClusterDE with two existing methods—the TN test [21] and Countsplit [22]—both designed to address the double-dipping issue in post-clustering DE analysis.

For the TN test, we used the Python module truncated-normal (version 0.4). We followed the GitHub tutorial for the implementation (`https://github.com/jessemzhang/tn_test/blob/master/experiments/experiments_pbmc3k.ipynb`). In the clustering step, we used the same procedure as in ClusterDE Step 2. In the DE analysis step, unlike ClusterDE and Countsplit, the TN test has its own DE test, so we did not use any DE tests from the R package Seurat (version 4.2.0).

For Countsplit, we used the R package countsplit (version 1.0) to split the original count matrix (i.e., the target data used by ClusterDE) into a training matrix (for clustering) and a test matrix (for DE analysis). In the clustering step, we used the same procedure as in ClusterDE Step 2. In the DE analysis step, we used the five DE tests provided in the R package Seurat (version 4.2.0).

**S1.5.5 Alternative strategies for synthetic null data generation** Although the model-X knockoffs method was originally developed for feature selection in multivariate predictive models (e.g., the Lasso) [23], rather than for marginal DE tests (where each feature is analyzed independently), we compared it to ClusterDE because both methods share the concept of generating synthetic null data (or negative-control data) based on real data.

To implement the model-X knockoffs method for post-clustering DE analysis, we used the R package knockoff (version 0.3.6) to construct knockoff data (i.e., the negative control). For feature selection, we applied the default `glmnet` method for binary logistic regression, treating the cluster labels as the response variable ($y$) and the genes as features. Testing this approach on 50 simulated datasets with logFC = 2.6 (see "Simulation setting with two cell types and 200 true DE genes"), we observed that it consistently selected 0 DE genes, resulting in a power of 0.

Moreover, we tested three alternative strategies for synthetic null data generation in ClusterDE Step 1: the knockoff data constructed above, permuted data (where each gene was independently permuted across all cells), and scDesign3 without copula modeling (assuming gene independence) (Fig. S11). Our results on simulated datasets showed that scDesign3 provided more robust FDR control and better statistical power compared to these three alternative strategies for synthetic null data generation (Fig. S12).

**S1.5.6 Validity checks of the contrast scores of ClusterDE and the *P* values of Seurat, Countsplit, and the TN test** For ClusterDE, the primary assumption is that the contrast scores of true non-DE genes are symmetric around zero. In Fig. S5 (left), we evaluated the symmetry of the contrast scores of ClusterDE using five DE tests (Wilcoxon, t-test, NB-GLM, LR, and bimod; corresponding to Fig. S5a–e, left) on a simulated one-cell-type dataset where all genes are true non-DE genes (see "Simulation setting with one cell type" in the Supplementary Methods). The dataset used is one of the 200 simulated replicates.

For Seurat, Countsplit, and the TN test, the validity of their FDR control relies on the assumption that the $P$ values of true non-DE genes follow the Uniform$[0, 1]$ distribution. To examine this, we divided the genes in the same simulated dataset into two groups using hierarchical clustering (via the default R function `hclust()`) applied to the estimated correlation matrix $\widehat{\mathbf{R}}$ from the Gaussian copula. Due to the block pattern in $\widehat{\mathbf{R}}$ (Fig. 2a, right), the two groups consisted of genes that are highly correlated and those that are less correlated, respectively. We analyzed the $P$ values of the genes in these two groups separately, noting that all genes in this simulated dataset are true non-DE genes. In Fig. S5 (middle and right), we plotted histograms of the $P$ values and the quantile-quantile plots (Q-Q plots) of the negative log-transformed $P$ values for Seurat and Countsplit (both using the five DE tests; corresponding to Fig. S5a–e) and for the TN test (using its own test; the same panels repeated five times in Fig. S5a–e). To quantify the deviation of the $P$ value distribution from the theoretical Uniform$[0, 1]$ distribution, we used the R function `KL.empirical` (from the R package entropy, version 1.3.1)) to calculate the empirical Kullback–Leibler divergence (KL divergence). A larger KL divergence indicates a more severe violation of the $P$ value uniformity assumption. The results demonstrated that Countsplit and the TN test produced nearly uniform $P$ values for the uncorrelated gene group. However, their $P$ values deviated substantially from the uniform distribution for the correlated gene group.

**S1.5.7 Random-seed stability and robustness to null-parameter perturbation** To assess sensitivity to random synthetic null generation, we generated 50 synthetic null datasets with distinct random seeds for a target dataset simulated with two cell types and 200 true DE genes. ClusterDE-Core was applied at a range of target FDR levels, and stability was summarized by the numbers of selected genes and pairwise overlaps across random seeds.

To assess robustness to null-model estimation error, we perturbed gene means, gene dispersions, and gene–gene correlations separately and jointly, across varying proportions of genes and two perturbation magnitudes. For each perturbed null, ClusterDE-Stable selection frequencies were compared with those obtained from the unperturbed null using the correlation between gene-level selection-frequency vectors.

#### S1.6 scRNA-seq real-data analyses

**S1.6.1 Pure cell-line datasets** We analyzed five scRNA-seq cell-line datasets. A549, H2228, and HCC827 were obtained from Tian et al. [24] and downloaded from https://github.com/LuyiTian/sc_mixology/tree/master/data. HEK293T and JURKAT were obtained from Zheng et al. [25] and downloaded from https://cf.10xgenomics.com/samples/cell-exp/1.1.0/293t/293t_filtered_gene_bc_matrices.tar.gz and https://cf.10xgenomics.com/samples/cell-exp/1.1.0/jurkat/jurkat_filtered_gene_bc_matrices.tar.gz, respectively. Genes expressed in fewer than 20% of cells were removed from A549, H2228, and HCC827, and genes expressed in fewer than 10% of cells were removed from HEK293T and JURKAT.

**S1.6.2 $CD14^+/CD16^+$ monocyte datasets** We analyzed eight PBMC datasets from Ding et al. [26], downloaded from https://github.com/satijalab/seurat-data. The datasets were generated from the same biological sample using two technical replicates and four protocols: 10X Genomics v2, 10X Genomics v3, Drop-seq, and inDrop. For each dataset, we retained cells annotated as $CD14^+$ or $CD16^+$ monocytes and removed genes expressed in fewer than 10% of the retained cells. During default Seurat clustering, cells with fewer than three expressed genes and genes expressed in fewer than 200 cells were filtered automatically.

**S1.6.3 Dimensionality reduction and visualization** To visualize the high-dimensional single-cell data, we first applied the PF-logPF transformation to the cell-by-gene count matrix, as described in [27]. We then calculated the top 50 principal components (PCs) of the transformed matrix using the R package irlba (version 2.3.5.1). These PCs were subsequently used as input to the R package umap (version 0.2.10.0) to project the cells from the 50-dimensional PC space to a 2-dimensional UMAP space. For the *Drosophila* dataset, the original authors used 120 PCs as input for t-SNE.

When comparing the target data with the synthetic null data, we calculated the PCs and UMAPs jointly by concatenating the two datasets. This ensured that the target cells and synthetic null cells were projected into the same 2-dimensional UMAP space.

All plots were generated using the R package ggplot2 (version 3.4.2).

For the UMAP visualizations in Fig. 3f, we truncated each gene's normalized expression levels to the 99th percentile to better illustrate gene expression patterns.

**S1.6.4 Gene set enrichment analysis for monocyte markers and housekeeping genes** We performed gene set enrichment analysis (GSEA) using the R package clusterProfiler (version 4.4.4). The test method used was `fgsea`, with the number of permutations set to 100,000. The gene set "$CD14^+/CD16^+$ Monocyte Markers" was obtained from the original study [26] and downloaded from https://bitbucket.org/jerry00/scumi-dev/raw/61f7f001d20b2fc8fa7c2f4f4147bff1b0d620d8/R/marker_gene/human_pbmc_marker.rda. The gene set "Housekeeping Genes" (`HSIAO_HOUSEKEEPING_GENES`) was sourced from the Molecular Signature Database (MSigDB) and originated from the study [28].

**S1.6.5 Adult *Drosophila* neuronal visual atlas** We used the processed adult *Drosophila* neuronal atlas from Ozel et al. [29], including the original cell annotations and clustering (GSE142789; https://www.ncbi.nlm.nih.gov/geo/query/acc.cgi?acc=GSE142789). In the original preprocessing, cells were required to contain at least 1,000 UMIs and less than 10% mitochondrial UMIs, and genes expressed in fewer than three cells were removed. Louvain clustering used resolution 10 and 120 PCs. Over-clustering was assessed with Seurat v2 `AssessNodes`, which fits a random-forest classifier for each candidate split and reports the out-of-bag error; retained splits were required to have an error below 5% and clear between-cluster expression differences.

*Gating plots.* We used gating plots to compare two clusters at a time. DE genes between the two clusters were identified using the Wilcoxon rank-sum test. DE genes were included as marker genes only if their adjusted $P$ values were less than 0.05 and their absolute log-fold changes between the two clusters exceeded 0.5. Identified markers were categorized into positive and negative markers for each cluster based on the log-fold change. To calculate the marker rate, we determined the proportion of counts attributed to each set of marker genes out of the total gene count for each cell.

*Heatmaps.* We generated gene expression heatmaps using scaled data, with genes ordered by hierarchical clustering. The heatmaps were created using the `heatmap.2` function from the R package gplots (version 3.1.3).

*Machine-learning evaluation.* For each machine learning model, the *Drosophila* visual atlas scRNA-seq data were subset to include only cell types Tm1 and Tm2, along with one of the three specified gene sets: (1) all 105 DE genes identified by ClusterDE, (2) the top 105 genes ranked by increasing $P$ value from the Wilcoxon rank-sum test in Seurat, and (3) a random selection of 105 genes. The data were split into an 80-20 train-test split, and this process was repeated 100 times. A new model was trained for each split using the genes from the respective set. Random forest models were trained using the `randomForest` function from the R package randomForest (version 4.7.1.1), and SVM models were trained using the `svm` function from the R package e1071 (version 1.7.13).

**S1.6.6 Retracted human fetal cerebellum TCP dataset** We reanalyzed the human fetal cerebellum single-cell RNA-seq dataset from Luo et al. [30], focusing on the putative "transitional cerebellar progenitor" (TCP) population and related progenitor states. Following Smith et al. [31], we used the authors' processed data and cell-type annotations, with neural stem cells (NSCs), TCPs, unipolar brush cell progenitors (UBC_P), granule cell progenitors (GCPs), and Purkinje cells as originally defined in the study. For our ClusterDE analysis, we restricted attention to cells annotated as TCP and UBC_P, treating these two annotations as the candidate cell types to be compared.

We applied the same preprocessing and dimensionality reduction steps as in the original study (including normalization, feature selection, and principal component analysis), and embedded the cells in the authors' low-dimensional space for clustering and visualization. ClusterDE was run on the TCP and UBC_P subset using the scRNA-seq null model described in "ClusterDE Step 1: synthetic null data generation" (Section S1.3.2). Synthetic null data were generated to preserve per-gene means, variances, and gene–gene rank correlations while representing a single homogeneous progenitor population, and we verified that synthetic and target cells were well mixed in the joint embedding (Fig. 4h). We then applied the clustering pipeline as in Section S1.3.3 to the synthetic null data and the same DE pipeline to both the target and synthetic null data, followed by contrast-score construction as in Section S1.3.5. For the DE pipeline, we applied Seurat's standard post-clustering DE workflow directly to cells of the two selected annotations (TCP and UBC_P), using four DE tests (Wilcoxon, t-test, LR, and bimod) with Benjamini–Hochberg correction at FDR 0.05. We summarized the total number of DE genes reported by ClusterDE and standard Seurat and examined their expression patterns (Fig. 4i–j).

We also performed Gene Ontology enrichment analysis on the Seurat Wilcoxon DE gene set, highlighting terms related to renal, kidney, cartilage development, gliogenesis, and glial cell differentiation (Fig. 4k), which are inconsistent with the expected neuronal fate of these cerebellar progenitors.

Gene Ontology enrichment. We performed Gene Ontology (GO) enrichment analysis using the R package clusterProfiler (version 4.4.4). The function `enrichGO` was applied to identify over-represented Biological Process (BP) categories among the selected genes, using `org.Hs.eg.db` as the annotation database. The enrichment test was based on a hypergeometric framework with default multiple-testing correction, and results were visualized using the `dotplot` function in clusterProfiler.

**S1.6.7 COVID-19 PBMC dataset** We analyzed the peripheral blood mononuclear cell (PBMC) scRNA-seq dataset from Wilk et al. [32], which profiled peripheral immune cells from hospitalized patients with COVID-19, including patients with acute respiratory distress syndrome (ARDS). In the original analysis, genes detected in at least 10 cells were retained, and the data were normalized using `SCTransform`, with mitochondrial, ribosomal, and rRNA percentages, total UMI counts, and the number of detected genes regressed out. PCA, UMAP embedding, nearest-neighbor graph construction, and clustering were performed in Seurat using the first 50 principal components, followed by removal of two low-quality clusters and re-clustering. These procedures yielded the original cell-type annotations, including IgM plasmablasts (IgM PBs) and "developing neutrophils." We used the original authors' processed data and cell-type annotations, available from the COVID-19 Cell Atlas at https://www.covid19cellatlas.org/#wilk20.

For the analysis reported in the main Results, we subsetted cells annotated as IgM PBs or "developing neutrophils" and applied the same ClusterDE workflow as in Section S1.6.6. Genes were ranked according to the ClusterDE and standard Wilcoxon results, respectively, and representative genes identified by ClusterDE or uniquely by the standard Wilcoxon analysis were visualized to compare their expression patterns between the two annotated populations (Fig. S24c–d).

**S1.6.8 PBMC CD4$^+$ and cytotoxic T-cell subsets** We analyzed a PBMC scRNA-seq dataset derived from the Zhengmix4eq benchmark panel [20], focusing on CD4$^+$ T cells and cytotoxic T cells as defined by the original annotations. Cells were restricted to these two subsets, and genes were preprocessed using the same filtering thresholds as in the Zhengmix4eq study (removal of genes expressed at extremely low levels and standard quality-control filters), consistent with the preprocessing used in our scRNA-seq simulation designs (Section S1.5).

For this subset, we applied ClusterDE-Stable (Section S1.4.1) using ClusterDE-Core to generate multiple synthetic null datasets, and we applied ClusterDE-Fast (Section S1.4.3) to approximate the synthetic null distribution in a low-dimensional PCA space. We compared gene-level selection frequencies between ClusterDE-Stable and ClusterDE-Fast across varying numbers of synthetic null replicates $B$, summarized runtimes (null generation plus DE analysis) for each method, and quantified agreement in ranked marker lists using the Kuncheva index. The results, reported in Fig. S38, show that ClusterDE-Fast substantially reduces computation time while maintaining high agreement with ClusterDE-Stable in selected markers for CD4$^+$ and cytotoxic T cells.

**S1.6.9 H2228 cell line and human glutamatergic neurogenesis datasets** We used two datasets known not to contain true bifurcating trajectories as negative controls for trajectory-based DE analysis: a homogeneous lung adenocarcinoma cell line, H2228 [24], which lacks any developmental trajectory, and a human embryonic glutamatergic neurogenesis dataset [33], in which cells progress along a single continuous lineage from radial glia to neuroblasts.

For each dataset, we applied a standard clustering→trajectory→DE workflow using Slingshot followed by Wilcoxon rank-sum testing between inferred terminal clusters, and compared it with ClusterDE trajectory-based DE (Section S1.4.5; see also "Practical guidelines for applying ClusterDE to trajectory-based analyses," Section S1.2.4). The standard pipeline identified 5,560 DE genes (77% of all genes) in H2228 and 1,216 DE genes in the neurogenesis dataset, whereas ClusterDE identified zero DE genes in both datasets (Fig. S25b,e), correctly indicating that the inferred bifurcations in both datasets were artifacts of over-clustering rather than true lineage structure.

**S1.6.10 Mouse bone marrow dataset** We evaluated ClusterDE's sensitivity to true lineage structure using a well-characterized mouse bone marrow scRNA-seq dataset [34], in which common myeloid progenitors (CMPs) bifurcate into granulocyte–monocyte progenitors (GMPs) and erythroid precursors.

Following the original study's clustering, we designated a CMP cluster as the root and identified terminal clusters corresponding to the GMP and erythroid branches. ClusterDE generated synthetic null data representing a hypothetical single-lineage scenario (Fig. S26a, middle and bottom) to assess whether the observed lineage differences were statistically supported, following the trajectory-based DE procedure in Section S1.4.5. Compared with the standard clustering→trajectory→DE pipeline, ClusterDE more accurately identified genes with clear lineage-specific expression patterns: the top DE genes from ClusterDE showed clear separation between the two lineages (Fig. S26b), whereas genes uniquely reported by the standard pipeline exhibited ambiguous differences and are likely false positives (Fig. S26c). Known lineage markers—including *Klf1*, *Elane*, *Gata1*, *Fcgr3*, *Cebpa*, and *Cd34*—received consistently higher rankings under ClusterDE (Fig. S26d), and heatmaps along pseudotime further showed that genes identified by ClusterDE display clear lineage-specific expression, whereas those unique to the standard pipeline do not (Fig. S26e).

**S1.6.11 Allen Human MTG atlas analysis** We analyzed the Allen Human Middle Temporal Gyrus (MTG) snRNA-seq taxonomy, in which transcriptomic "supertypes" define a hierarchy of neuronal and non-neuronal cell types across the cortical depth [35]. The MTG dataset was downloaded from `https://brain-map.org/consortia/sea-ad/human-mtg-10x-sea-ad`, which collected 166,868 nuclei from 5 healthy donors and was annotated with 127 clusters (transcriptomic supertypes), as well as the scVI embeddings of all nuclei across individuals (batch-corrected).

To refine this atlas-scale taxonomy, we applied ClusterDE-Tree (Section S1.4.2) to the MTG "supertypes." We first constructed a binary cluster tree by computing pairwise cosine similarities of cluster centroids based on batch-corrected scVI embeddings across all individuals, and then ran ClusterDE-Tree in its standard two-phase mode: recursive split testing followed by marker selection. For each internal node, ClusterDE-Tree computed a Ward-linkage split statistic and calibrated it against multiple synthetic null datasets generated with ClusterDE-Fast (Section S1.4.3). The $P$-value was determined by taking the mean of frequencies that for each individual the linkage statistic of null replicates exceeds the linkage statistic of observed data (so each individual received one frequency, and the frequencies were averaged across individuals). Then splits were pruned if $P$-values were greater than 0.05, and the corresponding clusters were merged. This procedure merged the original 127 MTG clusters into 76 refined clusters, reducing over-clustering while preserving transcriptomically distinct cell types, including a notable merge of ten layer 2/3 intratelencephalic (L2/3 IT) clusters into two refined clusters (L2/3 IT_C7 and L2/3 IT_C8) that exhibited clear between-cluster marker contrasts (Fig. 5c–e).

For each retained leaf in the pruned MTG tree, we then used ClusterDE-Stable (Section S1.4.1) to identify stable marker genes by comparing the leaf to its sister clade within each individual at a target FDR, so that every gene received a selection frequency within that individual. We then averaged the selection frequency across individuals and reported genes whose averaged selection frequency exceeded 0.5. To assess whether the ClusterDE-Tree–based refinement corresponded to biologically meaningful spatial organization, we mapped both the original and refined MTG clusters to an independent Allen Human MTG MERFISH dataset [36]. Spatial spots annotated by the two refined L2/3 IT clusters formed two well-segregated L2/3 IT regions whose spatial boundaries agreed with the expression patterns of ClusterDE markers, whereas spatial spots annotated by the original ten L2/3 IT clusters showed substantial overlap and poor spatial separability (Fig. 5f–g). These results indicate that ClusterDE-Tree can refine an atlas-scale MTG taxonomy by merging spurious over-resolved clusters while retaining clusters that are both transcriptomically and spatially distinct.

#### S1.7 Spatially resolved transcriptomics (SRT) data analyses

**S1.7.1 SRT simulation designs** We assume that the expression of non-domain marker genes changes gradually across spatial locations, while the expression of domain marker genes shows significant discontinuities (sudden changes) at domain boundaries. In our modeling of SRT data, gene $j$'s counts are assumed to follow a negative binomial (NB) distribution with spot-specific mean $\mu_{ij}$ and a common dispersion parameter $\sigma_j$ $(i = 1, \ldots, n; j = 1, \ldots, m)$. Additionally, all genes are modeled to follow an MVNB distribution specified by the Gaussian copula with correlation matrix $\mathbf{R}$. We simulated three settings: (1) one spatial domain; (2) two domains with 200 true domain marker genes; (3) three domains with 200 and 168 true domain marker genes between adjacent domains.

**S1.7.2 Simulation setting with one spatial domain** A single domain ($n = 989$ spots) was simulated using Layer 3 of slice 151673 from the LIBD human dorsolateral prefrontal cortex (DLPFC) dataset as the reference. We modeled gene $j$'s mean expression as $\log(\mu_{ij}) = \alpha_j + f_j^{GP}(x_{i1}, x_{i2}, K)$, where $\alpha_j$ denotes gene $j$'s specific intercept, and $f_j^{GP}(\cdot)$ represents a smooth function modeled by a penalized Gaussian Process regressor with $K$ bases. We set $K = 10$ to constrain the mean expression surface to be smooth. The R package scDesign3 was used to simulate a spot-by-gene count matrix $\mathbf{Y} \in \mathbb{N}_{\geq 0}^{n\times m}$, based on $\log(\widehat{\mu}_{ij}) = \widehat{\alpha}_j + \widehat{f}_j^{GP}(x_{i1}, x_{i2}, K)$, $\widehat{\sigma}_j$ $(j = 1, \ldots, m)$, and $\widehat{\mathbf{R}}$ estimated from the reference data. The simulated matrix, along with the spots' 2D locations (identical to those in the reference data), served as the one-domain target data for the simulation studies.

**S1.7.3 Simulation setting with two spatial domains and 200 true spatial-domain marker genes** All $n = 1207$ spots from Layers 3 and 4 of slice 151673 in the DLPFC dataset were used as reference data to simulate SRT data with two domains. We selected 200 genes as the true domain marker genes for Layer 4, meaning their expression values exhibited abrupt changes between the two domains.

To select these 200 genes, for each gene $j$, we modeled the mean parameter $\mu_{ij}$ as:

$$\log(\mu_{ij}) = \alpha_{0j} + \alpha_{1j}\mathbb{I}(Z_i = 1) + f_j^{GP}(x_{i1}, x_{i2}, K),$$

where $\alpha_{0j}$ is the intercept, $\alpha_{1j}$ represents the domain effect, $Z_i$ is the domain label for spot $i$ ($Z_i = 0$ corresponds to Layer 3, and $Z_i = 1$ corresponds to Layer 4), and $K = 10$. The model was fitted to the reference data to obtain parameter estimates $\widehat{\alpha}_{0j}$, $\widehat{\alpha}_{1j}$, and $\widehat{f}_j^{GP}$, as well as the dispersion estimate $\widehat{\sigma}_j$. The 200 true domain marker genes were then selected as those with the largest differences $\widehat{\alpha}_{1j}/\widehat{\alpha}_{0j} - 1$.

For the remaining genes, considered true non-marker genes, we modeled the mean parameter $\mu_{ij}$ as:

$$\log(\mu_{ij}) = \alpha_j + f_j^{GP}(x_{i1}, x_{i2}, K).$$

This model was fitted to data from both domains of the reference set to obtain estimates $\widehat{\alpha}_j$, $\widehat{f}_j^{GP}$, and the dispersion $\widehat{\sigma}_j$.

To amplify the differences between the two domains, a constant value of 2 was manually added to enhance the domain effect. Consequently, the mean parameter used to simulate a true domain marker gene $j$ for Layer 4 was defined as:

$$\log(\widehat{\mu}_{ij}) = \widehat{\alpha}_{0j} + (\widehat{\alpha}_{1j} + 2)\mathbb{I}(Z_i = 1) + \widehat{f}_j^{\,GP}(x_{i1}, x_{i2}, K).$$

The R package scDesign3 was used to simulate a spot-by-gene count matrix $\mathbf{Y} \in \mathbb{N}_{\geq 0}^{n \times m}$, based on $\log(\widehat{\mu}_{ij})$, $\widehat{\sigma}_j$ $(j = 1, \ldots, m)$, and $\widehat{\mathbf{R}}$ estimated from the reference data. The simulated matrix, along with the spots' 2D locations (identical to those in the reference data), served as the two-domain target dataset for the simulation studies.

**S1.7.4 Simulation setting with three spatial domains and 200 and 168 true spatial-domain marker genes** All $n = 1878$ spots from Layers 5, 6, and WM of slice 151673 in the DLPFC dataset were used as reference data to simulate SRT data with three domains. We selected 200 and 168 genes as true domain marker genes for Layers 5 and 6, respectively, indicating that their expression values exhibited abrupt changes between the adjacent domains.

To select these true domain marker genes, for each gene $j$, we modeled the mean parameter $\mu_{ij}$ as:

$$\log(\mu_{ij}) = \alpha_{0j} + \alpha_{1j}\mathbb{I}(Z_i = 1) + \alpha_{2j}\mathbb{I}(Z_i = 2) + f_j^{GP}(x_{i1}, x_{i2}, K),$$

where $\alpha_{0j}$ is the intercept, $\alpha_{1j}$ and $\alpha_{2j}$ represent domain effects, $Z_i$ is the domain label for spot $i$ ($Z_i = 0$ corresponds to Layer 5, $Z_i = 1$ to Layer 6, and $Z_i = 2$ to Layer WM), and $K = 10$. The model was fitted to the reference data to obtain parameter estimates $\widehat{\alpha}_{0j}$, $\widehat{\alpha}_{1j}$, $\widehat{\alpha}_{2j}$, and $\widehat{f}_j^{GP}$, as well as the dispersion estimate $\widehat{\sigma}_j$. The 200 true domain marker genes for Layer 5 were first selected as those with the largest $|\widehat{\alpha}_{0j}/\widehat{\alpha}_{1j} - 1|$. Then, an additional set of 200 genes for Layer 6 were selected based on the largest $|\widehat{\alpha}_{1j}/\widehat{\alpha}_{2j} - 1|$, with overlap from the first set removed, resulting in 168 unique genes.

For the remaining genes, considered true non-marker genes, we modeled the mean parameter $\mu_{ij}$ as:

$$\log(\mu_{ij}) = \alpha_j + f_j^{GP}(x_{i1}, x_{i2}, K).$$

This model was fitted to data from all three domains of the reference set to obtain estimates $\widehat{\alpha}_j$, $\widehat{f}_j^{GP}$, and the dispersion $\widehat{\sigma}_j$.

To amplify the differences between the three domains, constant values of 1 and 2.5 were manually added to the domain-specific intercepts for Layers 5 and 6, respectively. Consequently, the mean parameter used to simulate a true domain marker gene $j$ for Layer 5 was defined as:

$$\log(\widehat{\mu}_{ij}) = (\widetilde{\alpha}_j + 1) + \widetilde{\alpha}_j\mathbb{I}(Z_i = 1) + \widetilde{\alpha}_j\mathbb{I}(Z_i = 2) + \widehat{f}_j^{GP}(x_{i1}, x_{i2}, K),$$

and the mean parameter used to simulate a true domain marker gene $j$ for Layer 6 was defined as:

$$\log(\widehat{\mu}_{ij}) = \widetilde{\alpha}_j + (\widetilde{\alpha}_j + 2.5)\mathbb{I}(Z_i = 1) + \widetilde{\alpha}_j\mathbb{I}(Z_i = 2) + \widehat{f}_j^{GP}(x_{i1}, x_{i2}, K),$$

where $\widetilde{\alpha}_j$ was set to the minimum of $\widehat{\alpha}_{0j}$, $\widehat{\alpha}_{1j}$, and $\widehat{\alpha}_{2j}$.

The R package scDesign3 was used to simulate a spot-by-gene count matrix $\mathbf{Y} \in \mathbb{N}_{\geq 0}^{n \times m}$, based on $\log(\widehat{\mu}_{ij})$, $\widehat{\sigma}_j$ $(j = 1, \ldots, m)$, and $\widehat{\mathbf{R}}$ estimated from the reference data. The simulated matrix, along with the spots' 2D locations (identical to those in the reference data), served as the three-domain target dataset for the simulation studies.

**S1.7.5 DLPFC and PDAC SRT datasets** We analyzed slice 151673 from the LIBD human dorsolateral prefrontal cortex dataset [37], available from http://research.libd.org/spatialLIBD/, and the PDAC-B section from the human pancreatic ductal adenocarcinoma dataset [38], available under GSE111672 at https://www.ncbi.nlm.nih.gov/geo/query/acc.cgi?acc=GSE111672. Genes expressed in fewer than 20% of spots were removed from the DLPFC slice, and genes expressed in fewer than 5% of spots were removed from PDAC-B. Spatial clustering and post-clustering DE testing followed the BayesSpace and Seurat procedures described in the core ClusterDE workflow above.

#### S1.8 Analyses across other omics modalities

**S1.8.1 Single-cell multiome analysis** We analyzed the GM12878 matched RNA-seq and ATAC-seq multiome dataset from Ma et al. [39], available under GSE140203 at https://www.ncbi.nlm.nih.gov/geo/query/acc.cgi?acc=GSE140203. We retained the top 5,000 highly variable RNA genes for analysis. RNA-based clustering followed the Seurat workflow described above, and downstream differential accessibility analysis was conducted on the matched ATAC modality. For Gene Ontology analysis, we used the intersection of genes nearest to differentially accessible regions and the top 2,000 highly variable RNA genes. The 10x Genomics PBMC 10k Multiome dataset was obtained from the SeuratData pbmcMultiome package (version 0.1.4), which contains paired scRNA-seq and scATAC-seq count matrices measured from the same cells. CD4 Naive and CD4 TCM cells were extracted, and differential accessibility analyses were performed on the ATAC modality.

**S1.8.2 Population-scale bulk transcriptomics analysis** For bulk microarray analysis, we used the glioblastoma dataset from Verhaak et al. [40], retaining the Proneural, Classical, and Mesenchymal subtypes; the data are available from https://gdc.cancer.gov/about-data/publications/gbm_exp. The original preprocessing retained genes with cross-platform correlation $\geq 0.7$, median absolute deviation (MAD) $> 0.5$ on high-correlation platforms, and a platform-to-unified MAD ratio $< 1.5$. Because of the modest sample size and concentration of variation in the leading components, the first two PCs were used for visualization. For bulk RNA-seq analysis, we used the TCGA ovarian cancer dataset analyzed by Luo et al. [41], available from https://portal.gdc.cancer.gov/. Clustering and differential testing followed the data-specific procedures described in ClusterDE Steps 2 and 3.

**S1.8.3 Microbiome analysis** We analyzed WGS microbiome profiles from thyroid cancer [42] and lung cancer [43], available from https://github.com/yge15/TCGA_Microbial_Content, together with the 16S rRNA vaginal microbiome dataset from Symul et al. [44], available from https://purl.stanford.edu/gp215vr4425. Potential contaminants were removed with GRIMER [45], and genera with two or fewer nonzero counts across all samples were excluded. Clustering and differential-abundance testing followed the procedures described in ClusterDE Steps 2 and 3.

### S2 Theoretical justification of ClusterDE

#### S2.1 Assumptions for FDR control of ClusterDE

In this section, we explain why ClusterDE can asymptotically control the false discovery rate (FDR) and avoid FDR inflation caused by double dipping. The theoretical justification is based on the results of the mirror statistics framework, which relies on data splitting [46]. Recall that ClusterDE leverages a multivariate count model [4] to generate synthetic null data and employs the $P$ value-free FDR control framework, Clipper [14], to identify DE genes.

To achieve FDR control in ClusterDE, the contrast scores $\{C_j : j = 1, \dots, m\}$ must satisfy the following assumptions. Recall that the contrast scores $\{C_j : j = 1, \dots, m\}$ are derived from the target DE scores $S_1, \dots, S_m$ and the null DE scores $\widetilde{S}_1, \dots, \widetilde{S}_m$, and are defined as:

$$C_j := S_j - \widetilde{S}_j \,.$$

Let $\mathcal{N} \subset \{1, \dots, m\}$ denote the set of true non-DE genes, and define $m_0 = \text{card}(\mathcal{N})$.

*Assumption 1.* We assume that the contrast scores $\{C_1, \dots, C_m\}$ are continuous random variables, and that there exists a sequence $a_m > 0$ such that for each truly non-DE gene $j \in \mathcal{N}$,

$$\mathbb{P}(C_j > t) \leq \mathbb{P}(C_j < -t) + a_m, \quad \text{for all } t > 0.$$

Assumption 1 requires $C_j$ to be left-skewed around 0, that is, its right tail is no heavier than the mirror of its left tail. Compared with the exact symmetry assumption around 0 used in data splitting [46], Assumption 1 is weaker and easier to satisfy, which makes it more suitable for realistic single-cell data.

*Assumption 2.* We assume that the contrast scores $\{C_j : j = 1, \dots, m\}$ are continuous random variables. Moreover, there exist constants $c > 0$ and $\alpha \in (0, 2)$ such that

$$\text{Var}\left(\sum_{j \in \mathcal{N}} \mathbb{I}(C_j > t)\right) \leq c m_0^{\alpha}, \quad \text{for all } t \in \mathbb{R}.$$

Assumption 2 only restricts the correlations among the non-DE genes. In one extreme scenario, if the non-DE genes are all independent, we have:

$$\text{Var}\left(\sum_{j \in \mathcal{N}} \mathbb{I}(C_j > t)\right) = \sum_{j \in \mathcal{N}} \text{Var}\left(\mathbb{I}(C_j > t)\right) \leq c m_0 \,,$$

where $c$ can be any constant no smaller than 1/4, the maximum variance of a binary random variable. In this case, $\alpha = 1$, and Assumption 2 holds. In the other extreme scenario, if all non-DE genes have identical contrast scores, then:

$$\text{Var}\left(\sum_{j \in \mathcal{N}} \mathbb{I}(C_j > t)\right) = \text{Var}\left(\mathbb{I}(C_j > t)\right) \cdot m_0^2 \,, \quad \forall j \in \mathcal{N} \,,$$

and Assumption 2 does not hold. More generally, if all contrast scores have constant pairwise correlation or can be clustered into a fixed number of groups with constant within-group correlation, then $\alpha$ must be 2, and Assumption 2 does not hold. Hence, Assumption 2 imposes restrictions on the extent and strength of correlations among the contrast scores of non-DE genes but is less stringent than assuming independence among these contrast scores. For instance, if only a small proportion of non-DE genes' contrast scores are correlated—a realistic scenario in biology—Assumption 2 may still hold.

##### S2.2 Theoretical justification for preserving gene-gene correlation in synthetic null data

This subsection presents a population-level theoretical justification for preserving gene–gene correlation in synthetic null data, which is essential to prevent false discoveries in post-clustering DE analysis. The sample-level argument is analogous but involves more technical considerations (e.g., error bounds), which we omit for clarity.

We begin by defining the data-generating model and its corresponding synthetic null model, assuming $G$ distinct cell types in the target dataset. Each cell type is modeled as a multivariate Gaussian distribution representing the log-normalized expression data, denoted by $\mathbf{E}$. The overall data-generating model is a mixture of these Gaussian components. The synthetic null model is defined as a single multivariate Gaussian distribution whose mean vector and covariance matrix match the first two moments of the mixture model.

**Definition 1 (Data generating model).** *The data-generating process is specified as a finite mixture of multivariate Gaussian distributions. Let* $\mathbf{E}$ *denote the random vector of log-normalized gene expression for a cell. Then*

$$\mathbf{E} \sim \sum_{g=1}^{G} \alpha_g \mathcal{N}(\boldsymbol{\mu}_g, \mathbf{R}_g),$$

*where* $\alpha_g \in [0,1]$ *with* $\sum_{g=1}^{G} \alpha_g = 1$ *are the mixing proportions,* $\boldsymbol{\mu}_g = (\mu_{g1}, \ldots, \mu_{gm})^\top$ *satisfies* $\sum_{g=1}^{G} \alpha_g \boldsymbol{\mu}_g = \mathbf{0}$, *and* $\mathbf{R}_g$ *denotes the gene–gene correlation matrix for cell type* $g$. *The synthetic null model is given by*

$$\mathbf{E}' \sim \mathcal{N}(\mathbf{0}, \mathbf{R}^\dagger),$$

*where* $\mathbf{R}^\dagger$ *is defined so that the covariance structure is preserved, i.e.,* $\mathrm{cov}(\mathbf{E}) = \mathrm{cov}(\mathbf{E}')$.

Let $\mathbf{E}_1, \ldots, \mathbf{E}_n$ denote $n$ i.i.d. samples from the true data-generating model, and let $\mathbf{E}'_1, \ldots, \mathbf{E}'_n$ denote $n$ i.i.d. samples from the corresponding synthetic null model. Following standard practice in single-cell data analysis, we first perform principal component analysis (PCA) separately on $\{\mathbf{E}_i\}_{i=1}^n$ and $\{\mathbf{E}'_i\}_{i=1}^n$, and then retain the top $J$ principal components (PCs) for clustering. Since the covariance matrices of $\mathbf{E}$ and $\mathbf{E}'$ are identical, the top $J$ PCs from $\{\mathbf{E}_i\}_{i=1}^n$ and $\{\mathbf{E}'_i\}_{i=1}^n$ asymptotically span the same subspace as $n \to \infty$. In other words, the clustering is asymptotically based on the same projected directions for both the target data and the synthetic null data. Based on this observation, we have the following proposition.

**Proposition 1.** *Under the data-generating model and the synthetic null model defined above, the projected directions used for clustering are asymptotically identical for the target data and the synthetic null data as* $n \to \infty$.

*Moreover, assuming a linear clustering boundary, if the target data consist of two distinct clusters and clustering is performed using the top principal component, then the decision boundaries for the target data and the synthetic null data are asymptotically identical up to an intercept shift as* $n \to \infty$:

$$\begin{cases} \mathbf{E}_i^\top \mathbf{e}_1(\boldsymbol{\Sigma}) \geq t_1 & \textit{assign label } 1, \\ \mathbf{E}_i^\top \mathbf{e}_1(\boldsymbol{\Sigma}) < t_1 & \textit{assign label } 0, \end{cases}$$

*and*

$$\begin{cases} (\mathbf{E}'_i)^\top \mathbf{e}_1(\boldsymbol{\Sigma}) \geq t_2 & \textit{assign label } 1, \\ (\mathbf{E}'_i)^\top \mathbf{e}_1(\boldsymbol{\Sigma}) < t_2 & \textit{assign label } 0, \end{cases}$$

*where* $\mathbf{e}_1(\boldsymbol{\Sigma})$ *denotes the leading eigenvector of* $\boldsymbol{\Sigma} = \mathrm{cov}(\mathbf{E}) = \mathrm{cov}(\mathbf{E}')$, *and* $t_1, t_2$ *are intercepts.*

Because the synthetic null data preserve the key distributional properties of the target data—including per-gene means, variances, and gene-gene correlations—the two datasets share the same covariance structure. Proposition 1 formalizes this intuition by showing that their leading principal components span the same subspace as $n \to \infty$. Consequently, when clustering is performed using the top principal component, the decision boundaries for the target and synthetic null data are asymptotically identical up to an intercept shift.

Intuitively, for a non-DE gene $j$, the marginal distribution of $E_j$ is the same as that of $E'_j$. Moreover, because the clustering boundaries are asymptotically identical, ClusterDE tends to satisfy Assumption 1, which in turn ensures valid FDR control.

##### S2.3 Theoretical justification for FDR control

Before establishing rigorous FDR control, we introduce notation that will be used throughout this subsection. Let

$$\mathcal{N} \subseteq \{1, \dots, m\}$$

denote the set of truly non-DE genes, and let

$$m_0 = \mathrm{card}(\mathcal{N}), \qquad m_1 = m - m_0, \qquad r_m = \frac{m_1}{m_0}.$$

For $t > 0$, define

$$\widehat{G}_m^0(t) = \frac{1}{m_0} \sum_{j \in \mathcal{N}} \mathbb{I}(C_j > t), \qquad G_m^0(t) = \frac{1}{m_0} \sum_{j \in \mathcal{N}} \mathbb{P}(C_j > t),$$

$$\widehat{G}_m^1(t) = \frac{1}{m_1} \sum_{j \notin \mathcal{N}} \mathbb{I}(C_j > t), \qquad \widehat{V}_m^0(t) = \frac{1}{m_0} \sum_{j \in \mathcal{N}} \mathbb{I}(C_j < -t).$$

The realized false discovery proportion at threshold $t$ can then be written as

$$\mathrm{FDP}_m(t) = \frac{\widehat{G}_m^0(t)}{\widehat{G}_m^0(t) + r_m \widehat{G}_m^1(t)}$$

whenever the denominator is positive, with the convention that the ratio is zero otherwise. We also define the null-only mirror estimate

$$\mathrm{FDP}_m^\dagger(t) = \frac{\widehat{V}_m^0(t)}{\widehat{G}_m^0(t) + r_m \widehat{G}_m^1(t)},$$

and its population-centered counterpart

$$\overline{\mathrm{FDP}}_m(t) = \frac{G_m^0(t)}{\widehat{G}_m^0(t) + r_m \widehat{G}_m^1(t)}.$$

The implementable FDP estimator used by ClusterDE is

$$\widehat{\mathrm{FDP}}_m(t) = \frac{\sum_{j=1}^m \mathbb{I}(C_j < -t)}{\sum_{j=1}^m \mathbb{I}(C_j > t) \vee 1}.$$

Since

$$\sum_{j \in \mathcal{N}} \mathbb{I}(C_j < -t) \le \sum_{j=1}^m \mathbb{I}(C_j < -t),$$

we have

$$\mathrm{FDP}_m^\dagger(t) \le \widehat{\mathrm{FDP}}_m(t)$$

whenever the common discovery count is positive.

Define the data-dependent threshold

$$\tau_q = \inf \left\{ t > 0 : \widehat{\mathrm{FDP}}_m(t) \le q \right\},$$

with the convention that $\tau_q = \infty$ if the set is empty.

**Lemma 1.** *Suppose Assumptions 1 and 2 hold, where the variance condition in Assumption 2 is imposed on both*

$$\sum_{j \in \mathcal{N}} \mathbb{I}(C_j > t) \quad \text{and} \quad \sum_{j \in \mathcal{N}} \mathbb{I}(C_j < -t).$$

*If $m_0 \to \infty$ and $a_m \to 0$ as $m \to \infty$, then*

$$\sup_{t>0} \left| \widehat{G}_m^0(t) - G_m^0(t) \right| = o_p(1),$$

*and*

$$\inf_{t>0} \left\{ \widehat{V}_m^0(t) - G_m^0(t) \right\} \ge -o_p(1).$$

*Proof.* Fix any $\epsilon \in (0,1)$. Because $G_m^0(t)$ is continuous and nonincreasing in $t$, there exist

$$-\infty = \alpha_0^m < \alpha_1^m < \cdots < \alpha_{N_\epsilon}^m = \infty, \qquad N_\epsilon = \left\lceil \frac{2}{\epsilon} \right\rceil,$$

such that

$$G_m^0(\alpha_{k-1}^m) - G_m^0(\alpha_k^m) \le \frac{\epsilon}{2}, \qquad k = 1, \ldots, N_\epsilon.$$

Since both $\widehat{G}_m^0(t)$ and $G_m^0(t)$ are nonincreasing, for $t \in [\alpha_{k-1}^m, \alpha_k^m)$,

$$\widehat{G}_m^0(t) - G_m^0(t) \le \widehat{G}_m^0(\alpha_{k-1}^m) - G_m^0(\alpha_{k-1}^m) + \frac{\epsilon}{2}.$$

Consequently, Assumption 2 and Chebyshev's inequality give

$$\begin{aligned}
&\mathbb{P}\left( \sup_{t\in\mathbb{R}} \left\{ \widehat{G}_m^0(t) - G_m^0(t) \right\} > \epsilon \right) \\
&\quad \le \sum_{k=1}^{N_\epsilon} \mathbb{P}\left( \widehat{G}_m^0(\alpha_{k-1}^m) - G_m^0(\alpha_{k-1}^m) > \frac{\epsilon}{2} \right) \\
&\quad \le \frac{4cN_\epsilon}{m_0^{2-\alpha}\epsilon^2} \longrightarrow 0.
\end{aligned}$$

A symmetric discretization argument yields

$$\mathbb{P}\left( \inf_{t\in\mathbb{R}} \left\{ \widehat{G}_m^0(t) - G_m^0(t) \right\} < -\epsilon \right) \longrightarrow 0.$$

This proves

$$\sup_{t>0} \left| \widehat{G}_m^0(t) - G_m^0(t) \right| = o_p(1).$$

Next, Assumption 1 implies that, for every $j \in \mathcal{N}$ and $t > 0$,

$$\mathbb{P}(C_j < -t) \ge \mathbb{P}(C_j > t) - a_m.$$

Therefore,

$$\frac{1}{m_0} \sum_{j\in\mathcal{N}} \mathbb{P}(C_j < -t) \ge G_m^0(t) - a_m.$$

Applying the same discretization and variance argument to the negative-tail indicators gives

$$\sup_{t>0} \left| \widehat{V}_m^0(t) - \frac{1}{m_0} \sum_{j\in\mathcal{N}} \mathbb{P}(C_j < -t) \right| = o_p(1).$$

Hence,

$$\inf_{t>0} \left\{ \widehat{V}_m^0(t) - G_m^0(t) \right\} \ge -a_m - o_p(1) = -o_p(1),$$

which proves the second claim.

**Theorem 1.** *Let $q \in (0,1)$. Suppose Assumptions 1 and 2 hold, $m_0 \to \infty$, and $a_m \to 0$ as $m \to \infty$. Assume that, for every sufficiently small $\epsilon \in (0,q)$, there exists a constant $t_{q-\epsilon} > 0$ such that*

$$\mathbb{P}\left\{ \mathrm{FDP}_m(t_{q-\epsilon}) \le q - \epsilon \right\} \longrightarrow 1.$$

*Further assume that the normalized discovery count is bounded away from zero on the relevant threshold range, in the sense that, for some constant $d_\epsilon > 0$,*

$$\mathbb{P}\left[ \inf_{0<t\le t_{q-\epsilon}} \left\{ \widehat{G}_m^0(t) + r_m \widehat{G}_m^1(t) \right\} \ge d_\epsilon \right] \longrightarrow 1.$$

*Then*

$$\mathrm{FDP}_m(\tau_q) \le q + o_p(1),$$

*and consequently,*

$$\limsup_{m\to\infty} \mathrm{FDR}_m(\tau_q) \le q.$$

*Proof.* By Lemma 1,

$$\sup_{0<t\le t_{q-\epsilon}} \left| \widehat{G}^0_m(t) - G^0_m(t) \right| = o_p(1),$$

and

$$\inf_{0<t\le t_{q-\epsilon}} \left\{ \widehat{V}^0_m(t) - G^0_m(t) \right\} \ge -o_p(1).$$

Together with the lower bound on the normalized discovery count, these results imply

$$\sup_{0<t\le t_{q-\epsilon}} \left\{ \mathrm{FDP}_m(t) - \mathrm{FDP}^\dagger_m(t) \right\} \le o_p(1).$$

Thus, uniformly over the relevant threshold range, the null-only mirror estimate asymptotically upper-bounds the realized FDP.

At $t = t_{q-\epsilon}$, we have

$$\mathrm{FDP}^\dagger_m(t_{q-\epsilon}) \le \mathrm{FDP}_m(t_{q-\epsilon}) + o_p(1) \le q - \epsilon + o_p(1).$$

Moreover,

$$\mathrm{FDP}^\dagger_m(t) \le \widehat{\mathrm{FDP}}_m(t)$$

because the numerator of $\widehat{\mathrm{FDP}}_m(t)$ includes negative contrast scores from both null and nonnull genes. To guarantee that the implementable estimate crosses the target level, we additionally require

$$\mathbb{P}\left\{ \widehat{\mathrm{FDP}}_m(t_{q-\epsilon}) \le q \right\} \longrightarrow 1.$$

It then follows from the definition of $\tau_q$ that

$$\mathbb{P}\left(\tau_q \le t_{q-\epsilon}\right) \longrightarrow 1.$$

On the event $\{\tau_q \le t_{q-\epsilon}\}$, uniform convergence gives

$$\mathrm{FDP}_m(\tau_q) \le \mathrm{FDP}^\dagger_m(\tau_q) + o_p(1).$$

Since

$$\mathrm{FDP}^\dagger_m(\tau_q) \le \widehat{\mathrm{FDP}}_m(\tau_q) \le q,$$

we obtain

$$\mathrm{FDP}_m(\tau_q) \le q + o_p(1).$$

Finally, because

$$0 \le \mathrm{FDP}_m(\tau_q) \le 1,$$

boundedness and convergence in probability imply

$$\limsup_{m\to\infty} \mathbb{E}\left[\mathrm{FDP}_m(\tau_q)\right] \le q.$$

Therefore,

$$\limsup_{m\to\infty} \mathrm{FDR}_m(\tau_q) \le q.$$

#### S3 Supplementary Figures

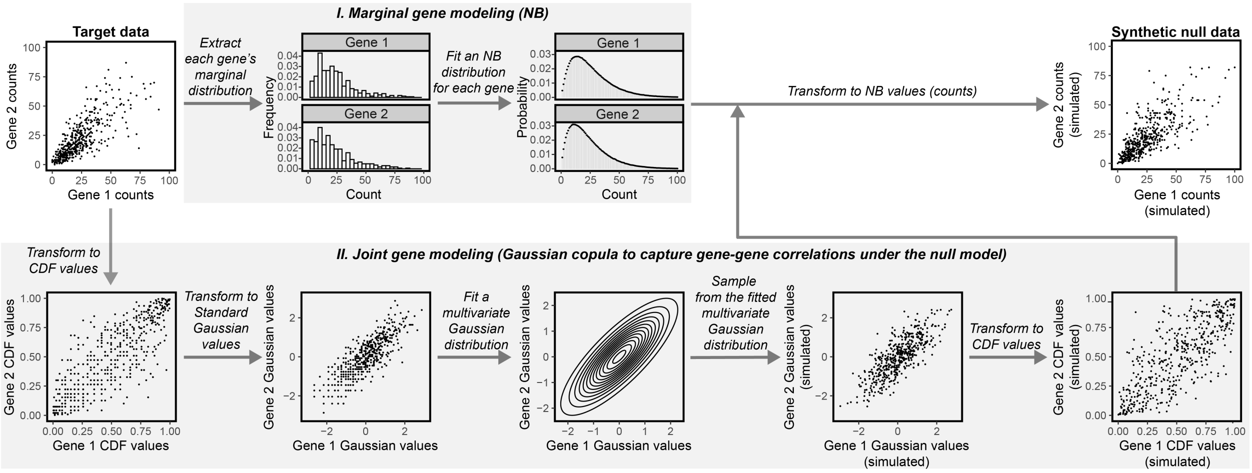


Fig. S1: **Overview of the synthetic null data generation procedure for scRNA-seq in ClusterDE.** The process for generating synthetic null data from target scRNA-seq data (top left). For illustration, we show the bivariate case (two genes), whereas the actual case involves high-dimensional data with thousands of genes. The null model consists of two components: **marginal gene modeling** and **joint gene modeling**. For marginal gene modeling (top), each gene's counts are modeled using a negative binomial (NB) distribution. The two NB parameters (mean and dispersion) are estimated from the target data for each gene. For joint gene modeling part (bottom), there are three steps. (1) *Transformation to cumulative distribution function (CDF) values*: Each gene's counts in the target data are transformed into CDF values using either the fitted NB distribution or the empirical distribution (if the target data contains a large number of cells), so the gene's CDF values are uniform between 0 and 1. (2) *Transformation to Gaussian values*: Each gene's CDF values are transformed into quantiles of the standard Gaussian distribution, $N(0,1)$. (3) *Modeling gene dependence*: A multivariate Gaussian distribution (illustrated here as a bivariate Gaussian) is fitted to the transformed Gaussian values of the genes whose correlations are to be modeled. The correlation matrix of this fitted multivariate Gaussian distribution specifies the Gaussian copula, capturing the gene dependence structure.

After the marginal and joint distributions are modeled, synthetic counts are generated in three steps. (1) *Sampling Gaussian values*: Standard Gaussian values for genes are jointly sampled from the fitted multivariate Gaussian distribution. (2) *Transformation to CDF values*: The sampled Gaussian values are transformed into CDF values of the standard Gaussian distribution. (3) *Transformation to counts*: The CDF values are transformed into quantiles of the fitted NB distribution for each gene, becoming synthetic counts that constitute the synthetic null data (top right).

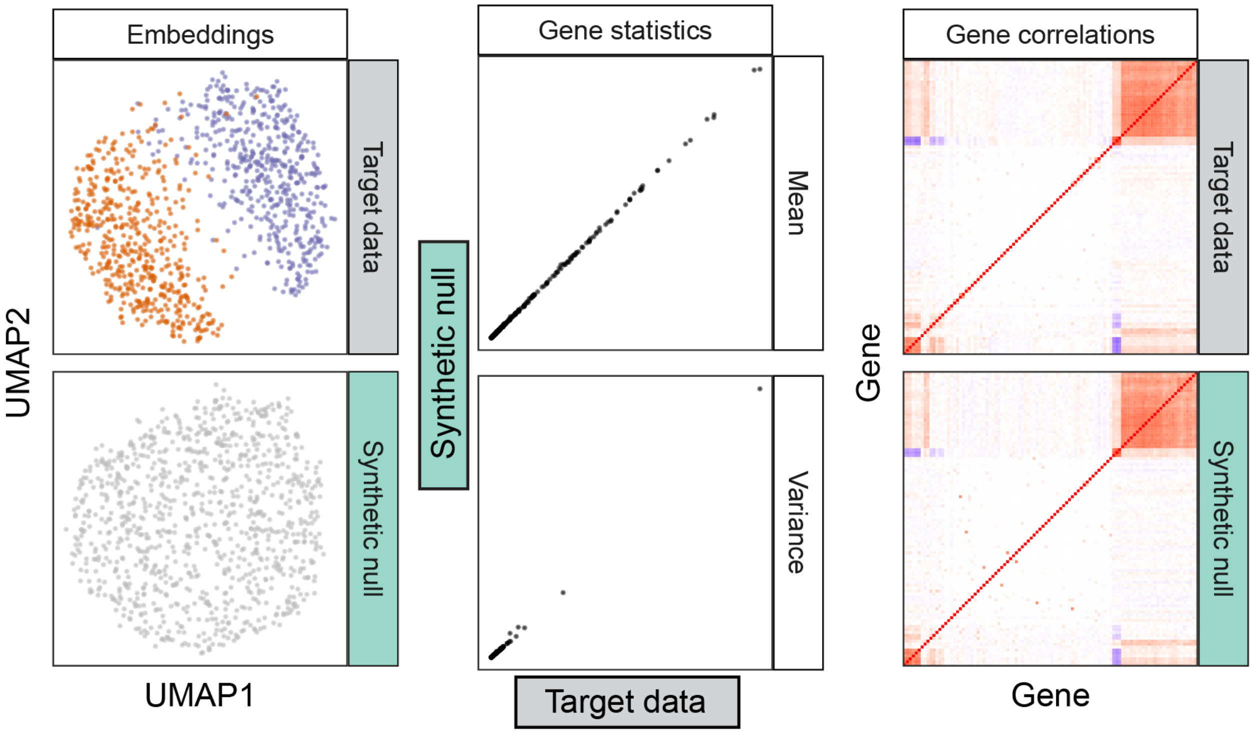


Fig. S2: **ClusterDE generates synthetic null scRNA-seq data that resemble two–cell-type target data while eliminating between-type separation.** When the target scRNA-seq data contains cells from two cell types (simulation; see "Simulation setting with two cell types and 200 true DE genes" in the Supplementary Methods), the synthetic null data generated by ClusterDE fills the gap between the two cell types but resembles the target data in other visual aspects of UMAP cell embeddings (left), per-gene expression mean and variance statistics (middle), and gene-gene rank correlations (right).

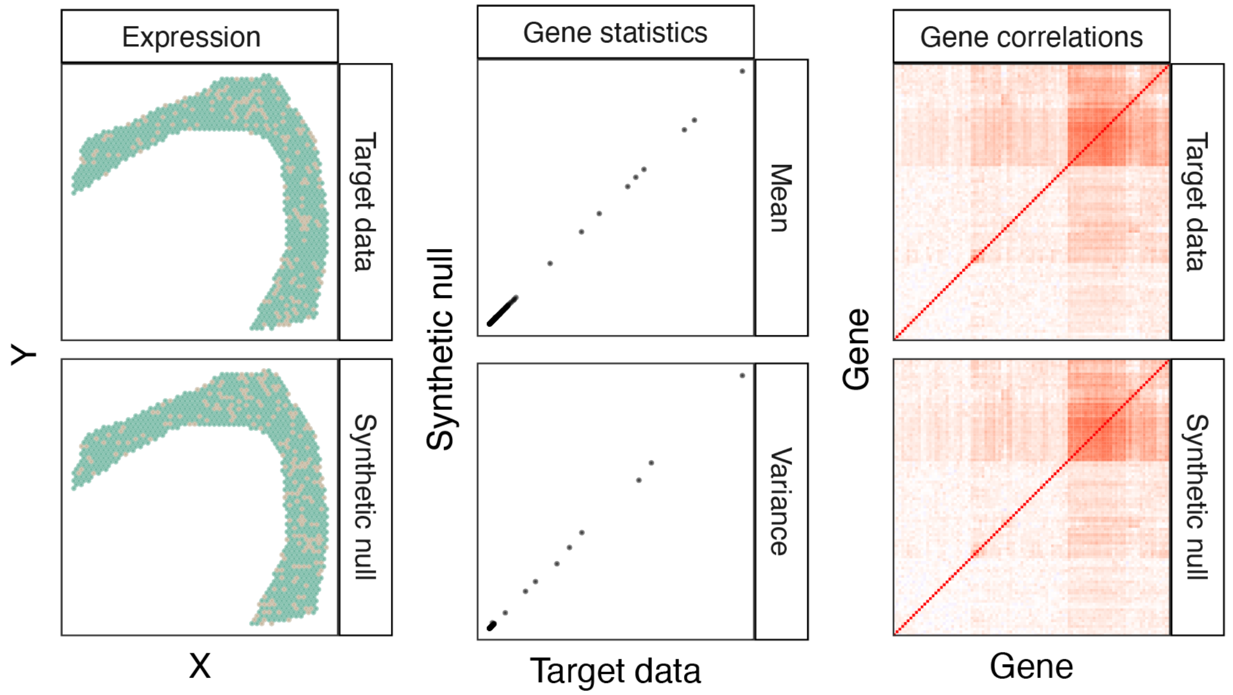


Fig. S3: **ClusterDE generates synthetic null SRT data that resemble one-domain target data while preserving key statistical properties.** When the target SRT data contains one spatial domain (simulation; see "Simulation setting with one spatial domain" in the Supplementary Methods), the synthetic null data generated by ClusterDE resembles the target data (left) while preserving per-gene expression mean and variance statistics (middle) as well as gene-gene rank correlations (right).

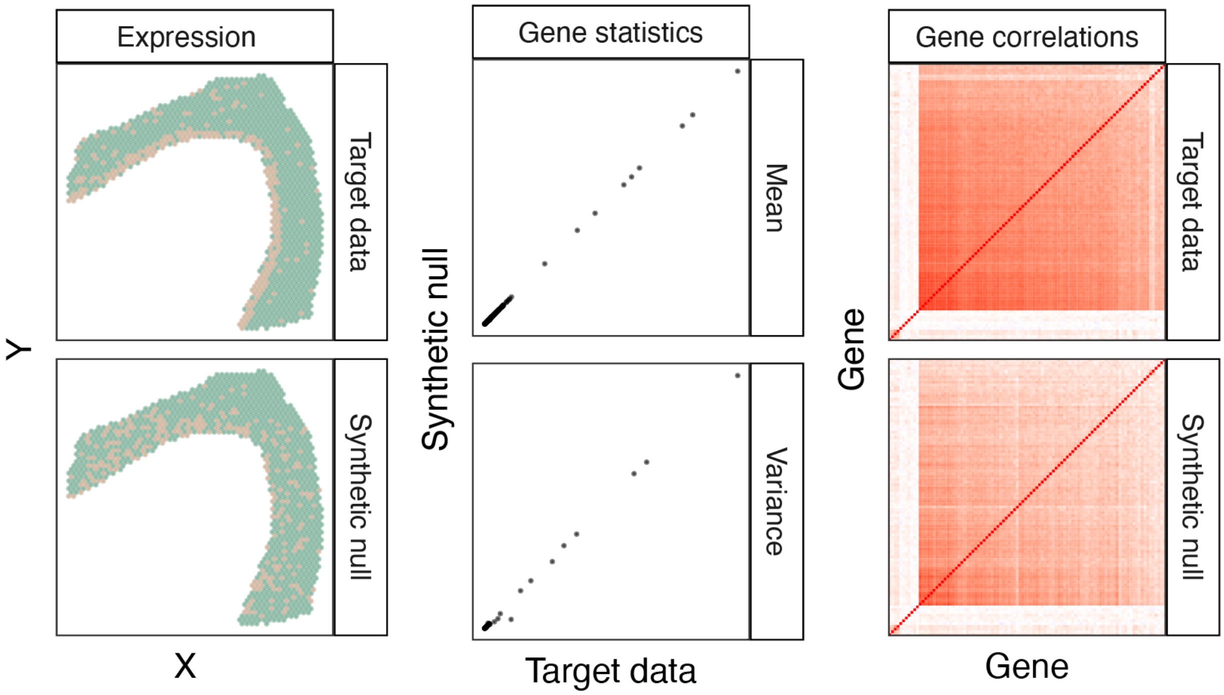


Fig. S4: **ClusterDE generates synthetic null SRT data that blur domain boundaries while preserving key statistical properties.** When the target SRT data contains two spatial domains (simulation; see "Simulation setting with two spatial domains and 200 true spatial-domain marker genes" in the Supplementary Methods), the synthetic null data generated by ClusterDE blurs the domain boundary (left) while preserving per-gene expression mean and variance statistics (middle) as well as gene-gene rank correlations (right).

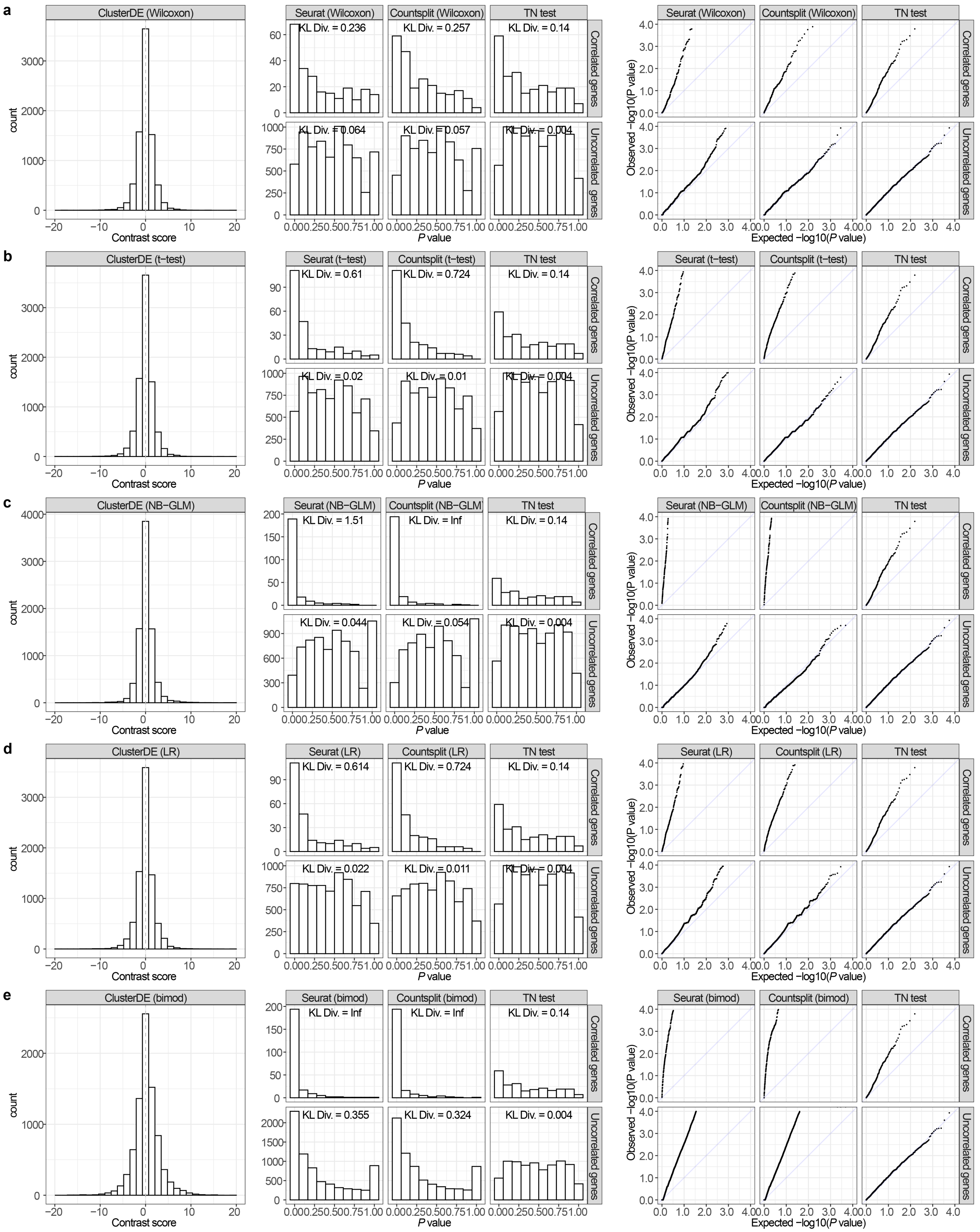
a
ClusterDE (Wilcoxon)
Seurat (Wilcoxon)
Countsplit (Wilcoxon)
TN test
KL Div. = 0.236
KL Div. = 0.257
KL Div. = 0.14
KL Div. = 0.064
KL Div. = 0.057
KL Div. = 0.004
b
ClusterDE (t−test)
Seurat (t−test)
Countsplit (t−test)
TN test
KL Div. = 0.61
KL Div. = 0.724
KL Div. = 0.14
KL Div. = 0.02
KL Div. = 0.01
KL Div. = 0.004
c
ClusterDE (NB−GLM)
Seurat (NB−GLM)
Countsplit (NB−GLM)
TN test
KL Div. = 1.51
KL Div. = Inf
KL Div. = 0.14
KL Div. = 0.044
KL Div. = 0.054
KL Div. = 0.004
d
ClusterDE (LR)
Seurat (LR)
Countsplit (LR)
TN test
KL Div. = 0.614
KL Div. = 0.724
KL Div. = 0.14
KL Div. = 0.022
KL Div. = 0.011
KL Div. = 0.004
e
ClusterDE (bimod)
Seurat (bimod)
Countsplit (bimod)
TN test
KL Div. = Inf
KL Div. = Inf
KL Div. = 0.14
KL Div. = 0.355
KL Div. = 0.324
KL Div. = 0.004
Correlated genes
Uncorrelated genes
count
Contrast score
P value
Observed −log10(P value)
Expected −log10(P value)

Fig. S5: **Validity checks of the contrast scores of ClusterDE and $P$ values of Seurat, Countsplit, and the TN test on an exemplary one-cell-type dataset.** In this dataset, no post-clustering DE genes are true cell-type markers by simulation design (see "Simulation setting with one cell type" in the Supplementary Methods). The panels (rows) **a–e** represent the five DE tests in Seurat used in ClusterDE, Seurat, and Countsplit (see "ClusterDE step 3: DE analysis" in the Supplementary Material); since the TN test has its own DE test, its results are the same in the panels **a–e**. The first column shows that the ClusterDE contrast scores for all genes (true non-marker genes) are approximately symmetric around 0, which meets the assumption of ClusterDE for the FDR control. The second column shows the histograms of the $P$ values of the correlated genes (top) and the uncorrelated genes (bottom) from Seurat, Countsplit, and the TN test. A larger empirical Kullback-Leibler divergence (KL div.) between the $P$ value distribution and the theoretical Uniform$[0, 1]$ distribution represents a more severe violation of the $P$ value uniformity assumption. The results show that Countsplit (for four out of the five DE tests) and the TN test have close-to-uniform $P$ values for the uncorrelated genes, but their $P$ values exhibit a severe departure from the uniform distribution for the correlated genes. The third column contains the quantile-quantile plots of the negative log-transformed $P$ values corresponding to the second column.

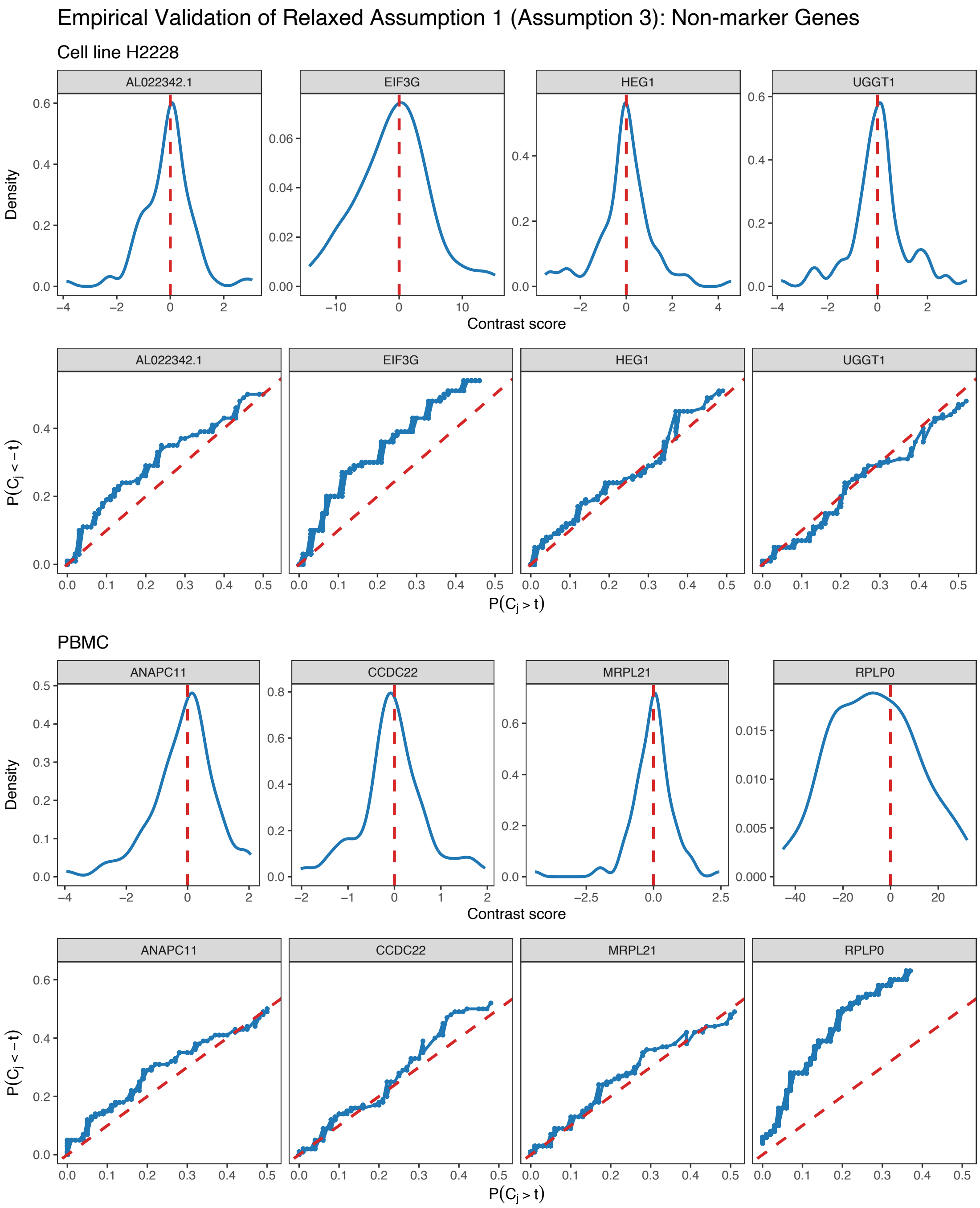


Fig. S6: **Left-skewness of contrast-score distributions: empirical checks.** Empirical validation of Relaxed Assumption 1 (Assumption 3)—left-skewness of the contrast-score distribution—using (i) four randomly selected non-marker genes from the H2228 cell-line dataset and (ii) four randomly selected housekeeping genes from the $CD4^+$ T vs. cytotoxic T comparison in the PBMC dataset.

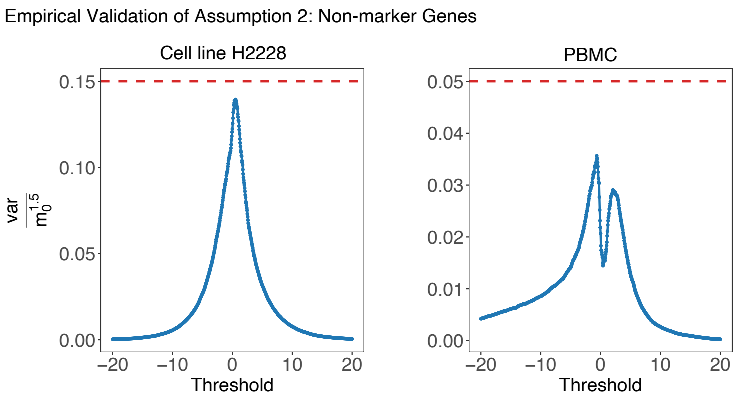


Fig. S7: **Weak dependence condition: empirical checks.** Empirical validation of Assumption 2—weak dependence condition: empirical checks—from the H2228 cell-line dataset and the CD4$^+$ T vs. cytotoxic T comparison in the PBMC dataset.

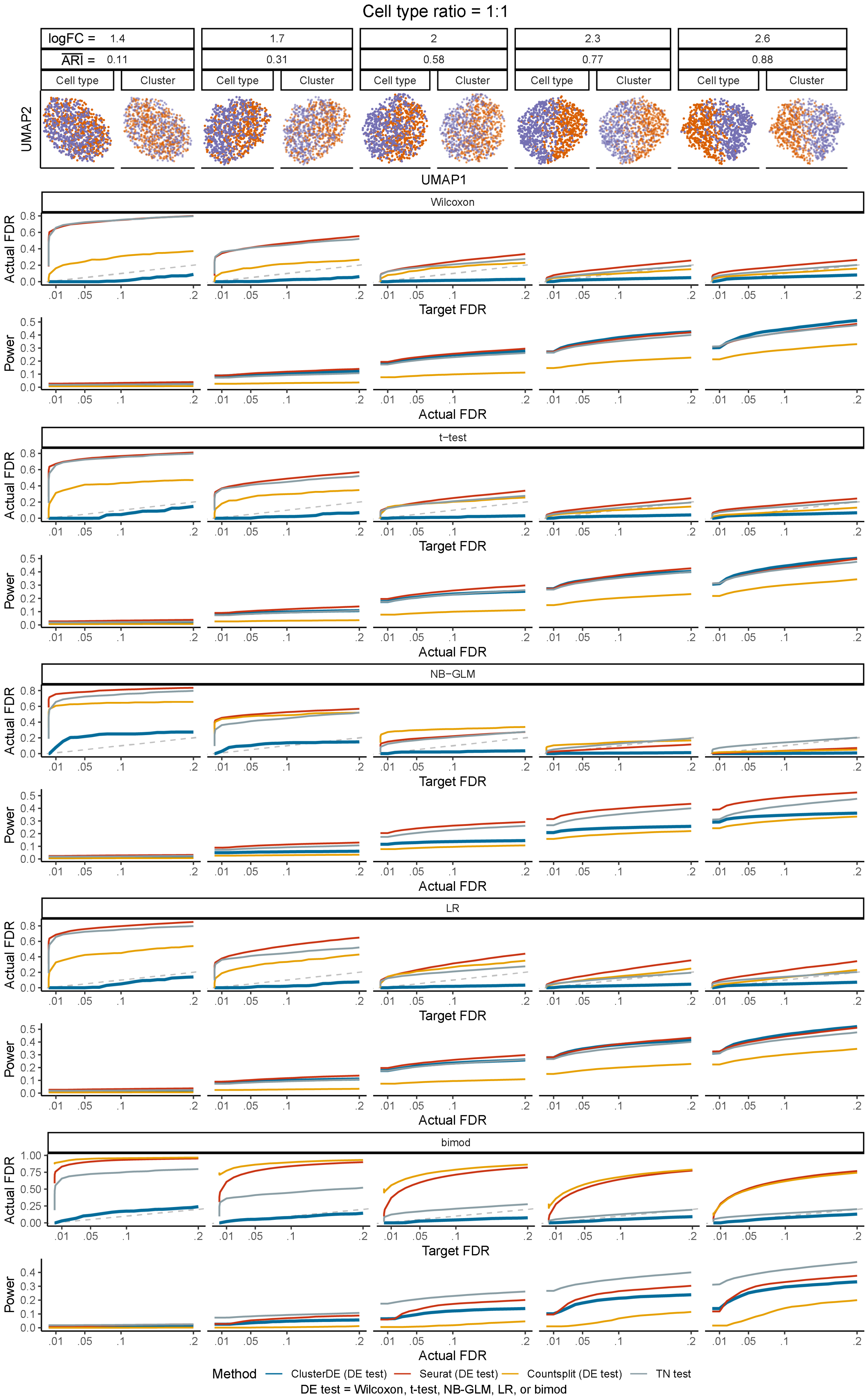
Cell type ratio = 1:1
logFC = 1.4
1.7
2
2.3
2.6
ARI = 0.11
0.31
0.58
0.77
0.88
Cell type
Cluster
UMAP2
UMAP1
Wilcoxon
t−test
NB−GLM
LR
bimod
Actual FDR
Target FDR
Power
Method
ClusterDE (DE test)
Seurat (DE test)
Countsplit (DE test)
TN test
DE test = Wilcoxon, t-test, NB-GLM, LR, or bimod

Fig. S8: **Comparison of the FDRs and power of ClusterDE with three existing methods (Seurat, Countsplit, and the TN test) under varying levels of double-dipping severity when the two cell types have a size ratio of** $1:1$**.** The log fold change (logFC) represents the average gene expression difference between the two cell types in the simulation (see "Simulation setting with two cell types and 200 true DE genes" in the Supplementary Methods). A small logFC leads to a small adjusted Rand index (ARI), indicating poor agreement between cell clusters and cell types and representing a more severe double-dipping issue. Across different levels of double-dipping severity and the five DE tests, ClusterDE consistently controls the FDRs below the target thresholds (gray diagonal dashed line) and, under the default Wilcoxon rank-sum test, achieves comparable or higher power relative to the three existing methods at equivalent actual FDR levels.

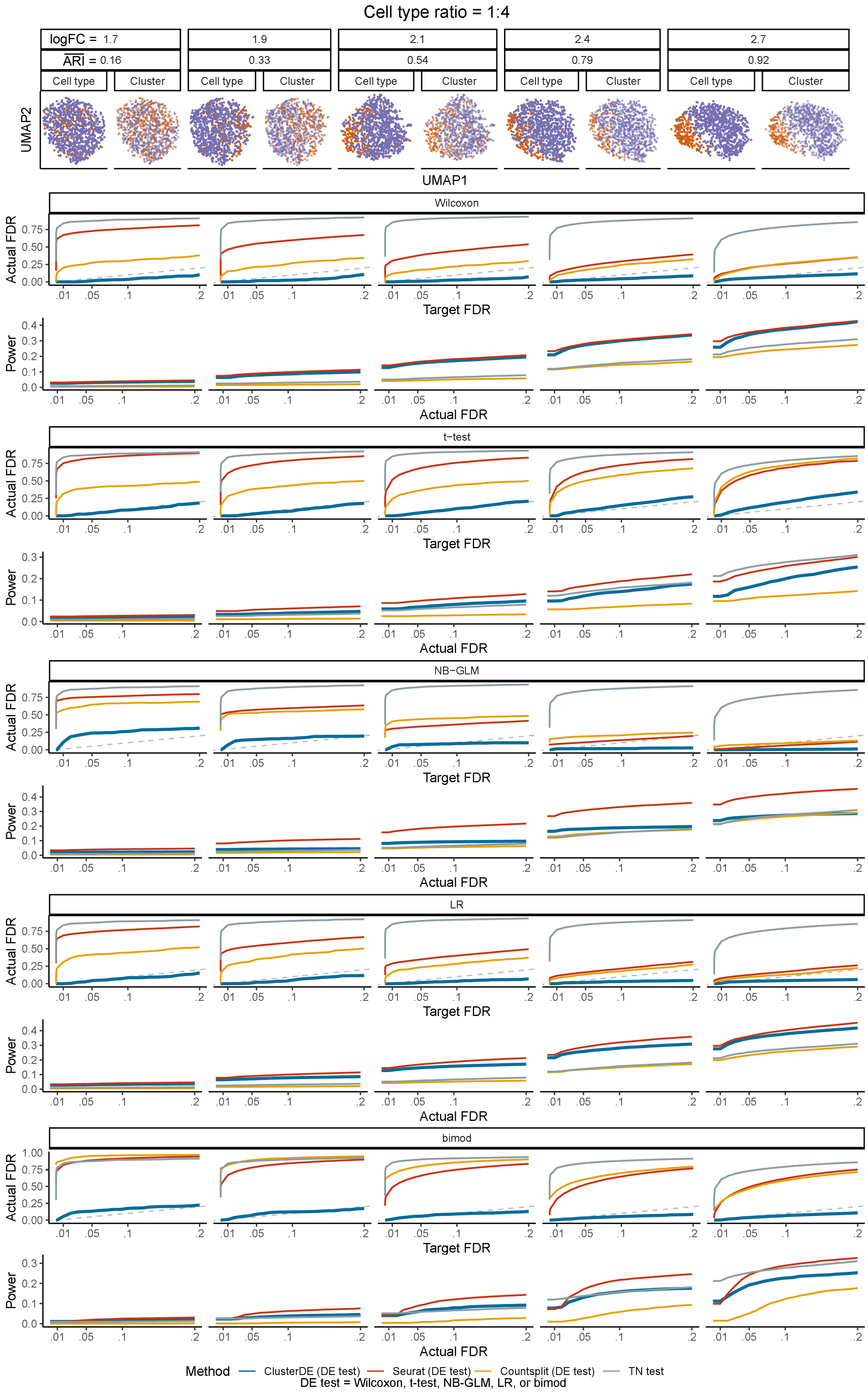

Cell type ratio = 1:4
logFC = 1.7
1.9
2.1
2.4
2.7
ARI = 0.16
0.33
0.54
0.79
0.92
Cell type
Cluster
UMAP2
UMAP1
Wilcoxon
t−test
NB−GLM
LR
bimod
Actual FDR
Target FDR
Power
Method
ClusterDE (DE test)
Seurat (DE test)
Countsplit (DE test)
TN test
DE test = Wilcoxon, t-test, NB-GLM, LR, or bimod

Fig. S9: **Comparison of the FDRs and power of ClusterDE with three existing methods (Seurat, Countsplit, and the TN test) under varying levels of double-dipping severity when the two cell types have a size ratio of** $1:4$**.** The log fold change (logFC) represents the average gene expression difference between the two cell types in the simulation (see "Simulation setting with two cell types and 200 true DE genes" in the Supplementary Methods). A small logFC leads to a small adjusted Rand index (ARI), indicating poor agreement between cell clusters and cell types and representing a more severe double-dipping issue. Across different levels of double-dipping severity and the five DE tests, ClusterDE consistently controls the FDRs below the target thresholds (gray diagonal dashed line) and, under the default Wilcoxon rank-sum test, achieves comparable or higher power relative to the three existing methods at equivalent actual FDR levels.

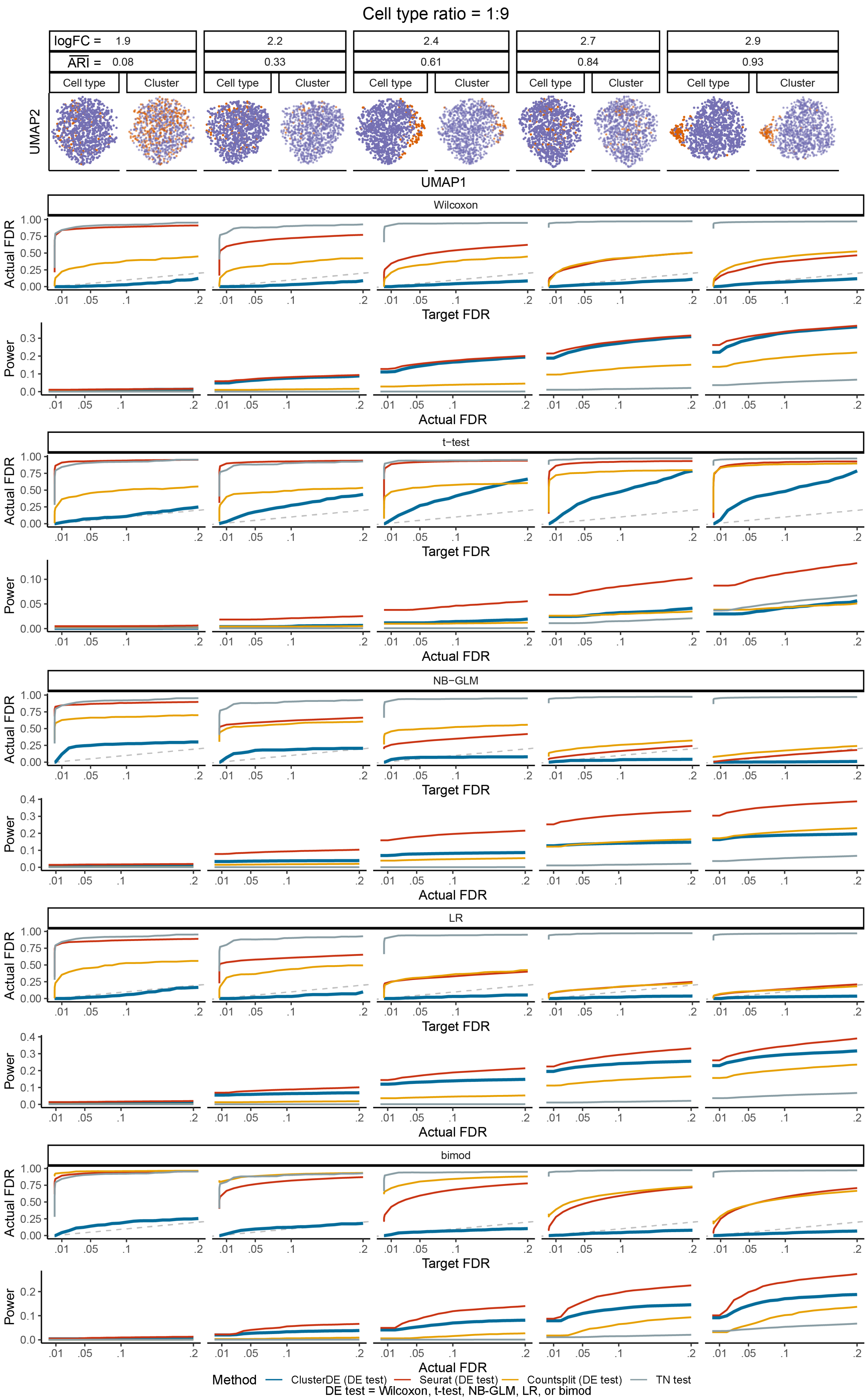
Cell type ratio = 1:9
logFC = 1.9
2.2
2.4
2.7
2.9
ARI = 0.08
0.33
0.61
0.84
0.93
Cell type
Cluster
UMAP2
UMAP1
Wilcoxon
t−test
NB−GLM
LR
bimod
Actual FDR
Target FDR
Power
Method
ClusterDE (DE test)
Seurat (DE test)
Countsplit (DE test)
TN test
DE test = Wilcoxon, t-test, NB-GLM, LR, or bimod

Fig. S10: **Comparison of the FDRs and power of ClusterDE with three existing methods (Seurat, Countsplit, and the TN test) under varying levels of double-dipping severity when the two cell types have a size ratio of** $1:9$**.** The log fold change (logFC) represents the average gene expression difference between the two cell types in the simulation (see "Simulation setting with two cell types and 200 true DE genes" in the Supplementary Methods). A small logFC leads to a small adjusted Rand index (ARI), indicating poor agreement between cell clusters and cell types and representing a more severe double-dipping issue. Across different levels of double-dipping severity and the five DE tests, ClusterDE consistently controls the FDRs below the target thresholds (gray diagonal dashed line) and, under the default Wilcoxon rank-sum test, achieves comparable or higher power relative to the three existing methods at equivalent actual FDR levels.

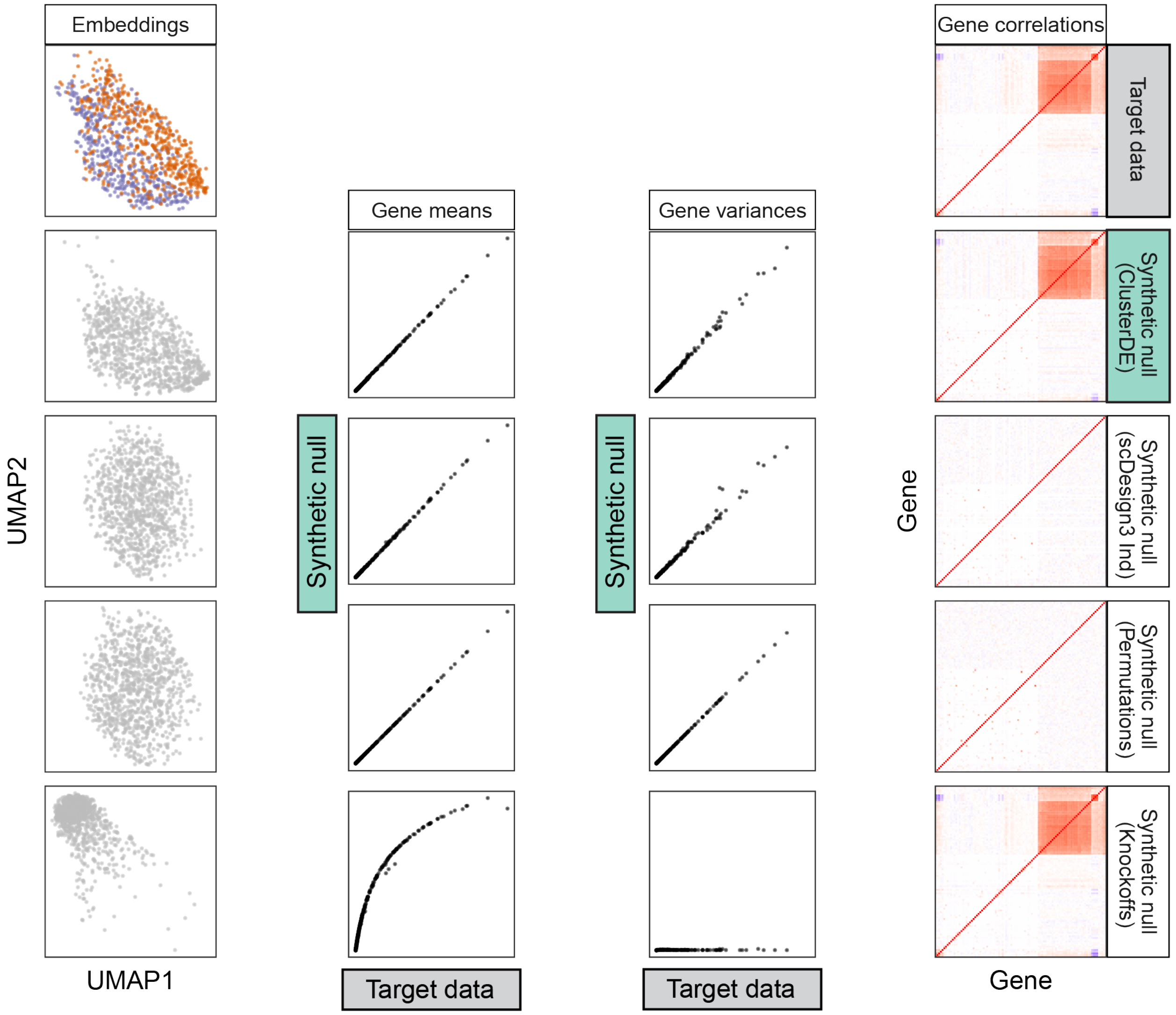


Fig. S11: **ClusterDE generates synthetic null data that more closely resembles the target data than three alternative strategies.** When the target data contains cells from two cell types (simulation; see "Simulation setting with two cell types and 200 true DE genes" in the Supplementary Methods), the synthetic null data generated by ClusterDE (second row) effectively fills the gap between the two cell types while closely resembling the target data in other visual aspects of UMAP cell embeddings (left), per-gene expression mean and variance statistics (middle), and gene-gene rank correlations (right). In contrast, the synthetic null data generated by scDesign3 without copula modeling ("scDesign3 Ind," third row), permutations (fourth row) and the model-X knockoffs (fifth row) do not resemble the target data. Specifically, the synthetic null data generated by the model-X knockoffs preserves the gene-gene correlations of the target data but does not preserve per-gene expression mean and variance statistics. Conversely, the synthetic null data generated by scDesign3 Ind and permutations preserves per-gene expression mean and variance statistics but does not preserve gene-gene rank correlations.

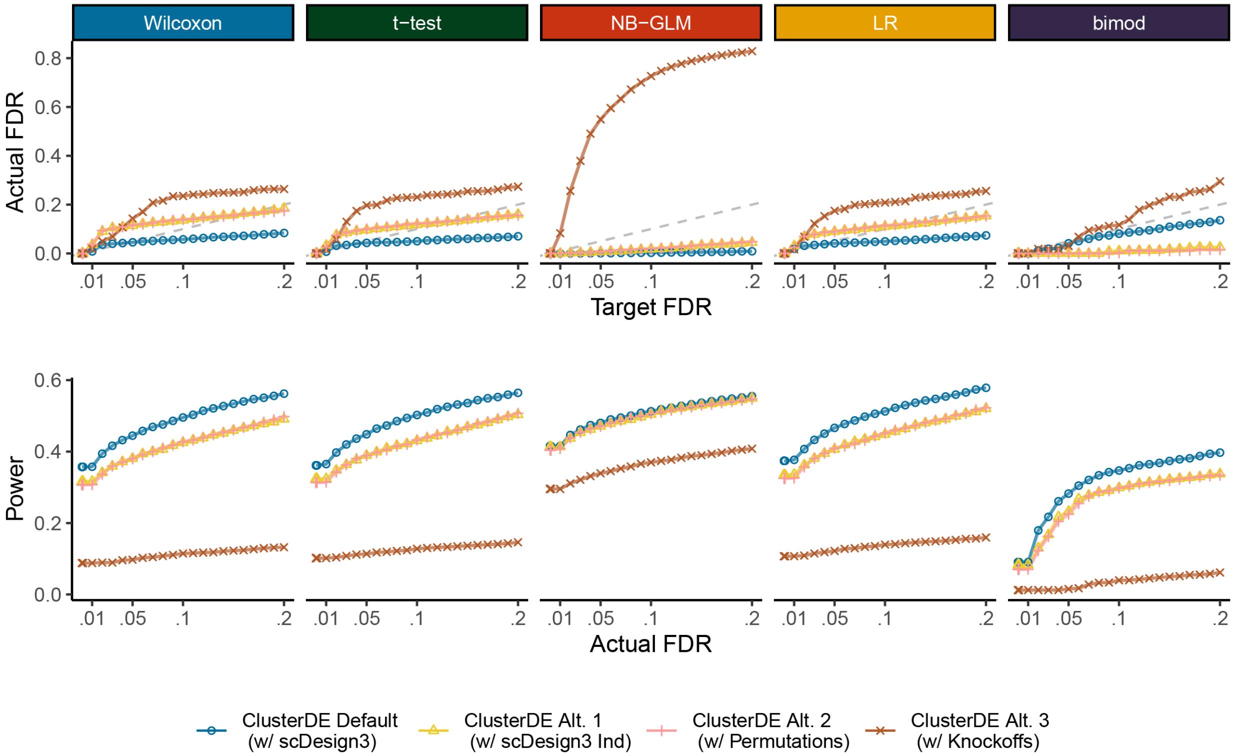


Fig. S12: **Comparison of the FDRs and power of ClusterDE with four strategies for synthetic null data generation in Step 1.** The four strategies include scDesign3 (the default in ClusterDE), scDesign3 Ind (without copula modeling, assuming gene independence), independent permutations of all genes across cells, and the model-X knockoffs. The default ClusterDE, employing scDesign3, demonstrates superior performance by effectively controlling the FDR and achieving higher power compared to the three alternative strategies.

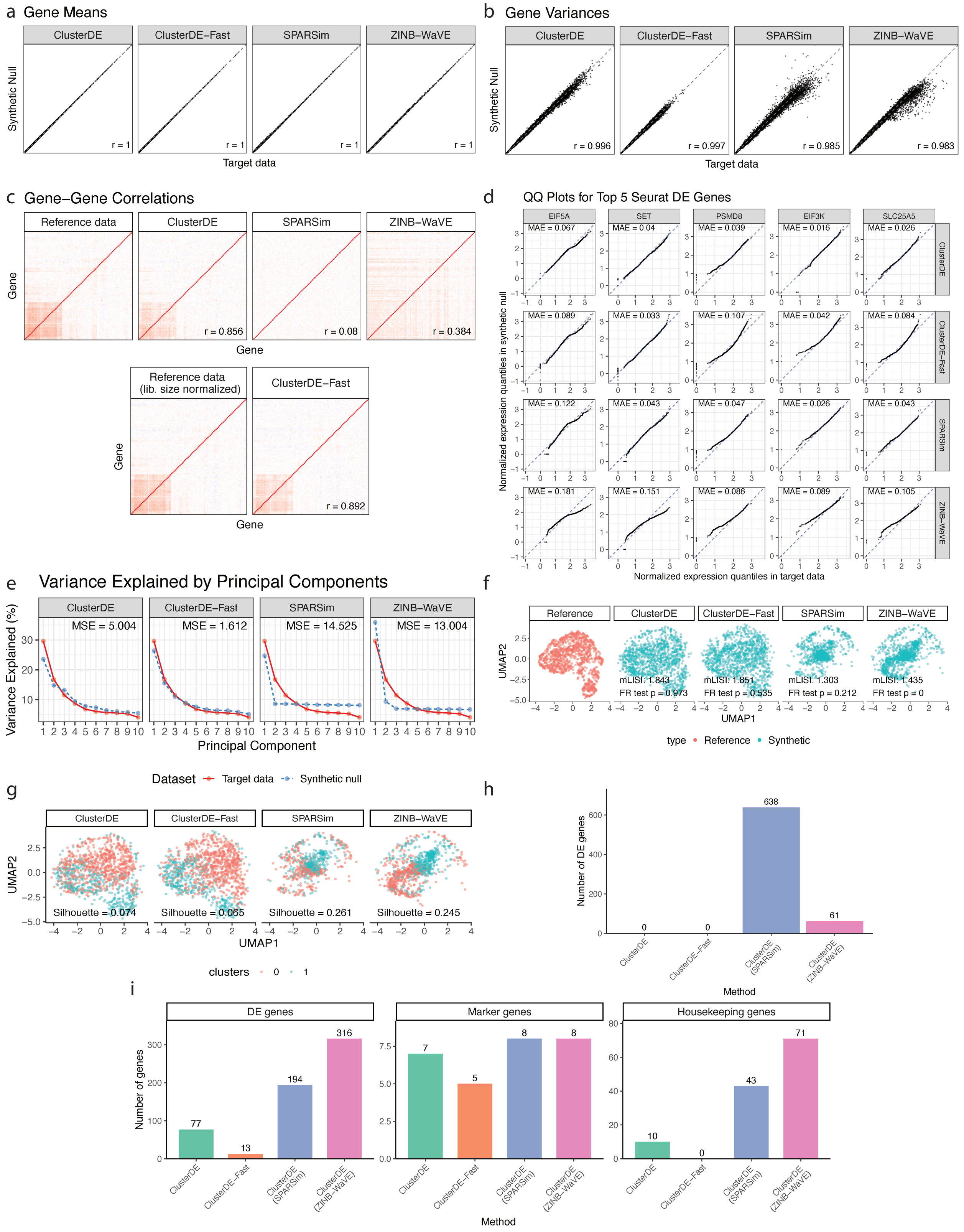
a Gene Means
ClusterDE
ClusterDE–Fast
SPARSim
ZINB–WaVE
Synthetic Null
r = 1
Target data
b Gene Variances
r = 0.996
r = 0.997
r = 0.985
r = 0.983
c Gene–Gene Correlations
Reference data
Gene
r = 0.856
r = 0.08
r = 0.384
Reference data (lib. size normalized)
r = 0.892
d QQ Plots for Top 5 Seurat DE Genes
EIF5A
SET
PSMD8
EIF3K
SLC25A5
MAE = 0.067
MAE = 0.04
MAE = 0.039
MAE = 0.016
MAE = 0.026
MAE = 0.089
MAE = 0.033
MAE = 0.107
MAE = 0.042
MAE = 0.084
MAE = 0.122
MAE = 0.043
MAE = 0.047
MAE = 0.026
MAE = 0.043
MAE = 0.181
MAE = 0.151
MAE = 0.086
MAE = 0.089
MAE = 0.105
Normalized expression quantiles in synthetic null
Normalized expression quantiles in target data
e Variance Explained by Principal Components
Variance Explained (%)
MSE = 5.004
MSE = 1.612
MSE = 14.525
MSE = 13.004
Principal Component
Dataset
Target data
Synthetic null
f
Reference
UMAP2
mLISI: 1.843
FR test p = 0.973
mLISI: 1.851
FR test p = 0.535
mLISI: 1.303
FR test p = 0.212
mLISI: 1.435
FR test p = 0
UMAP1
type
Synthetic
g
Silhouette = 0.074
Silhouette = 0.065
Silhouette = 0.261
Silhouette = 0.245
clusters
0
1
h
Number of DE genes
638
0
0
61
ClusterDE (SPARSim)
ClusterDE (ZINB–WaVE)
Method
i
DE genes
Marker genes
Housekeeping genes
Number of genes
77
13
194
316
7
5
8
8
10
0
43
71

Fig. S13: **Built-in quality checks for existing simulators that can generate null data. a**, For the A549 cell line data, the synthetic null data generated by ClusterDE, ClusterDE-Fast, SPARSim, and ZINB-WaVE resembled the target data well in terms of per-gene mean expression and **b**, variance. **c**, The synthetic null data generated by ClusterDE and ClusterDE-Fast also resembled the target data well in terms of gene–gene correlations. In contrast, the synthetic null data from SPARSim and ZINB-WaVE did not capture the gene–gene correlations of the target data. **d**, QQ plots comparing normalized expression distributions between the target and synthetic data for the top five Seurat-identified DE genes in the target data. The mean absolute error (MAE) between the corresponding quantiles is shown in each panel. **e**, Percentage of variance explained by the first 10 principal components in the target and synthetic data. The mean squared error (MSE) between the corresponding variance-explained profiles is shown for each method. **f**, The synthetic null data generated by ClusterDE and ClusterDE-Fast resembled the target data well in terms of UMAP cell embeddings, whereas SPARSim and ZINB-WaVE did not capture the global cell topology of the target data. UMAP mixing was evaluated using the mean local inverse Simpson's index (mLISI), which quantifies local mixing between target and synthetic cells in the UMAP space. Separately, the Friedman–Rafsky (FR) test was used to assess whether target and synthetic cells follow the same multivariate distribution based on normalized expression of the highly variable genes identified by Seurat in the target data. **g**, Default Seurat clustering was applied to each of the four synthetic null datasets to obtain two clusters, and the silhouette score was computed using Euclidean distances in the space of the first 50 PCs. **h**, Number of DE genes identified by ClusterDE with the Wilcoxon test using each of the four synthetic null datasets for the A549 cell line data. **i**, Number of DE genes (left), number of known marker genes among the DE genes (middle), and number of housekeeping genes among the DE genes (right) identified by ClusterDE with the Wilcoxon test using each of the four synthetic null datasets for the Rep1_10x(v3) dataset.

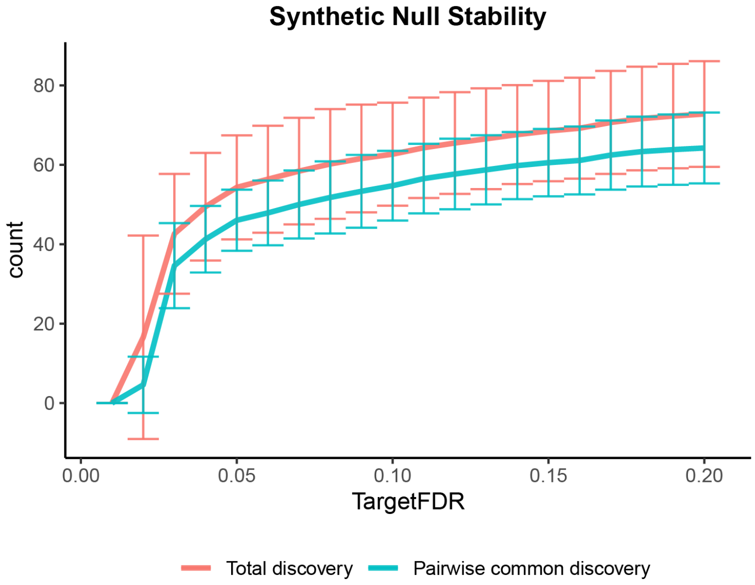


Fig. S14: **Stability of the DE genes identified by ClusterDE with respect to the randomness of synthetic null data generation.** Using a target dataset simulated with two cell types (see "Simulation setting with two cell types and 200 true DE genes" in the Supplementary Methods), 50 synthetic null datasets are generated with 50 random seeds, and DE genes are identified by ClusterDE at varying target FDRs for each synthetic null dataset. The red curve represents the mean number of DE genes identified at each target FDR, with the standard deviation shown as half the height of the vertical error bars, calculated across the 50 random seeds. The cyan curve represents the mean number of DE genes shared between results from two random seeds, with the standard deviation (half the height of the vertical error bars) calculated across all $\binom{50}{2}$ pairs of random seeds at each target FDR. The results indicate that DE genes identified by ClusterDE are relatively stable and robust to the randomness of synthetic null data generation.

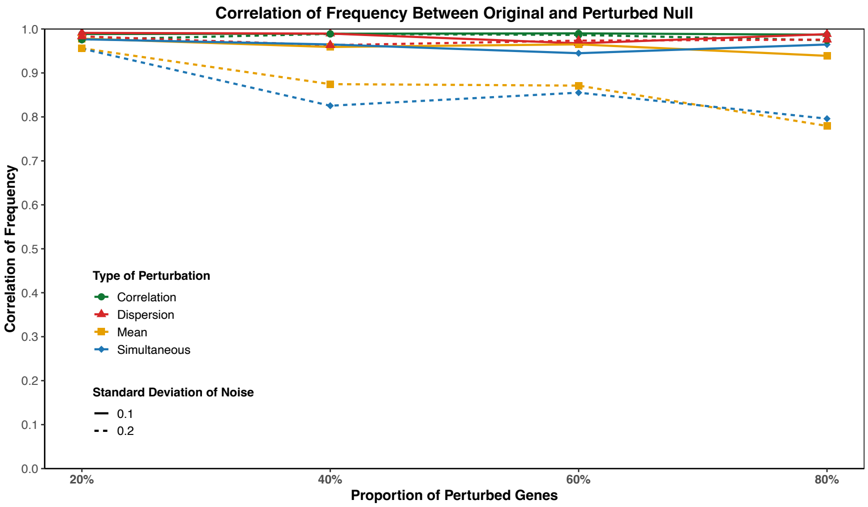


Fig. S15: **Robustness of ClusterDE-Stable to perturbations of the synthetic null.** Each point shows the correlation between gene-wise selection frequencies obtained by ClusterDE-Stable under the original (unperturbed) null and a perturbed null. We perturb the null model's key parameters—means, dispersions, and gene–gene correlations—separately and jointly, across varying proportions of perturbed genes and two noise magnitudes (standard deviations).

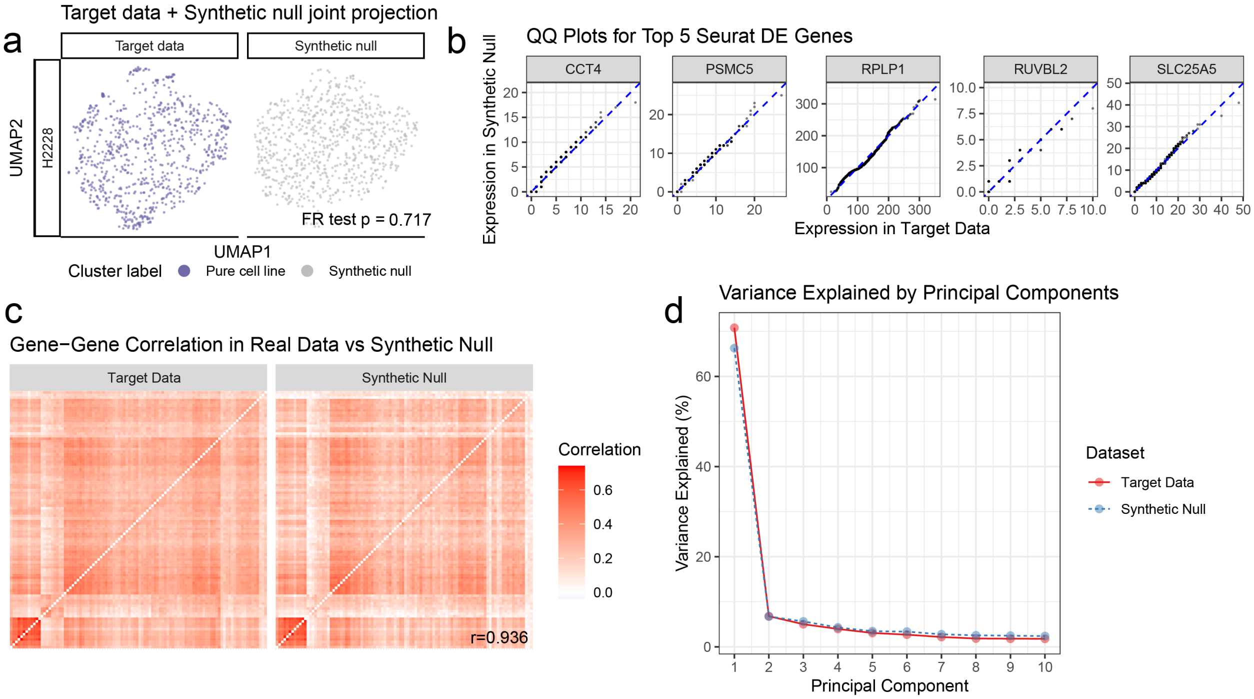


Fig. S16: **Similarity between the cell-line target dataset (Figure 3) and its corresponding synthetic null dataset. a**, UMAP embeddings of the target and synthetic null datasets; the Friedman–Rafsky (FR) test $P$ value assesses whether the two samples come from the same distribution. **b**, Quantile–quantile (QQ) plots for the top 5 Seurat DE genes (selected from the target data), comparing each gene's empirical quantiles in the target vs. synthetic null; points near the diagonal indicate similar distributions. **c**, Comparison of gene–gene rank-correlation (Kendall's $\tau$) matrices for the top 100 Seurat DE genes (selected from the target data) in the target vs. synthetic null; similarity is summarized by the Pearson correlation coefficient $r$ between the two matrices. **d**, Variance explained by the top 10 principal components (PCs), where PCs are computed on log(count+1)–transformed data for the target and synthetic null datasets, respectively.

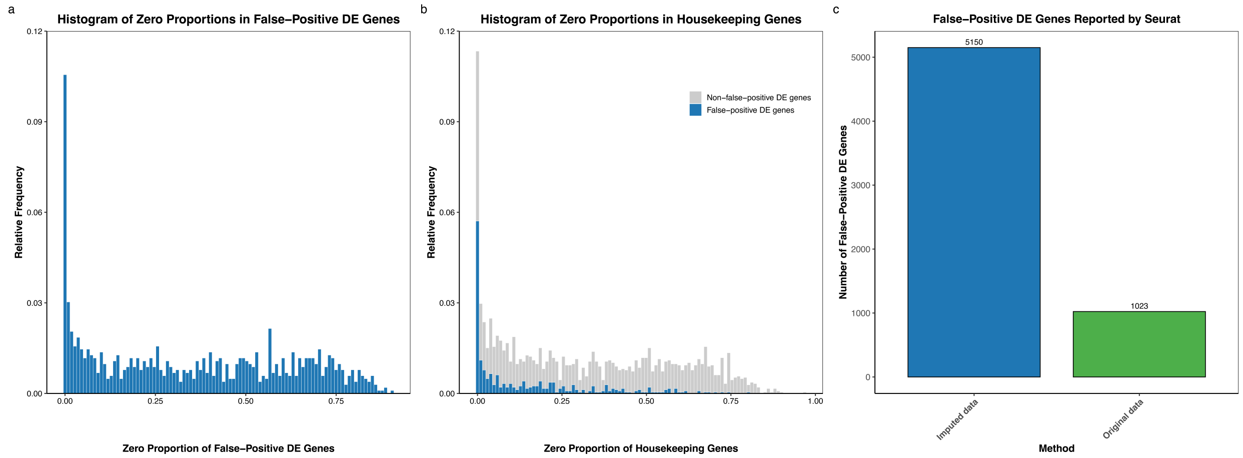


Fig. S17: **Relationship between gene dropout rate and false positive DE detection in the H2228 cell-line dataset** (which contains only a single cell type). **a**, Histogram of zero proportions among the 1,023 genes falsely identified as DE by Seurat. **b**, Histogram of zero proportions among the 310 housekeeping genes also identified as false positives (blue; gray background histogram is repeated from panel a). **c**, Number of DE genes reported by Seurat before (green) and after (blue) imputation using scImpute. Imputation increases false positives rather than resolving the double-dipping issue.

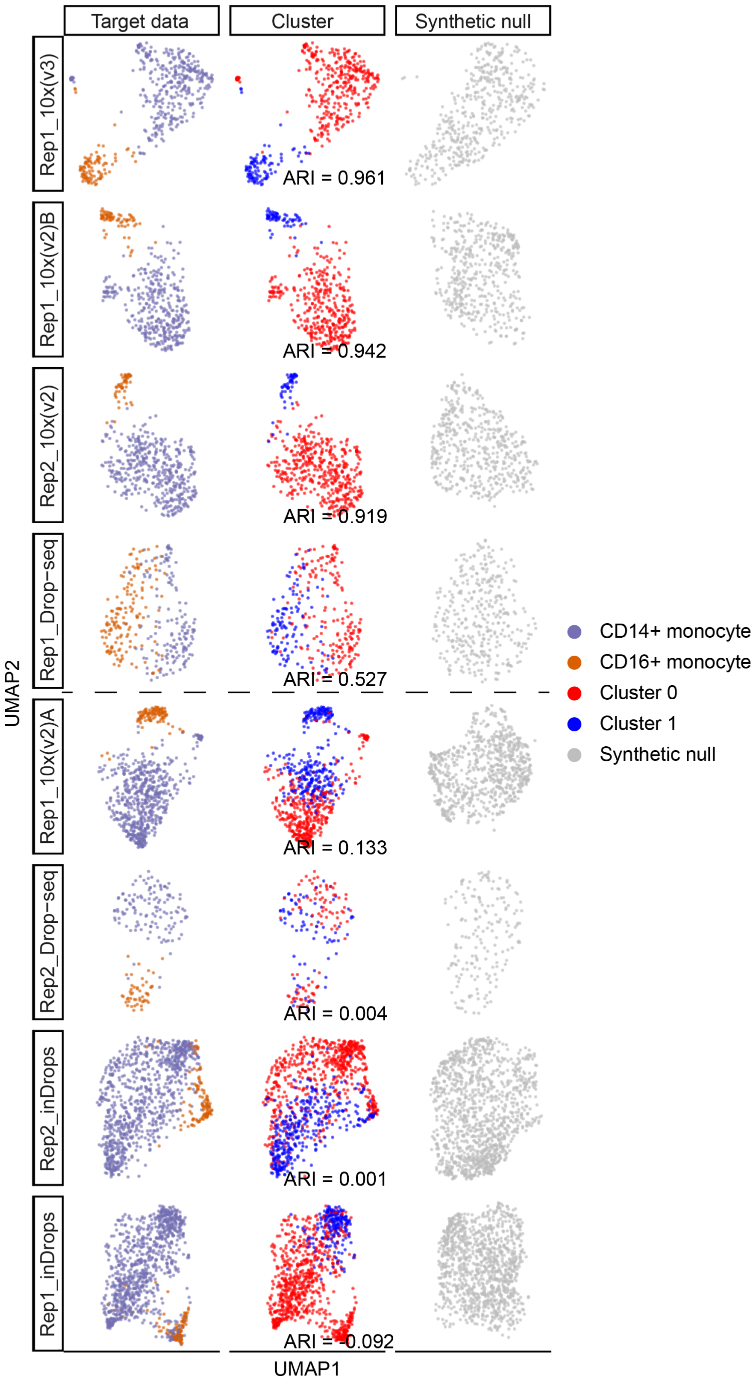


Fig. S18: **UMAP visualizations and Seurat clustering accuracies (ARIs) for the eight PBMC monocyte datasets (ordered by ARIs from high to low).** The first and second columns show the UMAP visualizations of the eight datasets (the target data), with cells labeled by monocyte subtypes (the first column) or Seurat clusters (the second column). The third column shows the UMAP visualizations of the synthetic null data corresponding to the eight target datasets. The horizontal dashed line separates the eight datasets into two groups (rows 1–4 and rows 5–8) based on clustering accuracy. Monocyte-subtype markers are expected to be more likely identified as post-clustering DE genes in the top four datasets (higher ARIs) compared to the bottom four datasets (lower ARIs).

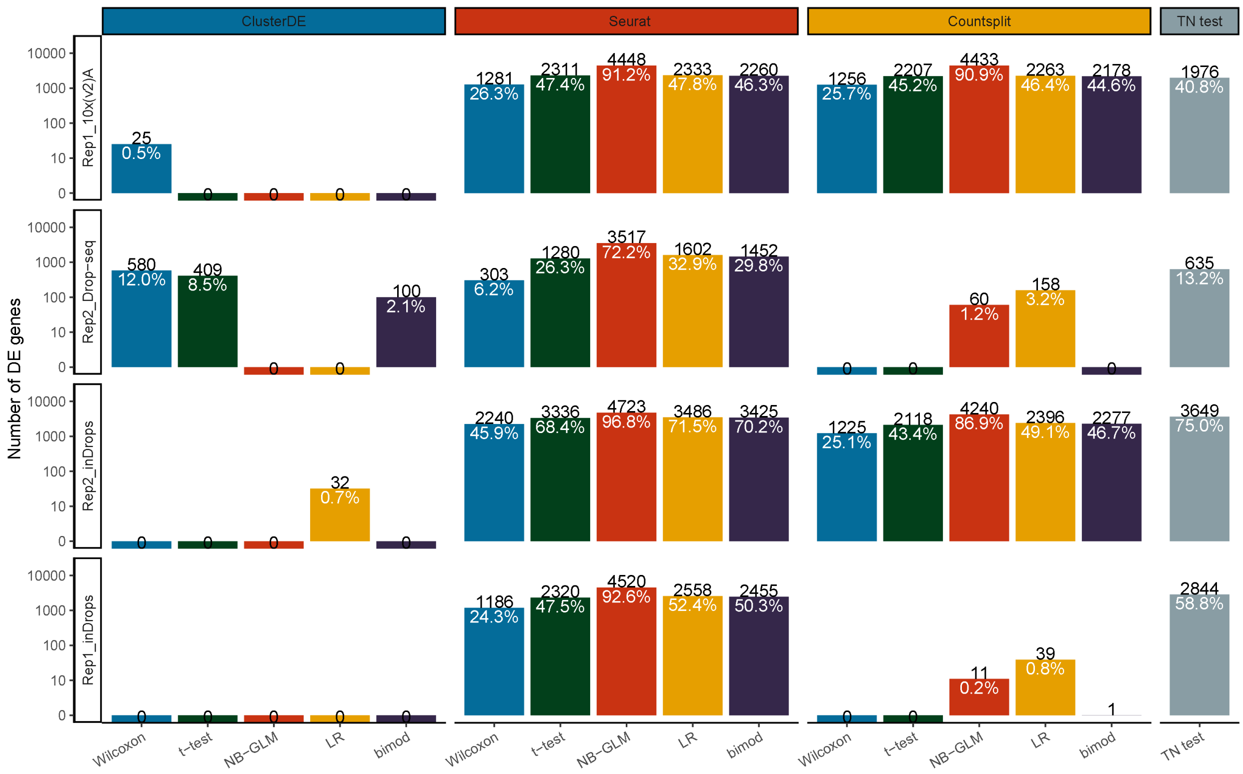


Fig. S19: **ClusterDE avoids false discoveries under double dipping.** Although the datasets contain two monocyte subtypes, the clustering results poorly align with the subtype labels (the bottom four datasets in Fig. S18), and therefore, few or no DE genes should be identified. The numbers in black indicate the numbers of DE genes identified, while the numbers in white represent the proportions of DE genes among all genes. As expected, in most cases, ClusterDE does not identify any DE genes.

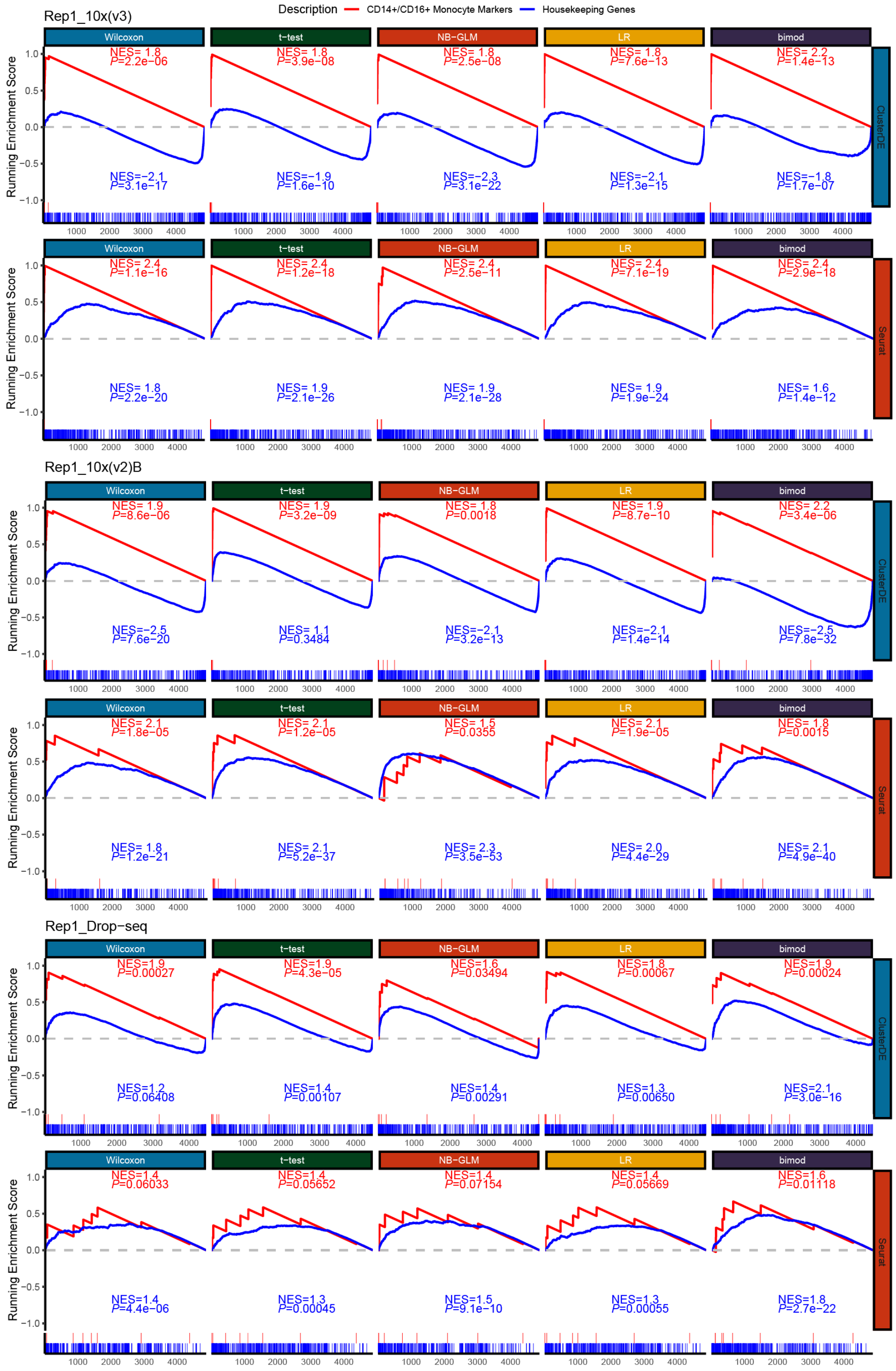


Fig. S20: **Gene set enrichment analysis (GSEA) of the ranked DE gene lists identified by ClusterDE and Seurat using five DE tests across three PBMC monocyte datasets.** The red lines represent the enrichment of the "CD14$^+$/CD16$^+$ Monocyte Marker Genes" set, while the blue lines represent the enrichment of the "Housekeeping Genes" set. The short vertical lines at the bottom indicate the rank distributions of genes from the two gene sets within each ranked DE gene list. The normalized enrichment score (NES) quantifies the direction and magnitude of enrichment, and the $P$ value reflects the statistical significance of the enrichment.

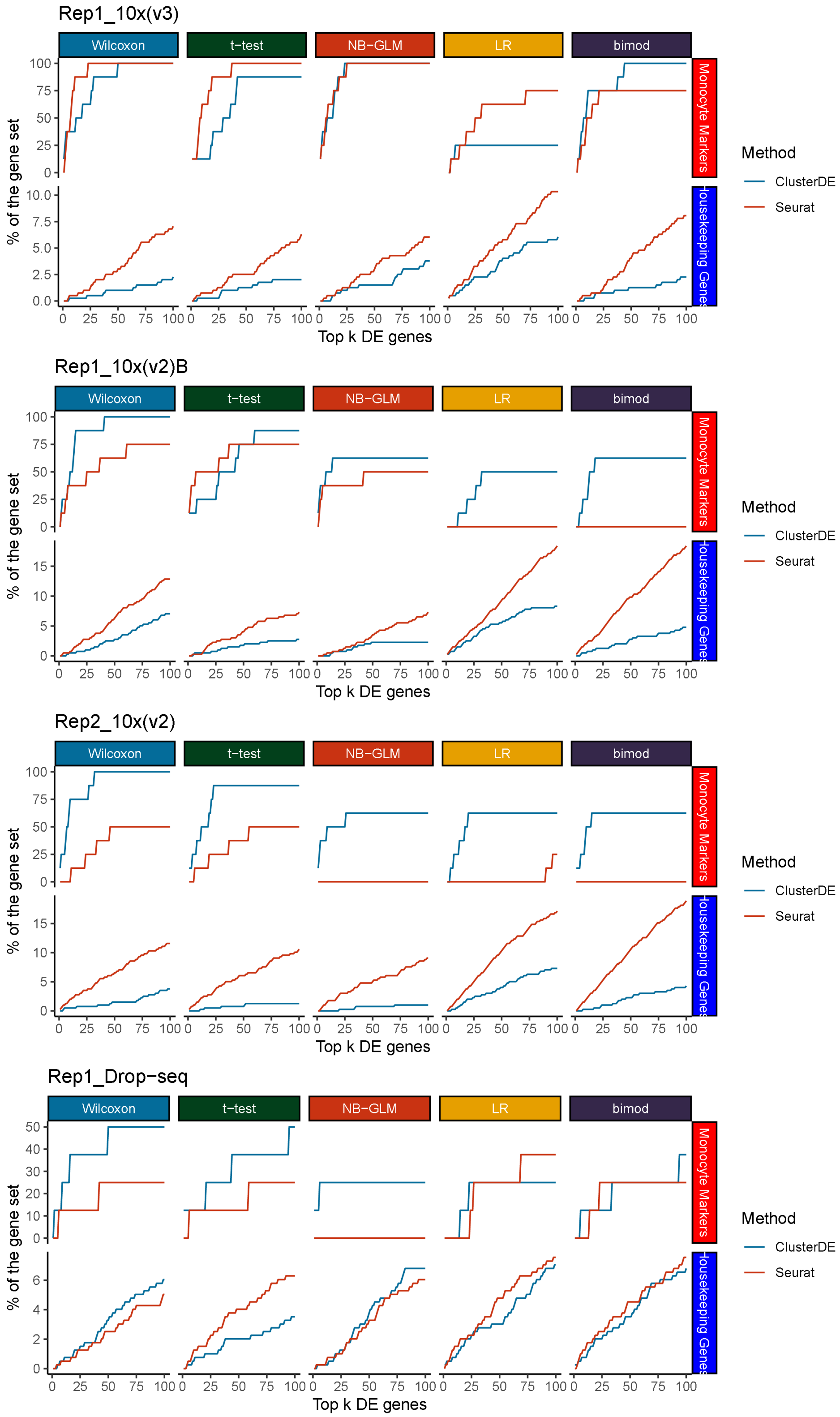


Fig. S21: **Overlaps between monocyte markers (or housekeeping genes) and the top $k$ DE genes, with $k$ ranging from 1 to 100.** The four panels correspond to four PBMC monocyte datasets. The horizontal axis represents $k$ (e.g., $k = 100$ corresponds to the top 100 DE genes from the DE gene lists), while the vertical axis indicates the proportion of detected monocyte subtype markers (top row in each panel) or housekeeping genes (bottom row in each panel) among the top $k$ DE genes identified by each of the five DE tests (columns). In most cases, ClusterDE (blue line) identifies a higher proportion of monocyte subtype markers and a lower proportion of housekeeping genes compared to Seurat (red line).

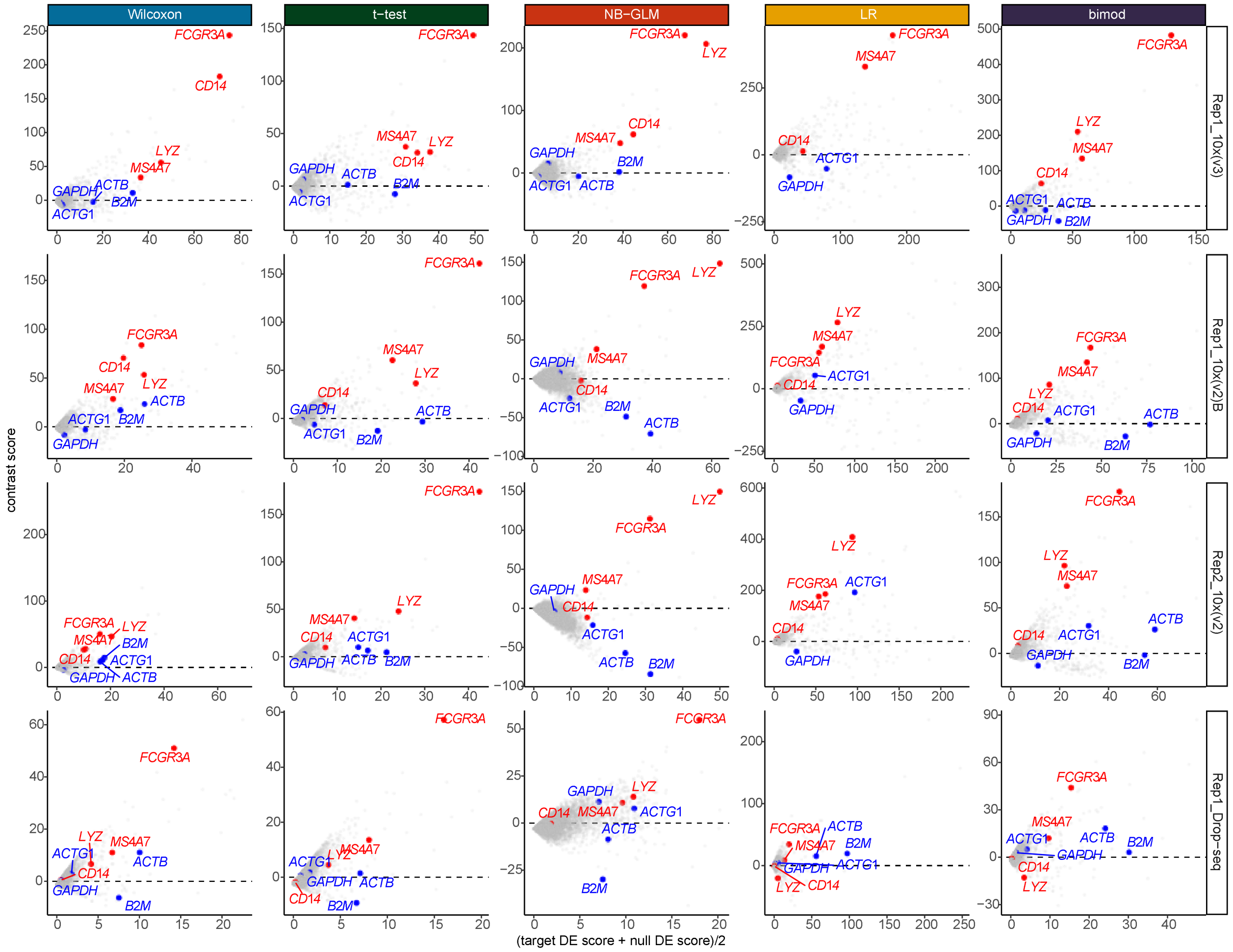


Fig. S22: **Minus-average (MA) plots of ClusterDE contrast scores (target DE score minus null DE score) vs. the averages of target DE scores and null DE scores.** Red points represent four well-known CD14$^+$/CD16$^+$ subtype markers, and blue points represent four well-known housekeeping genes. The dashed black line indicates contrast scores of 0. For housekeeping genes, their DE scores are large in both the target data and the synthetic null data, resulting in contrast scores centered around 0. As a result, these housekeeping genes would be ranked highly by Seurat (which only considers target DE scores) but not by ClusterDE. On the other hand, monocyte subtype markers have much larger DE scores in the target data than in the synthetic null data, leading to high contrast scores and top rankings by ClusterDE.

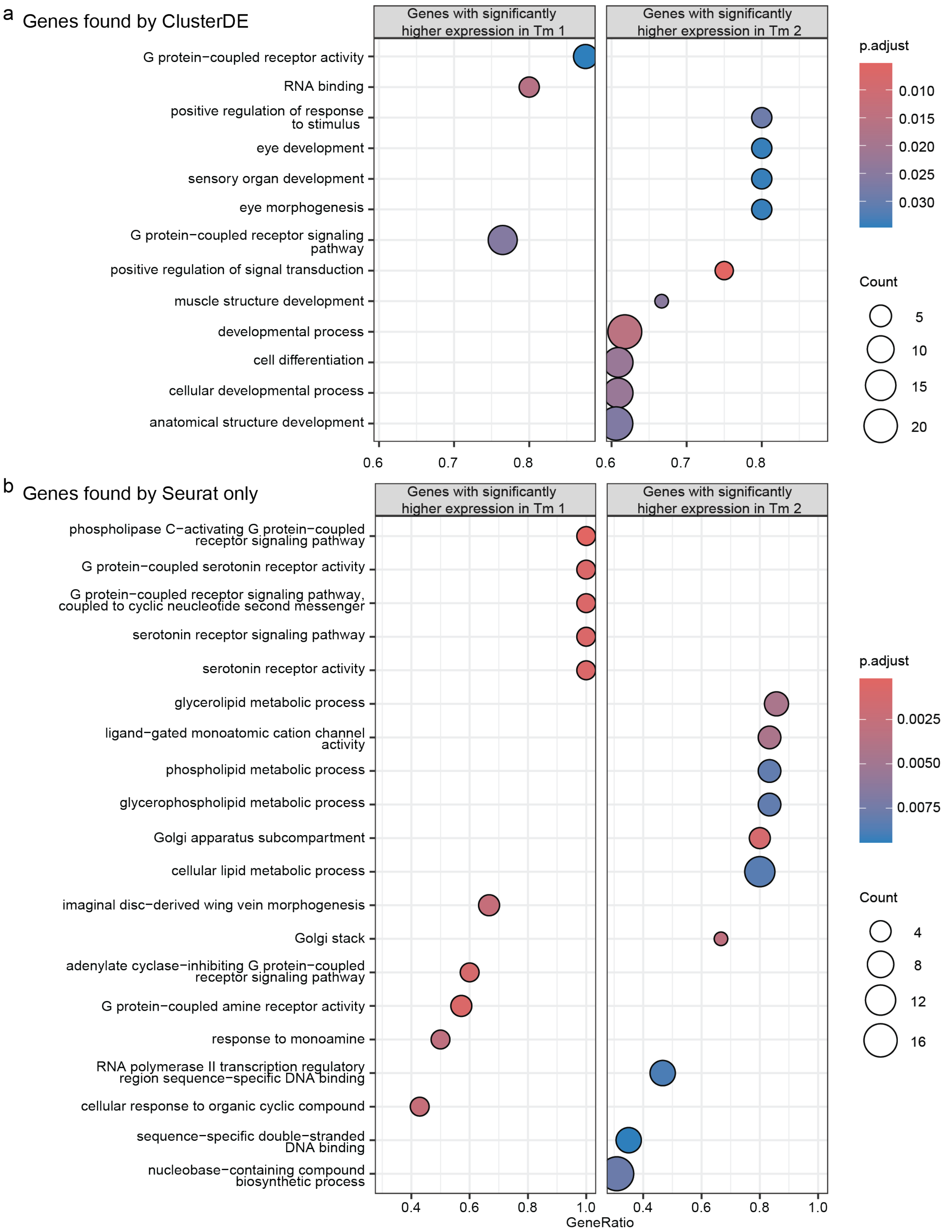


Fig. S23: **GSEA analysis for post-clustering DE genes between Tm1 and Tm2 cell types, identified by cell clustering, in the *Drosophila* neuronal visual system scRNA-seq dataset. a**, Results for DE genes found by ClusterDE. **b**, Results for DE genes found by Seurat using the Wilcoxon rank-sum test (not identified by ClusterDE).

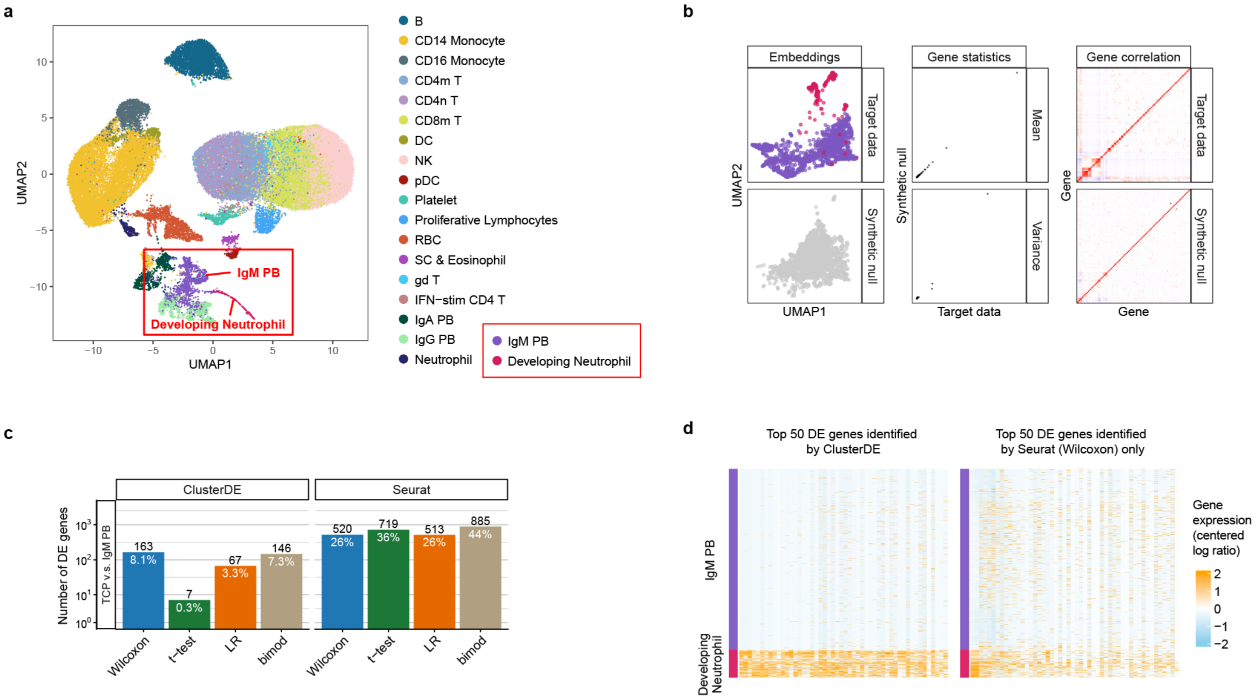


Fig. S24: **ClusterDE confirms the discrete nature of plasmablasts and developing neutrophils, refuting a continuous transdifferentiation trajectory found due to computational artifact. a**, UMAP visualization of the severe COVID-19 peripheral immune atlas based on annotations from [32], highlighting the purported developmental continuum bridging IgM plasmablasts (IgM PBs) and developing neutrophils. **b**, For the data subset containing the annotated IgM PBs and developing neutrophils, synthetic null data generated by ClusterDE closely resembled the target data in per-gene mean and variance (middle) and gene–gene rank correlations (right) while filling the gap between the two cell populations (left). **c**, Number and proportion of DE genes (at target FDR 0.05) between the IgM PBs and developing neutrophils identified by ClusterDE and Seurat (with DE tests Wilcoxon, t-test, LR, and bimod). The substantial number of DE genes confirmed by ClusterDE indicates these are discrete cell types rather than an artificially bisected continuum. (d) Heatmap displaying the expression profiles of the top 50 DE genes identified by ClusterDE versus those identified by Seurat (with Wilcoxon as the DE test) only.

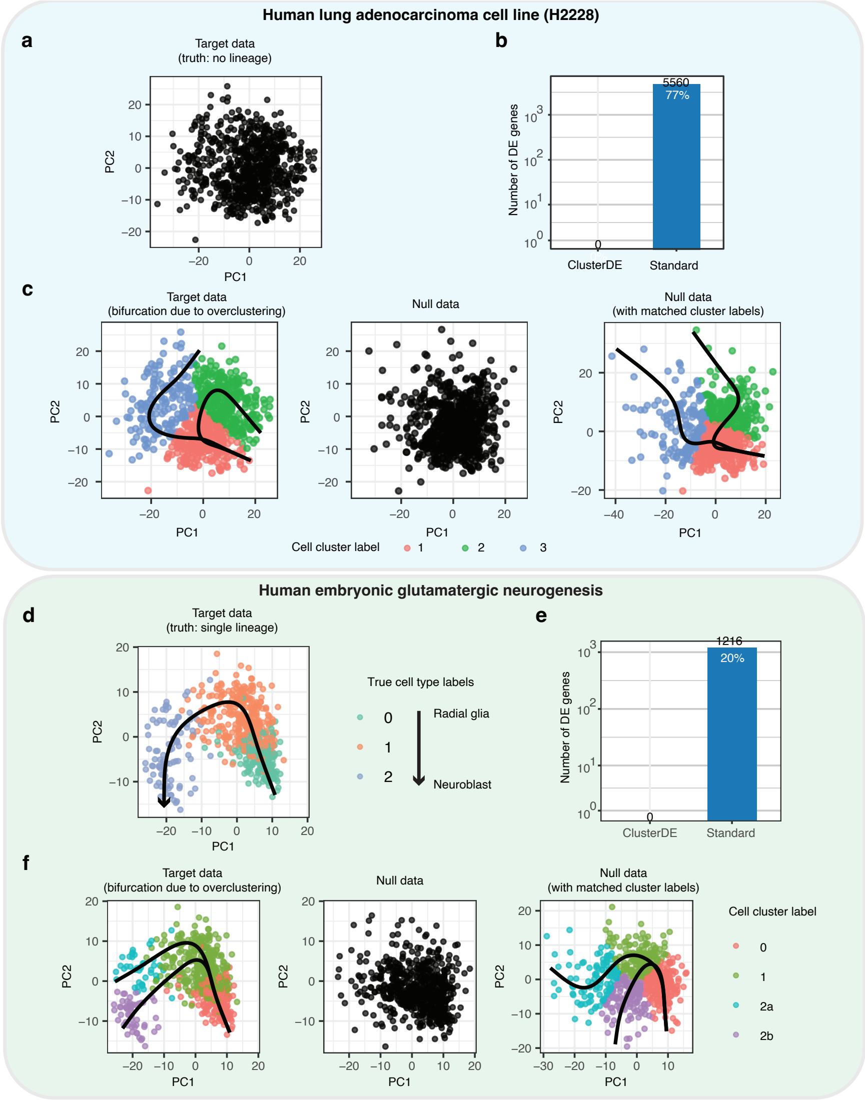


Fig. S25: **ClusterDE applied to two datasets lacking bifurcating trajectories.** (1) A human lung adenocarcinoma cell line dataset (H2228) [24] (**a**–**c**), which contains no lineage structure. (2) A human embryonic glutamatergic neurogenesis dataset [33] (**d**–**f**), in which cells follow a continuous, single-lineage trajectory from radial glia to neuroblast cells. **a**, H2228 dataset with no underlying lineage structure. **b**, ClusterDE identifies 0 DE genes, while the standard approach identifies 5,560 DE genes. **c**, Target data (left; H2228 dataset from panel a) and corresponding synthetic null data (middle), visualized using the top two principal components (PCs) of the target data to enable direct comparison of embeddings. Target cells are labeled with K-means clusters (mimicking over-clustering in the standard approach) and Slingshot-inferred bifurcating trajectories. Synthetic null data (right), with matched cluster and trajectory labels, is shown in its own PC space. **d**, Neurogenesis dataset with true cell type labels from [33], showing a continuous, single-lineage progression from radial glia to neuroblasts. **e**, ClusterDE identifies 0 DE genes, while the standard approach identifies 1,216 DE genes. **f**, Target data (left; same dataset as in panel d) and synthetic null data (middle), visualized using the top two PCs of the target data to enable direct comparison of embeddings. Target data is labeled with K-means clusters and Slingshot-inferred bifurcation trajectories. Synthetic null data (right), with matched cluster and trajectory labels, is shown in its own PC space.

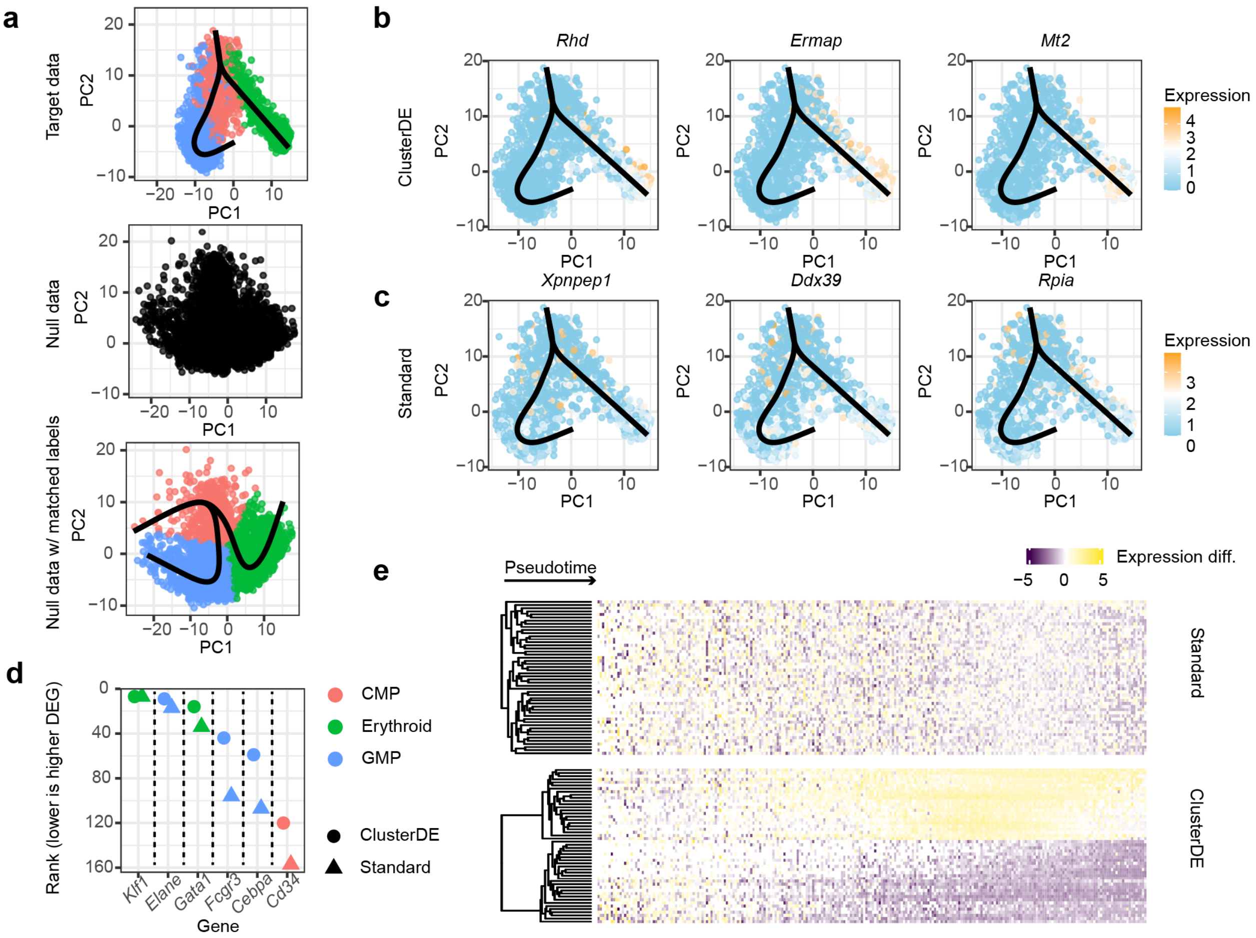


Fig. S26: **ClusterDE identifies lineage-specific genes in a true bifurcating hematopoietic trajectory. a**, Analysis of mouse bone marrow common myeloid progenitors (CMPs), which bifurcate into granulocyte–monocyte progenitors (GMPs) and erythroid cells [34]. The target data with annotated cell types and Slingshot-inferred trajectories are shown at the top. The synthetic null data, representing a single-lineage null model, are shown in the middle, and the null data after matching clusters to the target labels and re-inferring trajectories are shown at the bottom. Embeddings are displayed using the first two principal components of the target data. **b**, Expression patterns of the top three DE genes identified by ClusterDE, shown in principal-component space with the inferred trajectories overlaid. **c**, Expression patterns of the top three DE genes identified uniquely by the standard approach, which applies the Wilcoxon rank-sum test without synthetic null calibration. **d**, Rankings of known lineage markers (*Klf1*, *Elane*, *Gata1*, *Fcgr3*, *Cebpa* and *Cd34*) in the DE-gene lists from ClusterDE and the standard approach. Lower ranks indicate higher placement in the corresponding DE-gene list; circles and triangles denote ClusterDE and the standard approach, respectively. **e**, Heatmaps of expression differences between the two terminal lineages across binned pseudotime for genes identified by the standard approach (top) and ClusterDE (bottom). Columns are ordered by average pseudotime, and genes are ordered by hierarchical clustering.

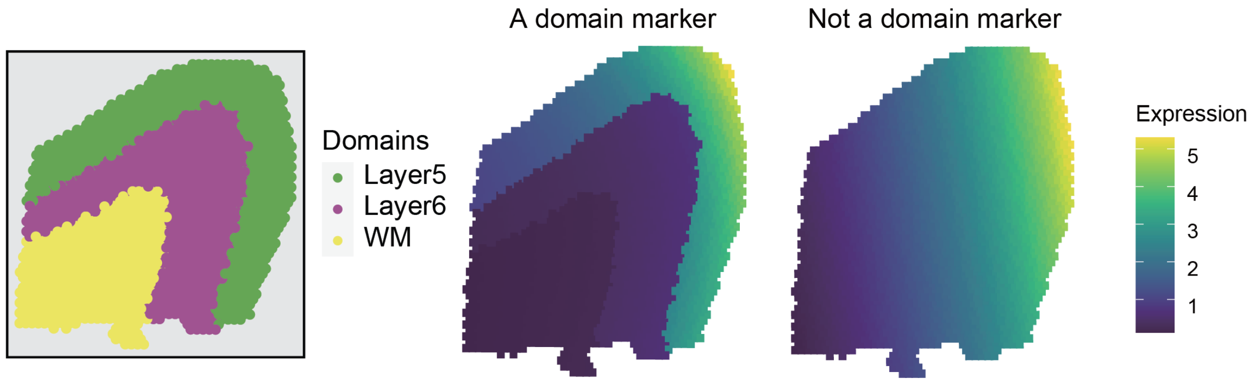


Fig. S27: **Illustration of the definition for spatial-domain marker genes under the ClusterDE framework.** The left panel depicts three domains. The right panel presents examples: a domain marker gene showing clear discontinuity at the domain boundary and a non-marker gene exhibiting a continuous transition across two domains.

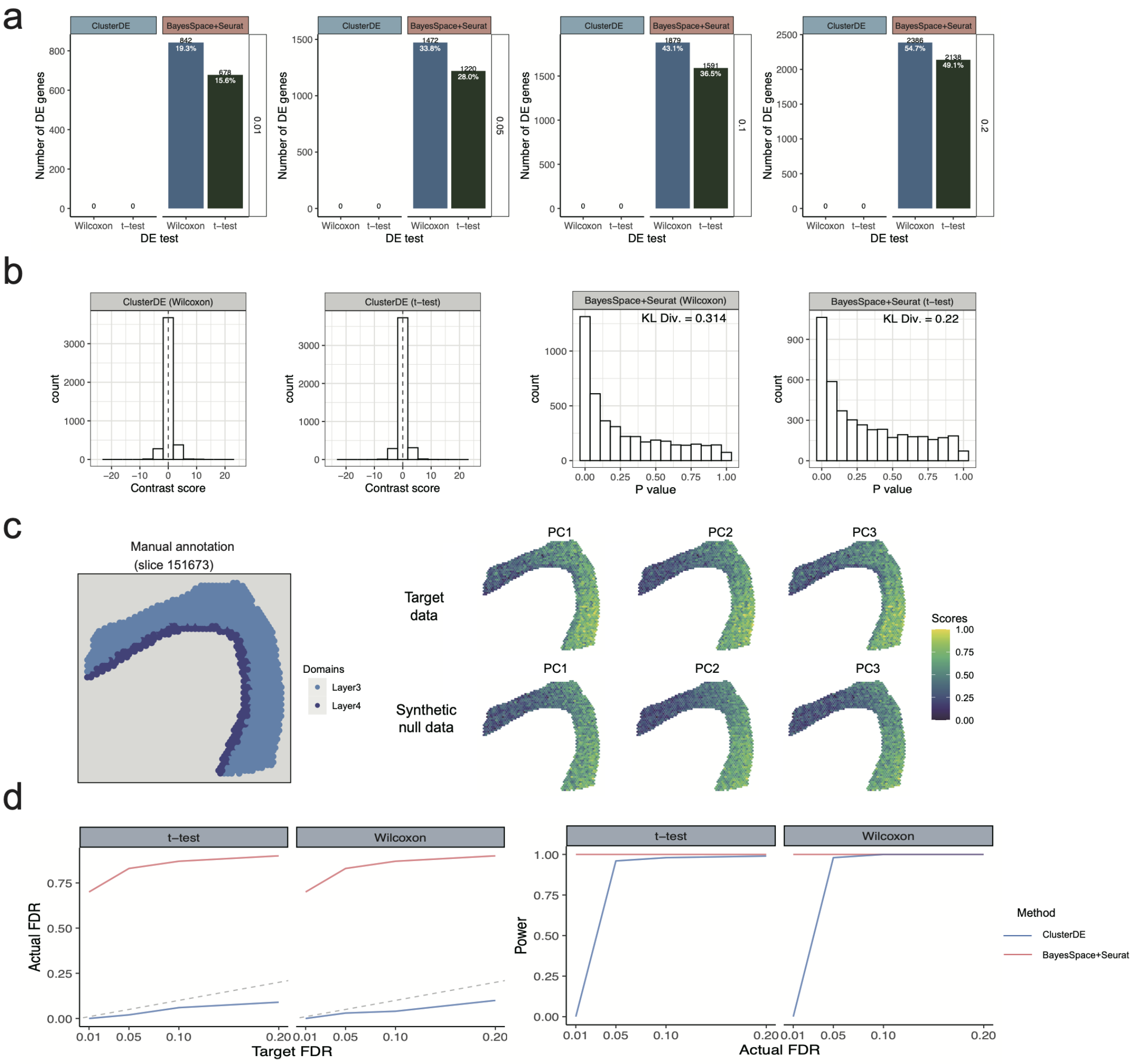


Fig. S28: **ClusterDE achieves reliable FDR control and good statistical power in identifying spatial-domain marker genes from simulated SRT datasets containing one or two spatial domains.** **a**, Numbers of DE genes identified by ClusterDE and BayesSpace+Seurat (at the target FDRs of 0.01, 0.05, 0.1, and 0.2) in the simulated data containing only one domain. The numbers in black indicate the numbers of DE genes, while the numbers in white represent their proportions among all genes. The statistical tests used in the DE step are the Wilcoxon rank-sum test (Wilcoxon) and t-test. **b**, Validity checks for the contrast scores of ClusterDE and the $P$ values of BayesSpace+Seurat on the simulated one-domain dataset. The first and second columns show that the ClusterDE contrast scores of all genes are approximately symmetric around 0, satisfying ClusterDE's assumption for FDR control. The third and fourth columns show histograms of $P$ values obtained from BayesSpace+Seurat. A larger empirical Kullback-Leibler divergence (KL div.) between the $P$ value distribution and the theoretical Uniform[0,1] distribution indicates a more severe violation of the $P$ value uniformity assumption. The results show that $P$ values from BayesSpace+Seurat exhibit severe deviations from the uniform distribution for all genes. **c**, Visualization of manual annotation of real SRT data and the spatial expression patterns of three representative principal components (PC1, PC2, and PC3) among the top five principal components in the target data and synthetic null data, respectively. **d**, The FDRs and power of ClusterDE and BayesSpace+Seurat using t-test and Wilcoxon at various target FDRs.

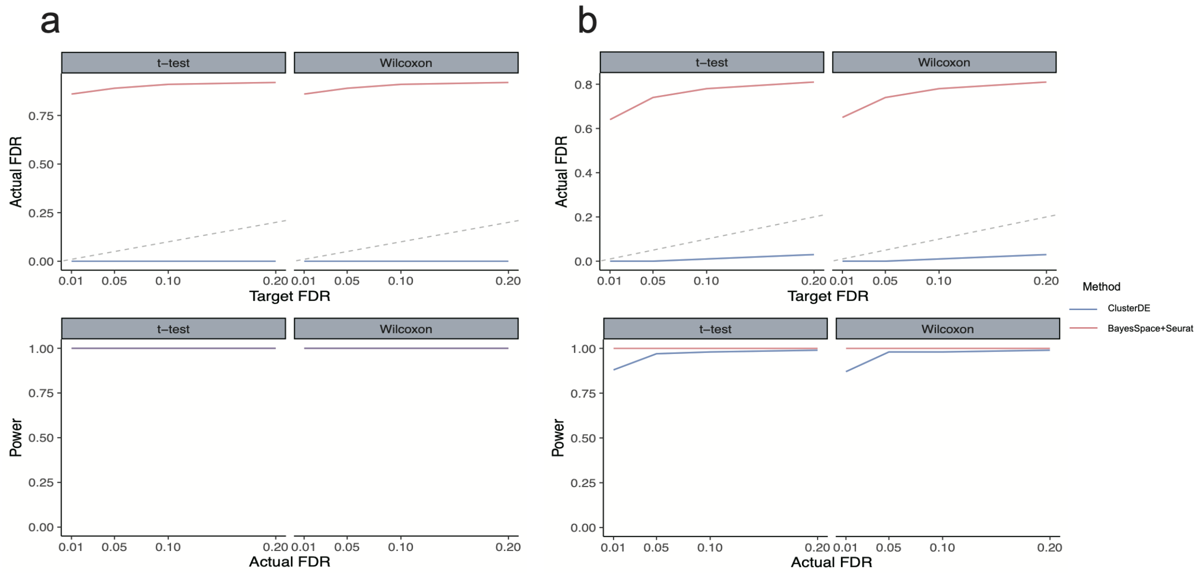


Fig. S29: **ClusterDE achieves reliable FDR control and good statistical power in identifying spatial-domain marker genes from a simulated SRT dataset containing three domains (Fig. 6a; see "Simulation setting with three spatial domains, including 200 and 168 true spatial-domain marker genes" in the Supplementary Methods). a**, The FDRs and power of ClusterDE and BayesSpace+Seurat for identifying marker genes between Layer5 and Layer6 at various target FDRs. **b**, The FDRs and power of ClusterDE and BayesSpace+Seurat for identifying marker genes between Layer6 and WM at various target FDRs.

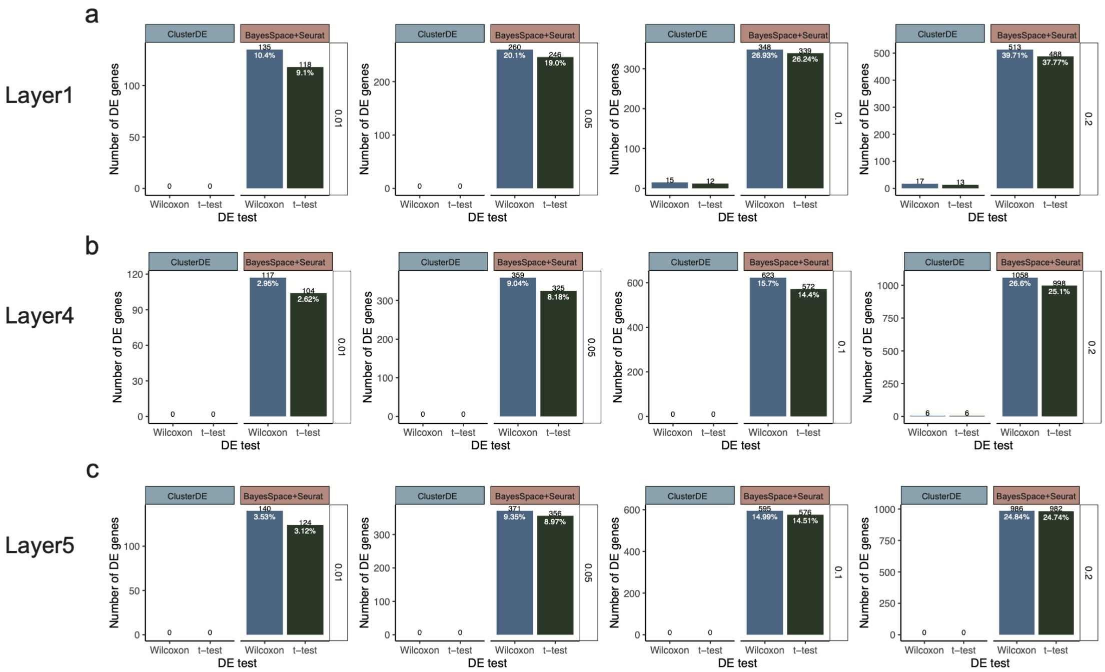


Fig. S30: **ClusterDE achieves reliable FDR control in identifying spatial-domain marker genes from three target SRT datasets, each derived from a single domain of the real DLPFC SRT dataset (Fig. S31).** The panels (rows) **a–c** represent the three target datasets: Layer1, Layer4, and Layer5. The numbers of DE genes identified by ClusterDE and BayesSpace+Seurat at target FDR levels of 0.01, 0.05, 0.1, and 0.2 are shown for each target dataset. Numbers in black indicate the numbers of identified DE genes, while numbers in white represent their proportions among all genes. The statistical tests used for the DE analysis are the Wilcoxon rank-sum test (Wilcoxon) and t-test.

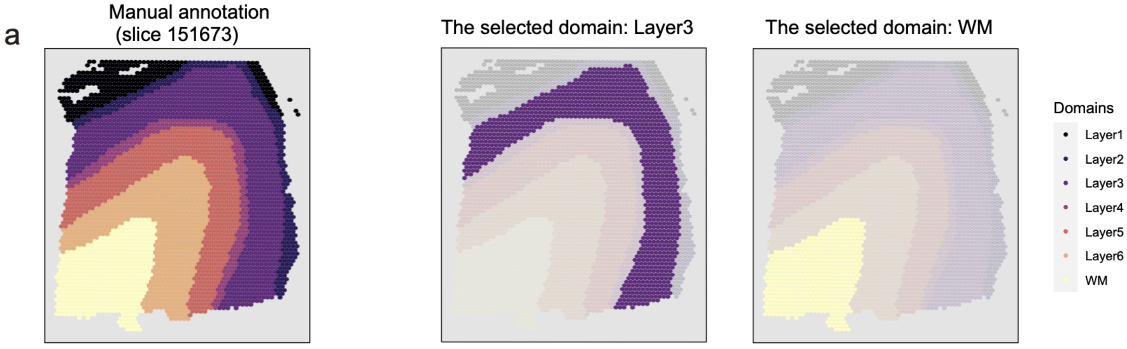


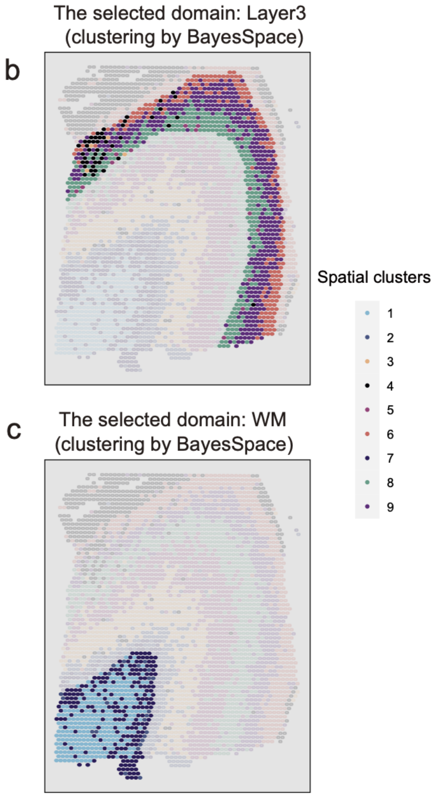


| Final identify | Within-layer clusters | Compared clusters | # of DE genes found by ClusterDE | # of DE genes found by BayesSpace+Seurat |
|---|---|---|---|---|
| Layer3 | 3,4,5,6,8,9 | 9 vs. 8 | 0 | 583 |
| | | (9 & 8) vs. 6 | 0 | 772 |
| | | (9 & 8 & 6) vs. 4 | 0 | 1609 |
| | | (9 & 8 & 6 & 4) vs. 5 | 0 | 303 |
| | | (9 & 8 & 6 & 4 & 5) vs.3 | 0 | 209 |

| Final identify | Within-layer clusters | Compared clusters | # of DE genes found by ClusterDE | # of DE genes found by BayesSpace+Seurat |
|---|---|---|---|---|
| WM | 1,2,7 | 1 vs. 7 | 0 | 915 |
| | | (1 & 7) vs. 2 | 0 | 531 |

Fig. S31: **Application of ClusterDE to the LIBD human dorsolateral prefrontal cortex (DLPFC) SRT dataset. a**, Visualization of the entire dataset with manual annotation and selected domains: Layer3 (purple) and WM (yellow). **b**, Left panel: Spatial clustering results for Layer3 using BayesSpace. Right panel: Numbers of DE genes identified at the target FDR of 0.05 by ClusterDE and BayesSpace+Seurat (using the Wilcoxon rank-sum test). **c**, Left panel: Spatial clustering results for WM using BayesSpace. Right panel: Numbers of DE genes identified at the target FDR of 0.05 by ClusterDE and BayesSpace+Seurat (using the Wilcoxon rank-sum test).

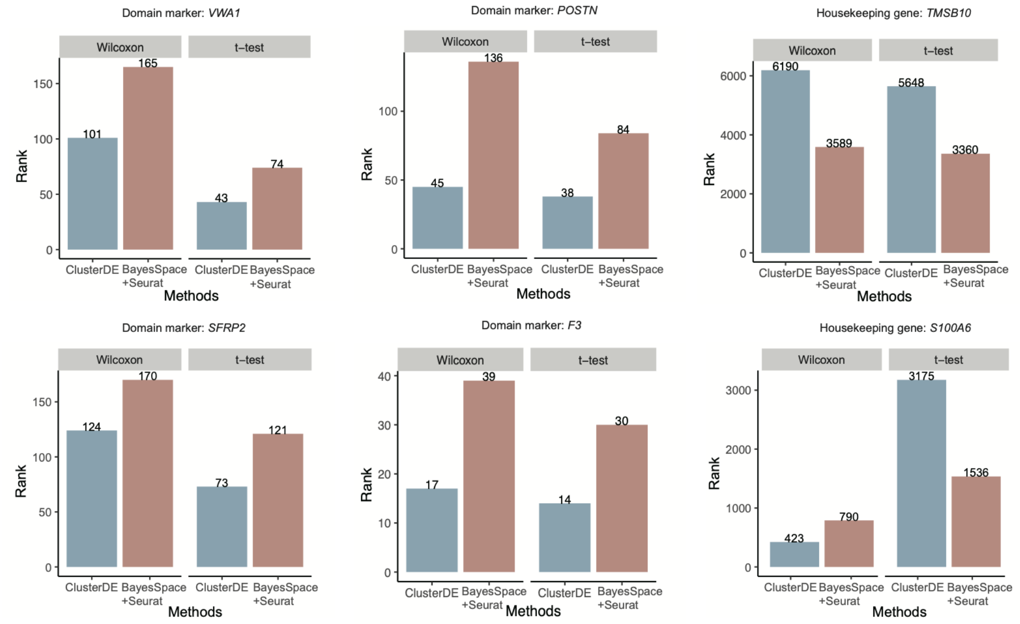


Fig. S32: **Application of ClusterDE to the PDAC-B cancer slice from the human primary pancreatic cancer (PDAC) SRT data.** Bar plots show the rankings of domain markers and housekeeping genes identified by ClusterDE and BayesSpace+Seurat. The top row shows the rankings of three exemplary genes between the Duct epithelium and Interstitium: the domain markers *VWA1* and *POSTN* and the housekeeping gene *TMSB10*. The bottom row shows the rankings of three exemplary genes between the Interstitium and Cancer region: the domain markers *SFRP2* and *F3* and the housekeeping gene *S100A6*.

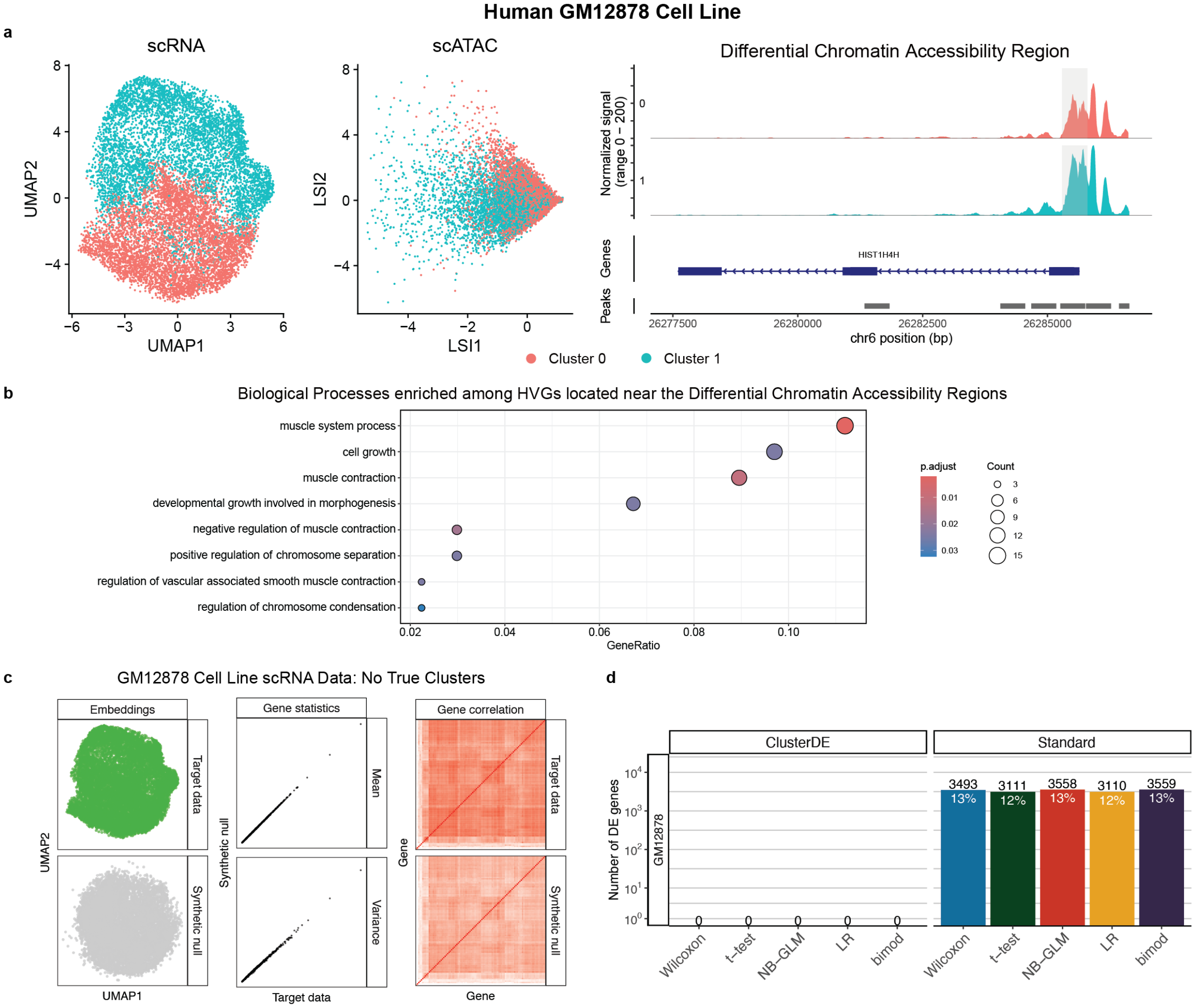


Fig. S33: **ClusterDE eliminates over-clustering artifacts and prevents false-positive discoveries in single-cell multiome data analysis.** **a**, UMAP visualization (left) of RNA-level gene expression and LSI visualization (middle) of ATAC-level chromatin accessibility for the GM12878 human lymphoblastoid cell line from SHARE-seq [39], with cells erroneously over-clustered into two groups based on RNA expression using Seurat's default VST workflow. The coverage plot (right) shows a differentially accessible (DA) region near the gene *HIST1H1C* identified by the standard Wilcoxon test using these RNA-based clusters—despite the data originating from a single homogeneous cell type. **b**, Gene Ontology (GO) enrichment analysis of 179 highly variable genes located nearest to the DA regions identified in (a) reveals strong enrichment for muscle-related biological processes, which are biologically implausible for lymphoblastoid cells. **c**, ClusterDE generates a synthetic null dataset that closely resembles the RNA-seq modality in UMAP (left), per-gene mean and variance (middle), and gene–gene correlation structure (right). **d**, ClusterDE consistently identifies zero DE genes between the two RNA-based clusters, correctly indicating that the clusters should be merged and that downstream comparisons—such as DA testing on the ATAC modality—should not be performed. In contrast, standard DE tests report thousands of significant genes, falsely supporting the spurious clustering and leading to propagation of over-clustering artifacts across modalities.

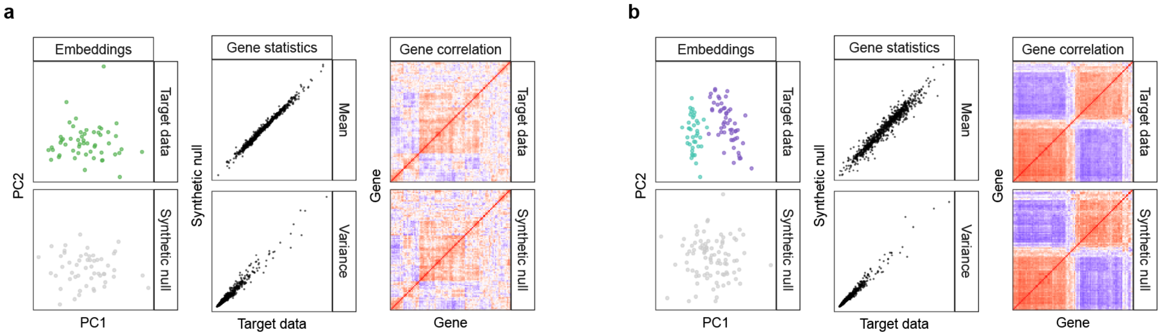


Fig. S34: **Sanity check of ClusterDE when applied to the bulk microarray data of glioblastoma (GBM) subtypes.** **a**, When the target data contained samples from a single subtype (negative case), the synthetic null data generated by ClusterDE closely resembled the target data in terms of PCA structure (left), per-gene mean and variance (middle), and gene–gene rank correlations (right). **b**, When the target data contained samples from two distinct subtypes (positive case), the synthetic null data generated by ClusterDE filled the gap between subtypes while preserving key statistical properties.

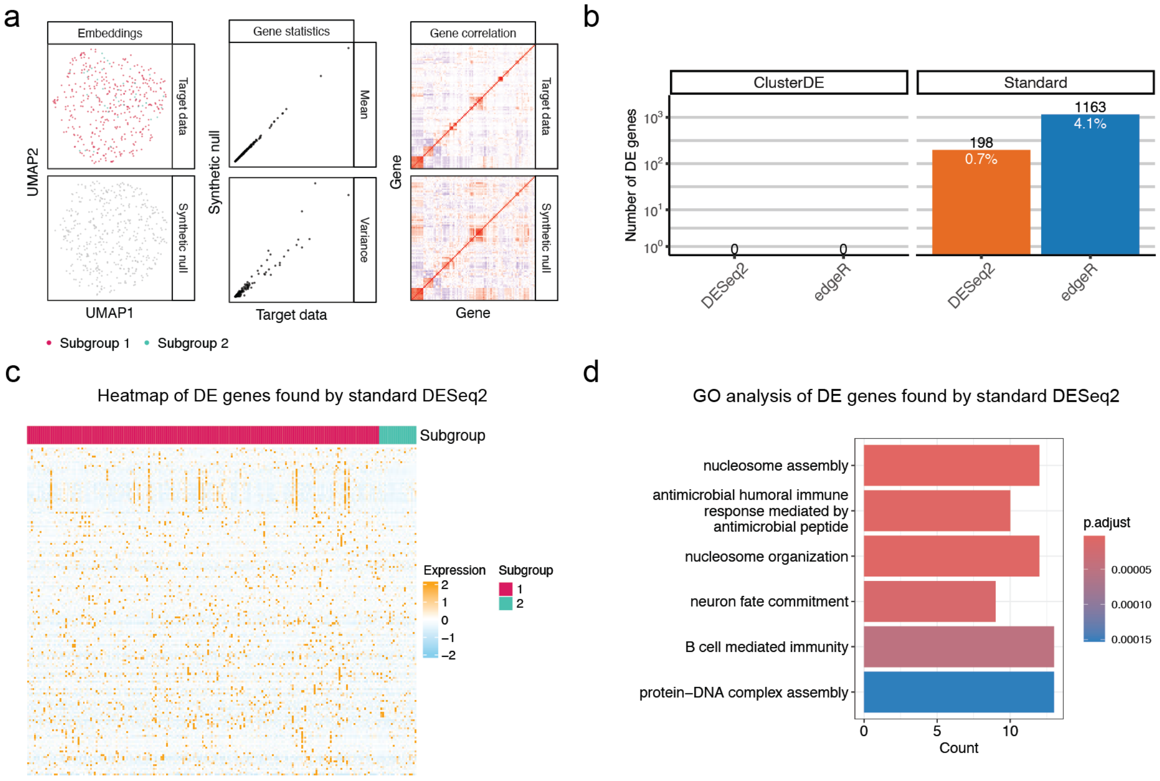


Fig. S35: **ClusterDE mitigates over-clustering bias in cluster marker identification for population-scale RNA-seq data.** **a**, UMAP visualization of the target TCGA ovarian cancer patient data and its corresponding synthetic null data (left), per-gene expression mean and variance (middle), and gene–gene rank correlations (right), showing that the synthetic null closely resembles the target data. **b**, Numbers of DE genes (at target FDR 0.05) identified by ClusterDE and by standard post-clustering DE analysis using DESeq2 and edgeR. Numbers in black and white indicate the counts and proportions of DE genes, respectively. **c**, Expression heatmap of DE genes identified by post-clustering DESeq2 analysis for the two patient subgroups. **d**, Gene Ontology (GO) enrichment analysis of the DE genes identified by post-clustering DESeq2 reveals strong enrichment for cell cycle processes and a developmental pathway, rather than to ovarian cancer–specific processes.

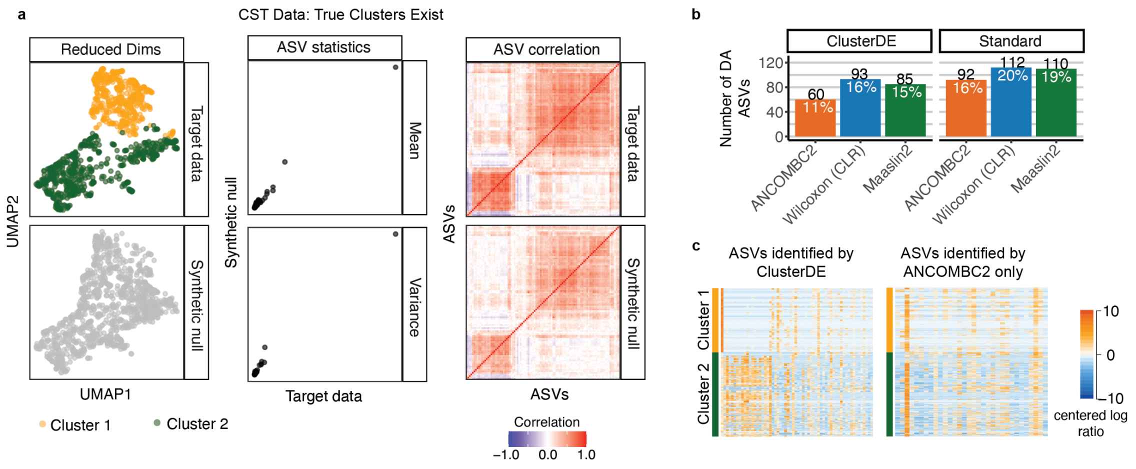


Fig. S36: **ClusterDE controls false positives and improves specificity in post-clustering differential abundance (DA) analysis of the 16S rRNA sequencing vaginal microbiome data.** **a**, For the vaginal microbiome dataset with two community state types, synthetic null data also maintained similarity to the target data in distributional and correlation structure, while removing subgroup structure. **b**, Bar plot comparing the number and percentage of DA ASVs found by standard methods versus ClusterDE. **c**, Heatmap showing CLR-normalized data for DA ASVs identified by ClusterDE (left), a subset of those found by ANCOMBC2, and those found by ANCOMBC2 only (right). Taxa order in all heatmaps is determined by default hierarchical clustering from `heatmap.2()`.

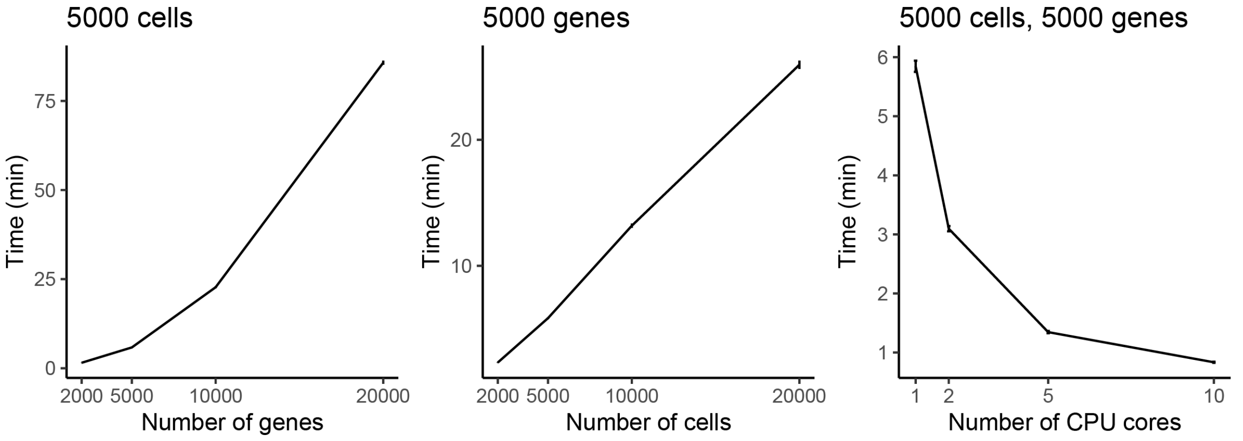


Fig. S37: **Time complexity of synthetic null data generation.** The computational time scales approximately quadratically with the number of genes, linearly with the number of cells, and is inversely proportional to the number of CPU cores under parallel computing.

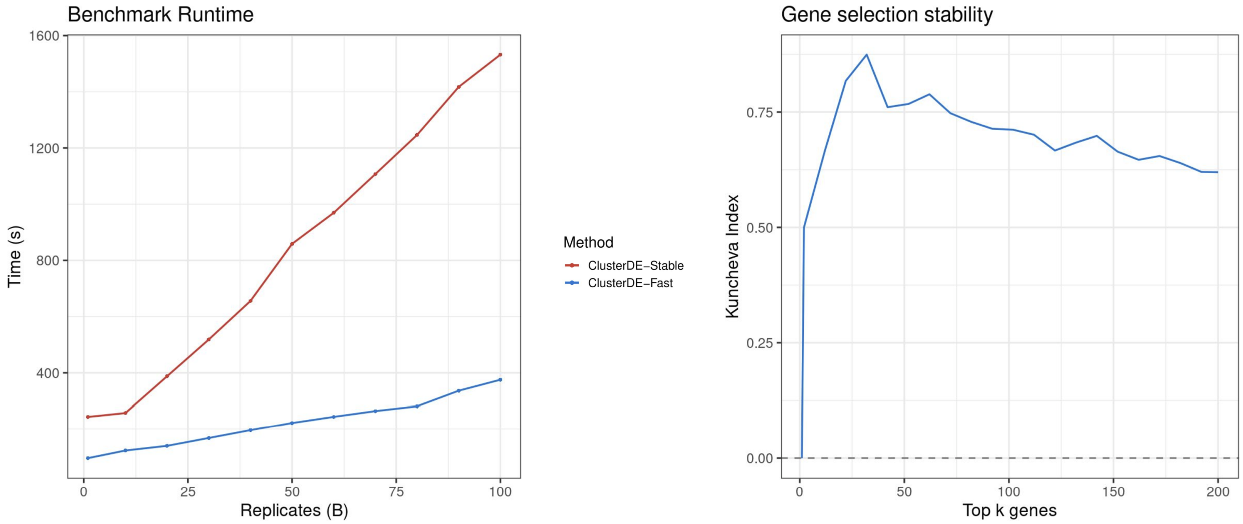


Fig. S38: **Performance of ClusterDE-Fast versus ClusterDE-Stable on PBMC CD4$^+$ and cytotoxic T cells.** Left: Runtime comparison between ClusterDE-Stable and ClusterDE-Fast across $B$ synthetic null datasets, showing total time required to generate synthetic null data, perform DE analysis, and obtain final DE genes. Right: Comparison of top $k$ gene selection results between ClusterDE-Stable and ClusterDE-Fast using the Kuncheva index across varying values of $k$, where higher values indicate stronger agreement. In high-dimensional omics settings, Kuncheva index values above 0.5 are generally considered evidence of strong selection stability; here, the observed indices are mostly greater than 0.5, indicating high concordance between ClusterDE-Fast and ClusterDE-Stable.

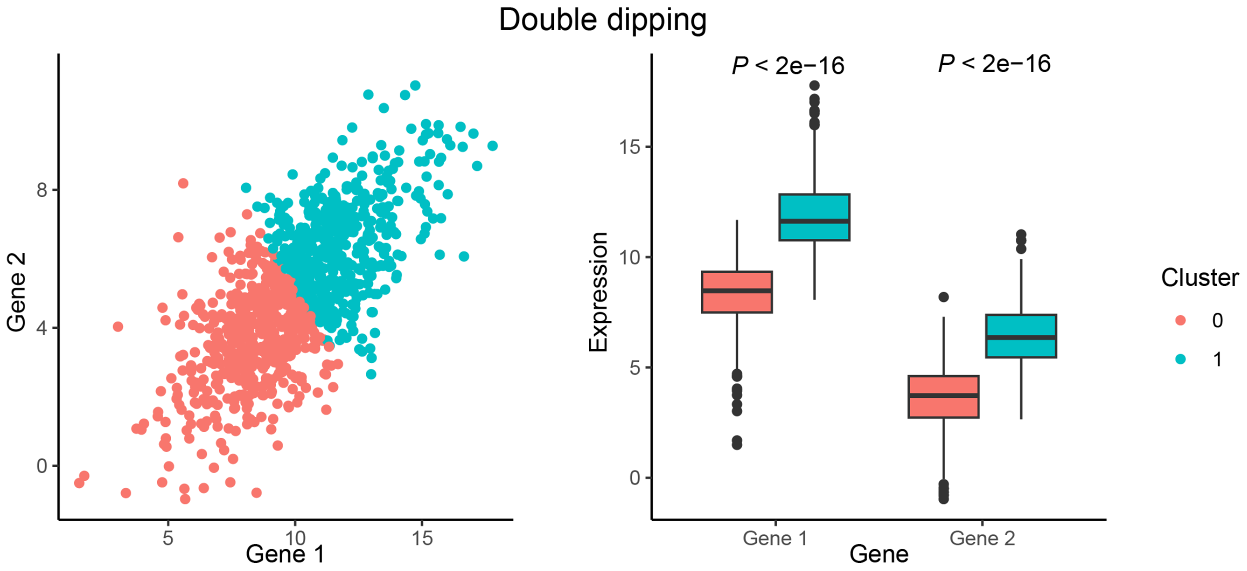


Fig. S39: A toy example illustrating the double-dipping issue. The expression levels of two genes follow a bivariate Gaussian distribution, representing a homogeneous cell type. However, when K-means clustering is applied to divide the cells into two clusters, the two genes are artificially forced to exhibit different distributions between the clusters. As a result, both genes are incorrectly identified as post-clustering DE genes, leading to the false conclusion that the two clusters represent distinct cell types defined by these genes.

#### S4 Supplementary Tables

| Replicates ($B$) | 2 | 4 | 6 | 8 | 10 | 12 | 14 | 16 | 18 | 20 |
|---|---|---|---|---|---|---|---|---|---|---|
| Correlation | 0.269 | 0.862 | 0.871 | 0.876 | 0.889 | 0.928 | 0.949 | **0.962** | **0.965** | **0.970** |

Table S1: **Correlation of gene-level selection frequencies from two independent runs of ClusterDE-Stable.** We ran ClusterDE-Stable on two disjoint sets of $B$ synthetic null replicates and, across genes, computed the correlation between the resulting selection-frequency vectors. The correlation exceeds 0.95 for $B > 14$.

#### S5 Supplementary Algorithms

Algorithm S1: **ClusterDE-Core workflow for scRNA-seq data**

**Input:** Cell-by-gene UMI count matrix $\mathbf{Y} = [Y_{ij}] \in \mathbb{N}_{\geq 0}^{n\times m}$; DE test $\mathcal{T}_{\mathrm{DE}}$; target FDR level $q$.

1. **Generate synthetic null data.**
For each gene $j = 1, \ldots, m$, estimate
$$(\widehat{\mu}_j, \widehat{\sigma}_j) \leftarrow \arg\max_{\mu_j,\sigma_j} \sum_{i=1}^{n} \log f_{\mathrm{NB}}(Y_{ij}; \mu_j, \sigma_j), \qquad \widehat{F}_j \leftarrow F_{\mathrm{NB}}(\,\cdot\,; \widehat{\mu}_j, \widehat{\sigma}_j),$$
where $f_{\mathrm{NB}}$ and $F_{\mathrm{NB}}$ denote the NB probability mass function and cumulative distribution function, respectively.
For $i = 1, \ldots, n$ and $j = 1, \ldots, m$, compute
$$V_{ij} \overset{\text{i.i.d.}}{\sim} \mathrm{Uniform}(0,1), \quad U_{ij} \leftarrow V_{ij}\widehat{F}_j(Y_{ij}) + (1 - V_{ij})\widehat{F}_j(Y_{ij} + 1), \quad G_{ij} \leftarrow \Phi^{-1}(U_{ij}).$$
Set $\mathbf{G} = [G_{ij}] \in \mathbb{R}^{n\times m}$ and estimate the Gaussian-copula correlation matrix as $\widehat{\mathbf{R}} \leftarrow \mathrm{Cor}(\mathbf{G})$.
For each synthetic null cell $i = 1, \ldots, n$, generate
$$\widetilde{\mathbf{G}}_i = (\widetilde{G}_{i1}, \ldots, \widetilde{G}_{im})^{\mathsf{T}} \overset{\text{i.i.d.}}{\sim} N_m(\mathbf{0}, \widehat{\mathbf{R}}), \qquad \widetilde{\mathbf{Y}}_i \leftarrow \left(\widehat{F}_1^{-1}\{\Phi(\widetilde{G}_{i1})\}, \ldots, \widehat{F}_m^{-1}\{\Phi(\widetilde{G}_{im})\}\right)^{\mathsf{T}}.$$
Set $\widetilde{\mathbf{Y}} = [\widetilde{\mathbf{Y}}_1, \ldots, \widetilde{\mathbf{Y}}_n]^{\mathsf{T}} \in \mathbb{N}_{\geq 0}^{n\times m}$, representing one hypothetical homogeneous cell type. The synthetic null cells are independently generated and do not correspond one-to-one to the target cells.

2. **Cluster the target and synthetic null cells.**
For $\mathbf{D} \in \{\mathbf{Y}, \widetilde{\mathbf{Y}}\}$, apply
$$\begin{aligned}
\mathbf{D}^{(1)} &\leftarrow \mathrm{LogNormalize}(\mathbf{D}, \text{scale factor} = 10{,}000),\\
\mathcal{H} &\leftarrow \mathrm{FindVariableFeatures}(\mathbf{D}^{(1)}, n_{\mathrm{HVG}} = 2{,}000),\\
\mathbf{D}^{(2)} &\leftarrow \mathrm{ScaleData}(\mathbf{D}^{(1)}_{\mathcal{H}}),\\
\mathbf{P}(\mathbf{D}) &\leftarrow \mathrm{PCA}(\mathbf{D}^{(2)}),\\
\mathbf{A}(\mathbf{D}) &\leftarrow \mathrm{KNN}(\mathbf{P}(\mathbf{D})_{\cdot,1:30}, k = 20),\\
\mathbf{Z}^{\mathrm{clust}}(\mathbf{D}) &\leftarrow \mathrm{Louvain}(\mathbf{A}(\mathbf{D}), \text{resolution}).
\end{aligned}$$
Adjust the resolution from 0.5 until two clusters are obtained, and set
$$\widehat{\mathbf{Z}} = (\widehat{Z}_1, \ldots, \widehat{Z}_n)^{\mathsf{T}} \leftarrow \mathbf{Z}^{\mathrm{clust}}(\mathbf{Y}), \qquad \widetilde{\mathbf{Z}} = (\widetilde{Z}_1, \ldots, \widetilde{Z}_n)^{\mathsf{T}} \leftarrow \mathbf{Z}^{\mathrm{clust}}(\widetilde{\mathbf{Y}}).$$

3. **Perform differential expression analysis.**
For a DE test $\mathcal{T}_{\mathrm{DE}} \in \{\text{Wilcoxon (default)}, \text{t-test}, \text{NB-GLM}, \text{LR}, \text{bimod}\}$, where $\mathcal{T}_{\mathrm{DE}}(\cdot,\cdot)$ returns a gene-level $P$ value.
For each gene $j = 1, \ldots, m$, compute the target and null $P$ values and their corresponding DE scores:
$$\begin{aligned}
P_j &\leftarrow \mathcal{T}_{\mathrm{DE}}(\mathbf{Y}_{\cdot j}, \widehat{\mathbf{Z}}), \quad \widetilde{P}_j \leftarrow \mathcal{T}_{\mathrm{DE}}(\widetilde{\mathbf{Y}}_{\cdot j}, \widetilde{\mathbf{Z}}),\\
S_j &\leftarrow -\log_{10}(P_j), \quad \widetilde{S}_j \leftarrow -\log_{10}(\widetilde{P}_j).
\end{aligned}$$

4. **Construct contrast scores and control the FDR.**
For each gene $j = 1, \ldots, m$, compute the contrast score
$$C_j \leftarrow S_j - \widetilde{S}_j.$$
Apply a one-sided Yuen trimmed-mean test:
$$p_{\mathrm{Yuen}} \leftarrow \mathrm{YuenTest}\left(\{C_j\}_{j=1}^{m}; \text{trim} = 0.1, \text{alternative} = \text{greater}\right).$$
**If** $p_{\mathrm{Yuen}} < 0.001$**:**
Fit the robust linear model
$$(\widehat{a}, \widehat{b}) \leftarrow \mathrm{RobustLinearModel}\left(\{(\widetilde{S}_j, S_j)\}_{j=1}^{m}\right), \qquad \widetilde{S}_j^{\mathrm{adj}} \leftarrow \widehat{a} + \widehat{b}\widetilde{S}_j,$$
and recompute $C_j \leftarrow S_j - \widetilde{S}_j^{\mathrm{adj}}$ for $j = 1, \ldots, m$.
**Otherwise:**
Retain $C_j = S_j - \widetilde{S}_j$ for $j = 1, \ldots, m$.
Let $\mathcal{C}$ denote the set of non-zero contrast-score values and select
$$T \leftarrow \min\left\{t \in \mathcal{C} : \frac{\mathrm{card}\{j : C_j \leq -t\} + 1}{\mathrm{card}\{j : C_j \geq t\} \vee 1} \leq q\right\}.$$

**Output:** A ranked list of reliable cell-type marker genes $\mathcal{D} = \{j : C_j \geq T\}$. If no eligible $T$ exists, return $\mathcal{D} = \varnothing$.

Algorithm S2: **ClusterDE-Core workflow for spatially resolved transcriptomics data**

**Input:** Spot-by-gene count matrix $\mathbf{Y} = [Y_{ij}] \in \mathbb{N}_{\geq 0}^{n \times m}$; spatial locations $\mathbf{X} = [X_1, \ldots, X_n]^\mathsf{T}$, where $X_i = (x_{i1}, x_{i2})^\mathsf{T}$; DE test $\mathcal{T}_{\text{DE}}$; target FDR level $q$.

1. **Generate synthetic null data.**
   For each gene $j = 1, \ldots, m$, fit
   $$Y_{ij} \overset{\text{ind.}}{\sim} \text{NB}(\mu_{ij}, \sigma_j), \qquad \log(\mu_{ij}) = \alpha_j + f_j^{\text{GP}}(x_{i1}, x_{i2}, 4),$$
   and estimate
   $$(\widehat{\alpha}_j, \widehat{f}_j^{\text{GP}}, \widehat{\sigma}_j) \leftarrow \text{FitSpatialNB}(\mathbf{Y}_{\cdot j}, \mathbf{X}), \qquad \widehat{F}_j \leftarrow F_{\text{NB}}(\,\cdot\,; \widehat{\mu}_{ij}, \widehat{\sigma}_j),$$
   where $\log(\widehat{\mu}_{ij}) = \widehat{\alpha}_j + \widehat{f}_j^{\text{GP}}(x_{i1}, x_{i2}, 4)$.
   For $i = 1, \ldots, n$ and $j = 1, \ldots, m$, compute
   $$V_{ij} \overset{\text{i.i.d.}}{\sim} \text{Uniform}(0, 1), \quad U_{ij} \leftarrow V_{ij}\widehat{F}_j(Y_{ij}) + (1 - V_{ij})\widehat{F}_j(Y_{ij} + 1), \quad G_{ij} \leftarrow \Phi^{-1}(U_{ij}).$$
   Set $\mathbf{G} = [G_{ij}] \in \mathbb{R}^{n \times m}$ and estimate $\widehat{\mathbf{R}} \leftarrow \text{Cor}(\mathbf{G})$.
   For each synthetic null spot $i = 1, \ldots, n$, generate
   $$\widetilde{\mathbf{G}}_i = (\widetilde{G}_{i1}, \ldots, \widetilde{G}_{im})^\mathsf{T} \overset{\text{i.i.d.}}{\sim} N_m(\mathbf{0}, \widehat{\mathbf{R}}), \qquad \widetilde{\mathbf{Y}}_i \leftarrow \left(\widehat{F}_1^{-1}\{\Phi(\widetilde{G}_{i1})\}, \ldots, \widehat{F}_m^{-1}\{\Phi(\widetilde{G}_{im})\}\right)^\mathsf{T}.$$
   Set $\widetilde{\mathbf{Y}} = [\widetilde{\mathbf{Y}}_1, \ldots, \widetilde{\mathbf{Y}}_n]^\mathsf{T}$, representing one hypothetical spatial domain with smooth spatial expression variation but no abrupt domain boundaries.

2. **Cluster the target and synthetic null spots.**
   For $\mathbf{D} \in \{\mathbf{Y}, \widetilde{\mathbf{Y}}\}$, apply
   $$\mathbf{P}(\mathbf{D}) \leftarrow \text{SpatialPreprocess}(\mathbf{D}, n_{\text{PC}} = 15, \text{log normalization}),$$
   $$\mathbf{Z}^{\text{clust}}(\mathbf{D}) \leftarrow \text{BayesSpace}(\mathbf{P}(\mathbf{D}), \mathbf{X}, q = 2, d = 7, \gamma = 2).$$
   Set
   $$\widehat{\mathbf{Z}} = (\widehat{Z}_1, \ldots, \widehat{Z}_n)^\mathsf{T} \leftarrow \mathbf{Z}^{\text{clust}}(\mathbf{Y}), \qquad \widetilde{\mathbf{Z}} = (\widetilde{Z}_1, \ldots, \widetilde{Z}_n)^\mathsf{T} \leftarrow \mathbf{Z}^{\text{clust}}(\widetilde{\mathbf{Y}}).$$

3. **Perform differential expression analysis.**
   For a DE test $\mathcal{T}_{\text{DE}} \in \{\text{Wilcoxon}, \text{t-test}\}$, where $\mathcal{T}_{\text{DE}}(\cdot, \cdot)$ returns a gene-level $P$ value.
   For each gene $j = 1, \ldots, m$, compute
   $$P_j \leftarrow \mathcal{T}_{\text{DE}}(\mathbf{Y}_{\cdot j}, \widehat{\mathbf{Z}}), \quad \widetilde{P}_j \leftarrow \mathcal{T}_{\text{DE}}(\widetilde{\mathbf{Y}}_{\cdot j}, \widetilde{\mathbf{Z}}),$$
   $$S_j \leftarrow -\log_{10}(P_j), \quad \widetilde{S}_j \leftarrow -\log_{10}(\widetilde{P}_j).$$

4. **Construct contrast scores and control the FDR.**
   For each gene $j = 1, \ldots, m$, compute $C_j \leftarrow S_j - \widetilde{S}_j$.
   Apply a one-sided Yuen trimmed-mean test:
   $$p_{\text{Yuen}} \leftarrow \text{YuenTest}\left(\{C_j\}_{j=1}^m; \text{trim} = 0.1, \text{alternative} = \text{greater}\right).$$
   **If** $p_{\text{Yuen}} < 0.001$**:**
   Fit the robust linear model
   $$(\widehat{a}, \widehat{b}) \leftarrow \text{RobustLinearModel}\left(\{(\widetilde{S}_j, S_j)\}_{j=1}^m\right), \qquad \widetilde{S}_j^{\text{adj}} \leftarrow \widehat{a} + \widehat{b}\widetilde{S}_j,$$
   and recompute $C_j \leftarrow S_j - \widetilde{S}_j^{\text{adj}}$ for $j = 1, \ldots, m$.
   **Otherwise:**
   Retain $C_j = S_j - \widetilde{S}_j$ for $j = 1, \ldots, m$.
   Let $\mathcal{C}$ denote the set of non-zero contrast-score values and select
   $$T \leftarrow \min\left\{t \in \mathcal{C} : \frac{\text{card}\{j : C_j \leq -t\} + 1}{\text{card}\{j : C_j \geq t\} \vee 1} \leq q\right\}.$$

**Output:** A ranked list of reliable spatial-domain marker genes $\mathcal{D} = \{j : C_j \geq T\}$. If no eligible $T$ exists, return $\mathcal{D} = \varnothing$.

Algorithm S3: **ClusterDE workflow for single-cell multiome data**

**Input:** Paired RNA count matrix $\mathbf{Y}^{\mathrm{RNA}} \in \mathbb{N}_{\geq 0}^{n \times m_{\mathrm{RNA}}}$; ATAC count matrix $\mathbf{Y}^{\mathrm{ATAC}} \in \mathbb{N}_{\geq 0}^{n \times m_{\mathrm{ATAC}}}$; RNA-derived cluster labels $\widehat{\mathbf{Z}} = (\widehat{Z}_1, \ldots, \widehat{Z}_n)^{\top}$; number of synthetic null replicates $B$; split threshold $\alpha_{\mathrm{split}}$; marker FDR $q_{\mathrm{marker}}$; stability threshold $t_S$.

1. **Evaluate the RNA-defined cluster split.**
   Following Step 1 of Algorithm S1, fit the NB–Gaussian-copula null model to $\mathbf{Y}^{\mathrm{RNA}}$ and generate $\widetilde{\mathbf{Y}}^{\mathrm{RNA},(1)}, \ldots, \widetilde{\mathbf{Y}}^{\mathrm{RNA},(B)}$.
   For each synthetic null replicate $b = 1, \ldots, B$, follow the clustering operations in Step 2 of Algorithm S1 to obtain the null representation (from PCA) $\widetilde{\mathbf{P}}^{\mathrm{RNA},(b)}$ and two null clusters $\widetilde{\mathbf{Z}}^{\mathrm{RNA},(b)}$, while retaining the original target labels $\widehat{\mathbf{Z}}$. Let $\mathbf{P}^{\mathrm{RNA}}$ denote the RNA representation used to define the target clusters.
   Compute
   $$W_{\mathrm{obs}} \leftarrow \mathrm{Ward}(\mathbf{P}^{\mathrm{RNA}}, \widehat{\mathbf{Z}}), \qquad W_b \leftarrow \mathrm{Ward}(\widetilde{\mathbf{P}}^{\mathrm{RNA},(b)}, \widetilde{\mathbf{Z}}^{\mathrm{RNA},(b)}),$$
   $$p_{\mathrm{split}} \leftarrow \frac{1}{B} \sum_{b=1}^{B} \mathbb{I}(W_b \geq W_{\mathrm{obs}}).$$
   **If** $p_{\mathrm{split}} \geq \alpha_{\mathrm{split}}$**:** merge the two clusters and stop.
   **Otherwise:** retain the split and proceed to marker identification.
2. **Identify stable RNA markers.**
   For a DE test $\mathcal{T}_{\mathrm{DE}}^{\mathrm{RNA}} \in \{\text{Wilcoxon (default)}, \text{t-test}, \text{NB-GLM}, \text{LR}, \text{bimod}\}$, where $\mathcal{T}_{\mathrm{DE}}^{\mathrm{RNA}}(\cdot, \cdot)$ returns a gene-level $P$ value, apply Steps 3–4 of Algorithm S1 to each target–null pair
   $$\left(\mathbf{Y}^{\mathrm{RNA}}, \widehat{\mathbf{Z}}\right) \quad \text{and} \quad \left(\widetilde{\mathbf{Y}}^{\mathrm{RNA},(b)}, \widetilde{\mathbf{Z}}^{\mathrm{RNA},(b)}\right),$$
   and obtain the selected gene set $\mathcal{G}_b^{\mathrm{RNA}}$ at FDR $q_{\mathrm{marker}}$.
   For each gene $j = 1, \ldots, m_{\mathrm{RNA}}$, compute
   $$s_j^{\mathrm{RNA}} \leftarrow \frac{1}{B} \sum_{b=1}^{B} \mathbb{I}(j \in \mathcal{G}_b^{\mathrm{RNA}}), \qquad \mathcal{G}_{\mathrm{stable}}^{\mathrm{RNA}} \leftarrow \{j : s_j^{\mathrm{RNA}} \geq t_S\}.$$
3. **Identify stable ATAC markers for the retained split.**
   Retain the same cells and RNA-derived cluster labels $\widehat{\mathbf{Z}}$. Following Step 1 of Algorithm S1, with genes replaced by peaks, fit the NB–Gaussian-copula null model to $\mathbf{Y}^{\mathrm{ATAC}}$ and generate $\widetilde{\mathbf{Y}}^{\mathrm{ATAC},(1)}, \ldots, \widetilde{\mathbf{Y}}^{\mathrm{ATAC},(B)}$.
   For each replicate $b = 1, \ldots, B$, obtain two null clusters using
   $$\widetilde{\mathbf{Y}}^{\mathrm{ATAC},(b)} \xrightarrow{\mathrm{TFIDF}} \widetilde{\mathbf{D}}^{(b)} \xrightarrow{\mathrm{TopFeatures}} \widetilde{\mathbf{D}}_{\mathcal{H}}^{(b)} \xrightarrow{\mathrm{SVD}} \widetilde{\mathbf{L}}^{(b)} \xrightarrow{\mathrm{KNN}(1:30, k=20)} \widetilde{\mathbf{A}}^{(b)} \xrightarrow{\mathrm{Louvain}} \widetilde{\mathbf{Z}}^{\mathrm{ATAC},(b)},$$
   adjusting the resolution from 0.5 until two clusters are obtained.
   For the Wilcoxon differential accessibility test $\mathcal{T}_{\mathrm{DA}}^{\mathrm{ATAC}}$, where $\mathcal{T}_{\mathrm{DA}}^{\mathrm{ATAC}}(\cdot, \cdot)$ returns a peak-level $P$ value, apply Steps 3–4 of Algorithm S1, with genes replaced by peaks, to each target–null pair and obtain $\mathcal{G}_b^{\mathrm{ATAC}}$ at FDR $q_{\mathrm{marker}}$.
   For each peak $k = 1, \ldots, m_{\mathrm{ATAC}}$, compute
   $$s_k^{\mathrm{ATAC}} \leftarrow \frac{1}{B} \sum_{b=1}^{B} \mathbb{I}(k \in \mathcal{G}_b^{\mathrm{ATAC}}), \qquad \mathcal{G}_{\mathrm{stable}}^{\mathrm{ATAC}} \leftarrow \{k : s_k^{\mathrm{ATAC}} \geq t_S\}.$$
4. **Evaluate modality-specific null quality.**
   Assess the synthetic null quality separately for the RNA and ATAC modalities before interpreting their stable markers.

**Output:** A retain-or-merge decision and, for a retained split, ranked lists of stable RNA markers $\mathcal{G}_{\mathrm{stable}}^{\mathrm{RNA}}$ and stable ATAC markers $\mathcal{G}_{\mathrm{stable}}^{\mathrm{ATAC}}$, ranked in decreasing order of $s_j^{\mathrm{RNA}}$ and $s_k^{\mathrm{ATAC}}$, respectively. If the split is not retained, return both marker sets as $\varnothing$.

Algorithm S4: **ClusterDE-Core workflow for bulk transcriptomic data**

**Input:** Sample-by-gene matrix $\mathbf{Y} = [Y_{ij}] \in \mathbb{R}^{n\times m}$; data type $b \in \{\text{microarray}, \text{RNA-seq}\}$; clustering method $g$; DE test $\mathcal{T}_{\text{DE}}$; target FDR level $q$.

1. **Generate synthetic null data.**
   **If $b$ = microarray:**
   Using preprocessed normalized expression data, estimate
   $$\widehat{\boldsymbol{\mu}} \leftarrow \frac{1}{n}\sum_{i=1}^{n}\mathbf{Y}_i, \qquad \widehat{\boldsymbol{\Sigma}} \leftarrow \frac{1}{n-1}\sum_{i=1}^{n}(\mathbf{Y}_i - \widehat{\boldsymbol{\mu}})(\mathbf{Y}_i - \widehat{\boldsymbol{\mu}})^\intercal.$$
   For $i = 1, \ldots, n$, generate $\widetilde{\mathbf{Y}}_i \overset{\text{i.i.d.}}{\sim} N_m(\widehat{\boldsymbol{\mu}}, \widehat{\boldsymbol{\Sigma}})$.
   **If $b$ = RNA-seq:**
   For each gene $j = 1, \ldots, m$, estimate
   $$(\widehat{\mu}_j, \widehat{\sigma}_j) \leftarrow \underset{\mu_j,\sigma_j}{\arg\max}\sum_{i=1}^{n}\log f_{\text{NB}}(Y_{ij}; \mu_j, \sigma_j), \qquad \widehat{F}_j \leftarrow F_{\text{NB}}(\,\cdot\,; \widehat{\mu}_j, \widehat{\sigma}_j).$$
   For $i = 1, \ldots, n$ and $j = 1, \ldots, m$, compute
   $$V_{ij} \overset{\text{i.i.d.}}{\sim} \text{Uniform}(0, 1), \quad U_{ij} \leftarrow V_{ij}\widehat{F}_j(Y_{ij}) + (1 - V_{ij})\widehat{F}_j(Y_{ij} + 1), \quad G_{ij} \leftarrow \Phi^{-1}(U_{ij}).$$
   Set $\mathbf{G} = [G_{ij}]$ and estimate $\widehat{\mathbf{R}} \leftarrow \text{Cor}(\mathbf{G})$.
   For each synthetic null sample $i = 1, \ldots, n$, generate
   $$\widetilde{\mathbf{G}}_i = (\widetilde{G}_{i1}, \ldots, \widetilde{G}_{im})^\intercal \overset{\text{i.i.d.}}{\sim} N_m(\mathbf{0}, \widehat{\mathbf{R}}), \qquad \widetilde{\mathbf{Y}}_i \leftarrow \left(\widehat{F}_1^{-1}\{\Phi(\widetilde{G}_{i1})\}, \ldots, \widehat{F}_m^{-1}\{\Phi(\widetilde{G}_{im})\}\right)^\intercal.$$
   In either case, set $\widetilde{\mathbf{Y}} = [\widetilde{\mathbf{Y}}_1, \ldots, \widetilde{\mathbf{Y}}_n]^\intercal$, representing one hypothetical homogeneous sample population.
2. **Cluster the target and synthetic null samples.**
   For $\mathbf{D} \in \{\mathbf{Y}, \widetilde{\mathbf{Y}}\}$, apply
   $$\mathbf{Z}^{\text{clust}}(\mathbf{D}) \leftarrow g(\mathbf{D}; \boldsymbol{\theta}_g), \qquad g \in \{\text{K-means}, \text{hierarchical clustering}\},$$
   where $\boldsymbol{\theta}_g$ contains the preprocessing, selected features, distance metric, linkage or initialization, and number of clusters used in the original study.
   Set
   $$\widehat{\mathbf{Z}} = (\widehat{Z}_1, \ldots, \widehat{Z}_n)^\intercal \leftarrow \mathbf{Z}^{\text{clust}}(\mathbf{Y}), \qquad \widetilde{\mathbf{Z}} = (\widetilde{Z}_1, \ldots, \widetilde{Z}_n)^\intercal \leftarrow \mathbf{Z}^{\text{clust}}(\widetilde{\mathbf{Y}}).$$
3. **Perform differential expression analysis.**
   For a DE test
   $$\mathcal{T}_{\text{DE}} \in \begin{cases}\{\text{limma}, \text{Wilcoxon}, \text{t-test}\}, & b = \text{microarray},\\ \{\text{DESeq2}, \text{edgeR}\}, & b = \text{RNA-seq},\end{cases}$$
   where $\mathcal{T}_{\text{DE}}(\cdot,\cdot)$ returns a gene-level $P$ value.
   For each gene $j = 1, \ldots, m$, compute
   $$P_j \leftarrow \mathcal{T}_{\text{DE}}(\mathbf{Y}_{\cdot j}, \widehat{\mathbf{Z}}), \quad \widetilde{P}_j \leftarrow \mathcal{T}_{\text{DE}}(\widetilde{\mathbf{Y}}_{\cdot j}, \widetilde{\mathbf{Z}}),$$
   $$S_j \leftarrow -\log_{10}(P_j), \quad \widetilde{S}_j \leftarrow -\log_{10}(\widetilde{P}_j).$$
4. **Construct contrast scores and control the FDR.**
   For each gene $j = 1, \ldots, m$, compute $C_j \leftarrow S_j - \widetilde{S}_j$.
   Apply a one-sided Yuen trimmed-mean test:
   $$p_{\text{Yuen}} \leftarrow \text{YuenTest}\left(\{C_j\}_{j=1}^{m}; \text{trim} = 0.1, \text{alternative} = \text{greater}\right).$$
   **If $p_{\text{Yuen}} < 0.001$:**
   Fit the robust linear model
   $$(\widehat{a}, \widehat{b}) \leftarrow \text{RobustLinearModel}\left(\{(\widetilde{S}_j, S_j)\}_{j=1}^{m}\right), \qquad \widetilde{S}_j^{\text{adj}} \leftarrow \widehat{a} + \widehat{b}\widetilde{S}_j,$$
   and recompute $C_j \leftarrow S_j - \widetilde{S}_j^{\text{adj}}$ for $j = 1, \ldots, m$.
   **Otherwise:**
   Retain $C_j = S_j - \widetilde{S}_j$ for $j = 1, \ldots, m$.
   Let $\mathcal{C}$ denote the set of non-zero contrast-score values and select
   $$T \leftarrow \min\left\{t \in \mathcal{C} : \frac{\text{card}\{j : C_j \le -t\} + 1}{\text{card}\{j : C_j \ge t\} \vee 1} \le q\right\}.$$

**Output:** A ranked list of reliable subgroup marker genes $\mathcal{D} = \{j : C_j \ge T\}$. If no eligible $T$ exists, return $\mathcal{D} = \varnothing$.

Algorithm S5: **ClusterDE-Core workflow for microbiome data**

**Input:** Sample-by-taxon abundance matrix $\mathbf{Y} = [Y_{ij}] \in \mathbb{N}_{\geq 0}^{n\times m}$; sequencing depths $d_1, \ldots, d_n$; clustering method $g$; differential-abundance test $\mathcal{T}_{\mathrm{DA}}$; target FDR level $q$.

1. **Generate synthetic null data.**
For each taxon $j = 1, \ldots, m$, fit
$$Y_{ij} \overset{\text{ind.}}{\sim} \mathrm{NB}(\mu_{ij}, \sigma_j), \qquad \log(\mu_{ij}) = \alpha_j + \beta_j \log(d_i),$$
and estimate
$$(\widehat{\alpha}_j, \widehat{\beta}_j, \widehat{\sigma}_j) \leftarrow \underset{\alpha_j,\beta_j,\sigma_j}{\arg\max} \sum_{i=1}^{n} \log f_{\mathrm{NB}}\left(Y_{ij}; \exp\{\alpha_j + \beta_j \log(d_i)\}, \sigma_j\right),$$
together with the fitted marginal CDF $\widehat{F}_j$.
For $i = 1, \ldots, n$ and $j = 1, \ldots, m$, compute
$$V_{ij} \overset{\text{i.i.d.}}{\sim} \mathrm{Uniform}(0,1), \quad U_{ij} \leftarrow V_{ij}\widehat{F}_j(Y_{ij}) + (1 - V_{ij})\widehat{F}_j(Y_{ij}+1), \quad G_{ij} \leftarrow \Phi^{-1}(U_{ij}).$$
Set $\mathbf{G} = [G_{ij}] \in \mathbb{R}^{n\times m}$ and estimate $\widehat{\mathbf{R}} \leftarrow \mathrm{Cor}(\mathbf{G})$.
For each synthetic null sample $i = 1, \ldots, n$, generate
$$\widetilde{\mathbf{G}}_i = (\widetilde{G}_{i1}, \ldots, \widetilde{G}_{im})^{\mathsf{T}} \overset{\text{i.i.d.}}{\sim} N_m(\mathbf{0}, \widehat{\mathbf{R}}), \qquad \widetilde{\mathbf{Y}}_i \leftarrow \left(\widehat{F}_1^{-1}\{\Phi(\widetilde{G}_{i1})\}, \ldots, \widehat{F}_m^{-1}\{\Phi(\widetilde{G}_{im})\}\right)^{\mathsf{T}},$$
using the observed sequencing depth $d_i$.
Set $\widetilde{\mathbf{Y}} = [\widetilde{\mathbf{Y}}_1, \ldots, \widetilde{\mathbf{Y}}_n]^{\mathsf{T}}$, representing one hypothetical homogeneous sample population.

2. **Cluster the target and synthetic null samples.**
For $\mathbf{D} \in \{\mathbf{Y}, \widetilde{\mathbf{Y}}\}$, apply
$$\mathbf{Z}^{\mathrm{clust}}(\mathbf{D}) \leftarrow g(\mathbf{D}; \boldsymbol{\theta}_g), \qquad g \in \{\text{K-means}, \text{hierarchical clustering}\},$$
where $\boldsymbol{\theta}_g$ contains the normalization, transformation, feature selection, distance metric, linkage or initialization, and number of clusters used in the original study.
Set
$$\widehat{\mathbf{Z}} = (\widehat{Z}_1, \ldots, \widehat{Z}_n)^{\mathsf{T}} \leftarrow \mathbf{Z}^{\mathrm{clust}}(\mathbf{Y}), \qquad \widetilde{\mathbf{Z}} = (\widetilde{Z}_1, \ldots, \widetilde{Z}_n)^{\mathsf{T}} \leftarrow \mathbf{Z}^{\mathrm{clust}}(\widetilde{\mathbf{Y}}).$$

3. **Perform differential-abundance analysis.**
For a differential-abundance test $\mathcal{T}_{\mathrm{DA}} \in \{\text{ANCOM-BC2}, \text{Wilcoxon}, \text{MaAsLin2}\}$, where $\mathcal{T}_{\mathrm{DA}}(\cdot,\cdot)$ returns a taxon-level $P$ value.
For each taxon $j = 1, \ldots, m$, compute
$$P_j \leftarrow \mathcal{T}_{\mathrm{DA}}(\mathbf{Y}_{\cdot j}, \widehat{\mathbf{Z}}), \quad \widetilde{P}_j \leftarrow \mathcal{T}_{\mathrm{DA}}(\widetilde{\mathbf{Y}}_{\cdot j}, \widetilde{\mathbf{Z}}),$$
$$S_j \leftarrow -\log_{10}(P_j), \quad \widetilde{S}_j \leftarrow -\log_{10}(\widetilde{P}_j).$$
Apply identical normalization, covariates, and model specifications to the target and synthetic null analyses.

4. **Construct contrast scores and control the FDR.**
For each taxon $j = 1, \ldots, m$, compute $C_j \leftarrow S_j - \widetilde{S}_j$.
Apply a one-sided Yuen trimmed-mean test:
$$p_{\mathrm{Yuen}} \leftarrow \mathrm{YuenTest}\left(\{C_j\}_{j=1}^{m}; \text{trim} = 0.1, \text{alternative} = \text{greater}\right).$$
**If $p_{\mathrm{Yuen}} < 0.001$:**
Fit the robust linear model
$$(\widehat{a}, \widehat{b}) \leftarrow \mathrm{RobustLinearModel}\left(\{(\widetilde{S}_j, S_j)\}_{j=1}^{m}\right), \qquad \widetilde{S}_j^{\mathrm{adj}} \leftarrow \widehat{a} + \widehat{b}\widetilde{S}_j,$$
and recompute $C_j \leftarrow S_j - \widetilde{S}_j^{\mathrm{adj}}$ for $j = 1, \ldots, m$.
**Otherwise:**
Retain $C_j = S_j - \widetilde{S}_j$ for $j = 1, \ldots, m$.
Let $\mathcal{C}$ denote the set of non-zero contrast-score values and select
$$T \leftarrow \min\left\{t \in \mathcal{C} : \frac{\mathrm{card}\{j : C_j \leq -t\} + 1}{\mathrm{card}\{j : C_j \geq t\} \vee 1} \leq q\right\}.$$

**Output:** A ranked list of reliable differentially abundant taxa $\mathcal{D} = \{j : C_j \geq T\}$. If no eligible $T$ exists, return $\mathcal{D} = \varnothing$.
